# Ranking species based on sensitivity to perturbations under non-equilibrium community dynamics

**DOI:** 10.1101/2022.07.23.501258

**Authors:** Lucas P. Medeiros, Stefano Allesina, Vasilis Dakos, George Sugihara, Serguei Saavedra

## Abstract

Managing ecological communities requires fast detection of species that are sensitive to perturbations. Yet, the focus on recovery to equilibrium has prevented us from assessing species responses to perturbations when abundances fluctuate over time. Here, we introduce two data-driven approaches (expected sensitivity and eigenvector rankings) based on the time-varying Jacobian matrix to rank species over time according to their sensitivity to perturbations on abundances. Using several population dynamics models, we demonstrate that we can infer these rankings from time-series data to predict the order of species sensitivities. We find that the most sensitive species are not always the ones with the most rapidly changing or lowest abundance, which are typical criteria used to monitor populations. Finally, using two empirical time series, we show that sensitive species tend to be harder to forecast. Our results suggest that incorporating information on species interactions can improve how we manage communities out of equilibrium.

## Introduction

Ecological communities—the collection of interacting populations in a given place and time—are subject to external perturbations, which are increasing in magnitude and frequency due to anthropogenic impacts (Barlow *et al*., 2018, Jackson *et al*., 2001). Indeed, strong and frequent perturbations can lead to species extinctions and, as a consequence, to the loss of critical ecosystem services (Cardinale *et al*., 2012, Levin & Lubchenco, 2008). In order to avoid the loss of biodiversity and ecosystem services under these circumstances, it is crucial to understand not only the response of the whole community to perturbations, but also the response of its constituent species. Individual species may vary in their sensitivity to perturbations—that is, how much their abundance change after a perturbation—and such sensitivity may be linked to their role in the community (Beauchesne *et al*., 2021, Dirzo *et al*., 2014, Estes *et al*., 2011). For instance, keystone species such as apex predators can be highly sensitive to perturbations and also crucial to maintain community functioning (Estes *et al*., 2011). Thus, detecting sensitive species has the potential to greatly improve management and conservation strategies for maintaining community functioning and avoiding biodiversity loss.

Traditional studies in theoretical population ecology have established several important measures of how single species respond to external perturbations (Caswell, 2000, Morris *et al*., 2002). Following these theoretical developments, indicators such as species abundance or rate of decline are routinely used to characterize the behavior of populations and determine extinction risks, serving as crucial management and conservation tools (Mace *et al*., 2008). More recently, several studies have incorporated information on species interactions to further explore how individual species respond to perturbations (Arnoldi *et al*., 2018, Beauchesne *et al*., 2021, Ives *et al*., 1999, Medeiros *et al*., 2021, Saavedra *et al*., 2011, Weinans *et al*., 2019) and, in turn, how individual species can inform us about whole-community changes (i.e., best-indicator or sensor species) (Aparicio *et al*., 2021, Dakos, 2018, Ghadami *et al*., 2020, Lever *et al*., 2020, Patterson *et al*., 2021). These studies often rely on the assumption of a population dynamics model under a stable equilibrium to which the community returns after a small perturbation on abundances. Under this assumption, information on the Jacobian matrix—the matrix containing the local effects of each species on the growth rate of other species and itself (Song & Saavedra, 2021)—can be used to partition the recovery rate of the community into its constituent species (Arnoldi *et al*., 2018, Ives *et al*., 1999, Medeiros *et al*., 2021). A community slightly displaced from equilibrium will asymptotically return along the direction spanned by the leading eigenvector of the Jacobian matrix, that is, the eigenvector associated with the leading (i.e., largest) eigenvalue (Dakos, 2018, Patterson *et al*., 2021, Strogatz, 2018). Thus, different species may show distinct recovery rates after a perturbation depending on the direction of the leading eigenvector (Dakos, 2018, Ghadami *et al*., 2020, Patterson *et al*., 2021, Weinans *et al*., 2019), although this effect may not be instantaneous due to long transients (Hastings *et al*., 2018, Rinaldi & Scheffer, 2000). Nevertheless, these ideas cannot be directly applied to communities for which abundances exhibit non-equilibrium fluctuations over time such as many natural communities with cyclic or chaotic dynamics (Ben-incà *et al*., 2015, 2009, Clark & Luis, 2020, Krebs *et al*., 1995, Sugihara, 1994, Ushio *et al*., 2018). Moreover, from a practical point of view, it can be unfeasible to monitor how species respond to perturbations using parameterized models given the large amounts of empirical data required to test model assumptions and infer parameter values (Bartomeus *et al*., 2021, Bender *et al*., 1984).

The above limitations raise the question of whether we can measure species responses to perturbations in communities for which dynamics are not at equilibrium (Hastings *et al*., 2018, Sugihara, 1994). To address this problem, recent methodologies have focused on extracting information directly from abundance time series and measuring how non-equilibrium communities respond to perturbations (Cenci & Saavedra, 2019, Ushio *et al*., 2018). Using a data-driven method known as the S-map to reconstruct the time-varying Jacobian matrix (Cenci *et al*., 2019, Deyle *et al*., 2016, Sugihara, 1994), recent studies have investigated how communities respond to perturbations on abundances (Ushio *et al*., 2018) and to perturbations on the laws that govern community dynamics (Cenci & Saavedra, 2019). Regarding perturbations on abundances, it has been suggested that the leading eigenvalue of the Jacobian matrix can be used to quantify how communities respond to small perturbations at any given time (Ushio *et al*., 2018). Differently from a recovery rate in a community with a stable equilibrium as described in the previous paragraph, under non-equilibrium dynamics the leading eigenvalue approximates the local growth rate of small perturbations along a given direction (Eckmann & Ruelle, 1985, Mease *et al*., 2003, Vallejo *et al*., 2017). Thus, in contrast to a community at equilibrium with a constant capacity to recover from perturbations, a community under non-equilibrium dynamics has a response to perturbations that depends on how species abundances change over time (i.e., state-dependent) (Cenci & Saavedra, 2019). In particular, the state of a community may determine its response to perturbations not only through the local species effects on each other (i.e., Jacobian matrix) but also through the local time scale of the dynamics (e.g., perturbation effects may take a long time to appear under a long transient) (Hastings *et al*., 2018, Rinaldi & Scheffer, 2000). The question that remains to be answered is whether we can decompose this community-level information to monitor the time-varying sensitivity of each species and whether this can complement traditional demographic indicators that do not use community-level information.

Here, we develop two complementary approaches based on dynamical systems theory and nonlinear time series analysis to rank species over time under non-equilibrium dynamics according to their sensitivity to small perturbations on abundances. By doing so, we provide a data-driven framework to detect which species in a community are the most and least sensitive to perturbations at any given time. Our ranking approaches consist of an analytical measure of the expected sensitivity of each species and an alignment measure of each species with the leading eigenvector of the time-varying Jacobian matrix. We test both approaches by performing perturbation analyses using five synthetic time series generated from population dynamics models. We show that we can accurately rank species sensitivities, especially using the expected sensitivity approach. However, the eigenvector approach performs better when information on perturbations is greatly misspecified for the computation of expected sensitivities. Importantly, we show that both ap-proaches remain accurate when inferring the Jacobian matrix directly from the time series with the S-map. Finally, we apply both approaches to two empirical non-equilibrium time series and show that species that are more sensitive to perturbations at a given time are harder to forecast, especially when the local growth rate of perturbations is high.

### Quantifying species sensitivities to perturbations

To quantify species sensitivities to perturbations, we assume that species abundances in a community with *S* species change through time according to a generic function: 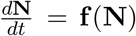, where **f** (**f**: ℝ^*S*^ → ℝ^*S*^) is an unknown nonlinear model and **N** = [*N*_1_, …, *N*_*S*_]^⊤^ is the vector of species abundances (Cenci *et al*., 2020, Cenci & Saavedra, 2019). At any given time, the community can be affected by a pulse perturbation **p** = [*p*_1_, …, *p*_*S*_]^⊤^ that changes **N** into **Ñ**(i.e., **Ñ**= **N** + **p**) (Bender *et al*., 1984). The vector **Ñ** would then change in time according to **f**. Following similar definitions in ecology, we conceptually define sensitivity as the amount of change in species abundances following a perturbation (Dakos, 2018, Domínguez-García *et al*., 2019). Mathematically, we define the sensitivity of species *i* to a specific perturbation **p** from time *t* to *t* + *k* as the squared difference between its perturbed and unperturbed abundance at time *t* + *k* in relation to the initial squared difference caused by the perturbation at time *t*:

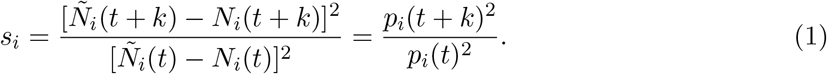

Therefore, *s*_*i*_ quantifies the distance between perturbed and unperturbed states over time, similarly to measures of sensitivity to initial conditions in nonlinear dynamics (Eckmann & Ruelle, 1985, Strogatz, 2018, Vallejo *et al*., 2017). However, *s*_*i*_ is completely dependent on **p**. Because we typically have no prior information about the direction and magnitude of external perturbations in natural communities, here we focus on a collection of randomly perturbed abundances. Thus, we define the sensitivity of species *i* from time *t* to *t* + *k* as the average squared difference between a set of *n* randomly perturbed abundances and its unperturbed abundance at time *t* + *k* in relation to the initial average squared difference at time *t*:

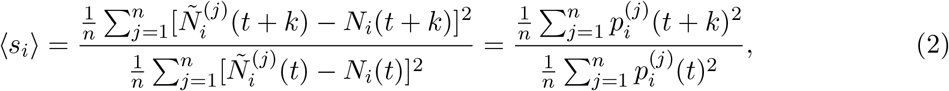

where 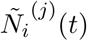 is the *j*th perturbed abundance of species *i* at time *t*. Note that we use ⟨*s_i_*⟩ as a notation for the ratio of the mean squared deviations. The denominator in equation (2) controls for the initial displacement of species abundances, but can be ignored if all species are under the same perturbation regime (*SI Section 4*). Also note that ⟨*s*_*i*_⟩ ≥ 0 and is not bounded because the numerator may be arbitrarily large.

Under non-equilibrium dynamics, the identity of the most and least sensitive species can change over time. We illustrate this statement using the following 3-species food chain model that exhibits chaotic dynamics (Hastings & Powell, 1991) (parameter values given in *SI Section 3*):

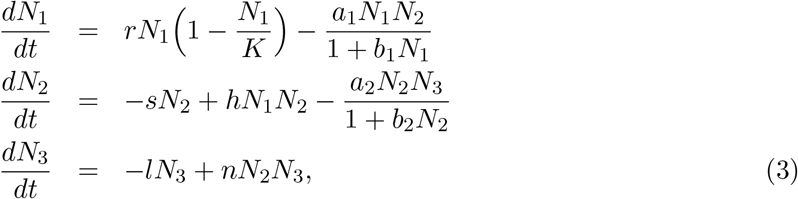

where *N*_1_, *N*_2_, and *N*_3_ are the abundances of the primary producer, primary consumer, and secondary consumer, respectively. To study species sensitivities under this model, we numerically integrate equation (3) producing abundance time series (Fig. 1A, B) that can be visualized as an attractor in the state space of abundances (Fig. 1C, D). Then, we perform a small arbitrary pulse perturbation **p** that increases the abundance of all species at time *t* (orange vertical line in Fig. 1A, B) and compute species sensitivities to it (*s*_*i*_) after *k* = 1 time step (red vertical line in Fig. 1A, B). Note that we use *k* = 1 as an example here, but explore the effects of changing this time step in multiple analyses. Fig. 1A, B shows that even under the same perturbation **p**, species exhibit drastically different sensitivities depending on when the perturbation occurs. That is, the species that has the largest sensitivity to this particular perturbation can change from the primary (species 2 in Fig. 1A) to the secondary consumer (species 3 in Fig. 1B) after just a few time steps. Next, we extend our illustration and consider how multiple randomly perturbed abundances (**Ñ**(*t*), orange points in Fig. 1C, D) change after one time step (**Ñ**(*t*+1), red points in Fig. 1C, D) by computing species sensitivities (⟨*s*_*i*_⟩). Fig. 1C, D confirms that the most sensitive species changes from the primary (species 2 in Fig. 1C) to the secondary consumer (species 3 in Fig. 1D) under random perturbations. Therefore, the problem we aim to solve in this study is how to predict the order of the ⟨*s*_*i*_⟩ values of all species in a community at any given time. Clearly, in natural communities we cannot produce multiple random perturbations to compare the responses of different species in perturbed and unperturbed communities. Therefore, in what follows, we provide a rationale for using the Jacobian matrix at time *t* to predict the order of ⟨*s*_*i*_⟩ values.

**Figure 1.**
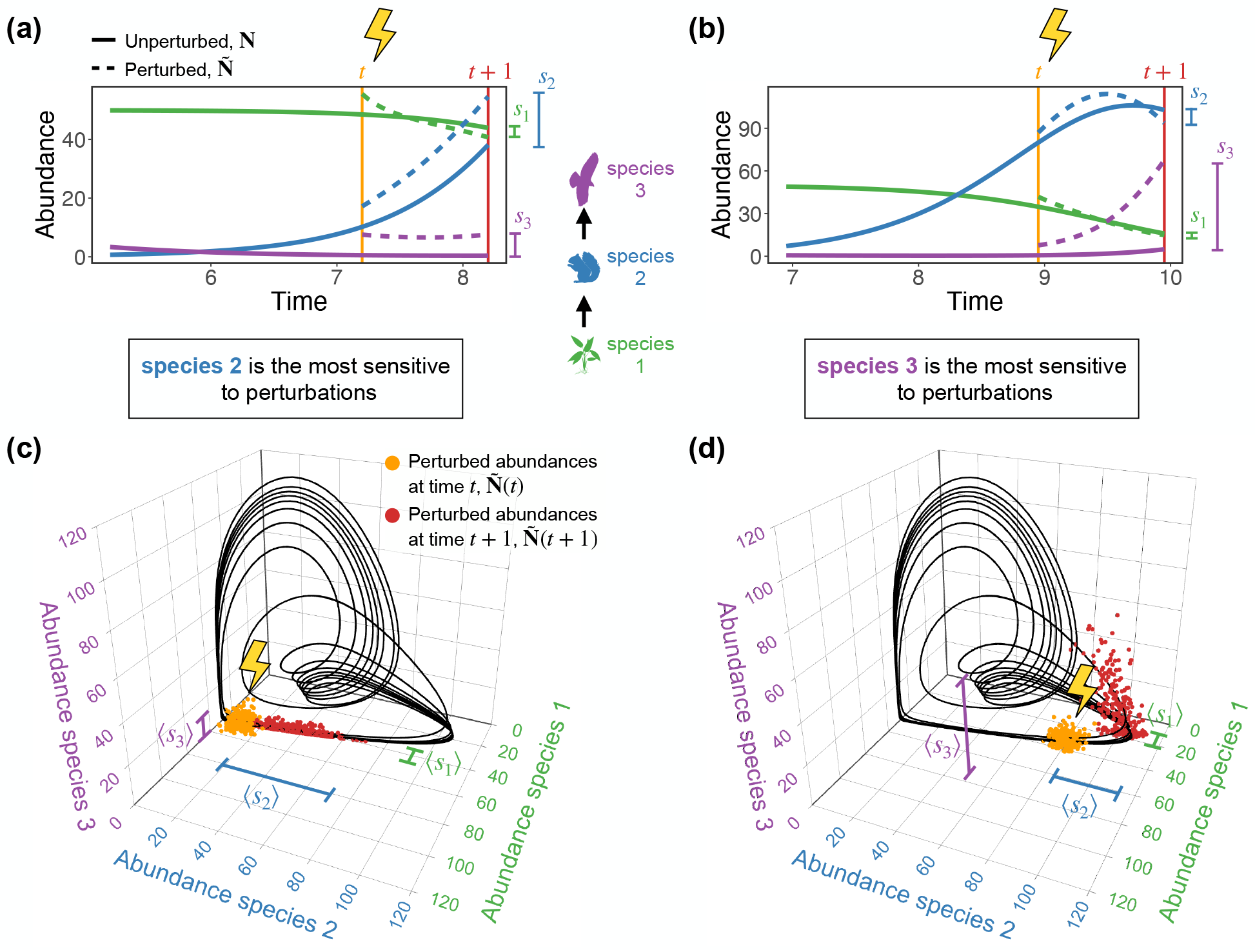
Identity of most sensitive species to perturbations changes through time under non-equilibrium dynamics. (a) and (b) Abundance time series generated from a 3-species chaotic food chain model (equation (3)) showing the effect of a pulse perturbation **p** = [7, 7, 7]^⊤^ that increases all abundances at different times *t*. Whereas species 2 (primary consumer, blue) shows the highest sensitivity *s*_*i*_ to **p** (i.e., the largest squared difference between perturbed and unperturbed abundance at *t*+1) in (a), species 3 (secondary consumer, purple) shows the highest *s*_*i*_ to **p** just a few time steps ahead in (b). (c) and (d) Chaotic attractor of the food chain model (black) with multiple perturbed abundances around **N**(*t*) (**Ñ**(*t*), orange points) at different times *t*. The red points show these perturbed abundances after one time step (**Ñ**(*t*+1)). We can measure the sensitivity of species *i* to random perturbations (⟨*s*_*i*_⟩) by computing the average squared difference between its set of perturbed abundances (*Ñ*_*i*_(*t* + 1)) and its unperturbed abundance (*N*_*i*_(*t* + 1)) at *t* + 1. Note that this sensitivity measure is normalized by the average squared difference between *Ñ*_*i*_(*t*) and *N*_*i*_(*t*) at time *t* (equation (2)). Whereas species 2 shows the highest ⟨*s*_*i*_⟩ in (c), species 3 shows the highest ⟨*s*_*i*_⟩ in (d).

### Ranking species sensitivities to perturbations

Without loss of generality, we can write the linearized dynamics of a small perturbation on abundances as 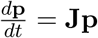, where **J** is the Jacobian matrix of **f** evaluated at **N** (*SI Section 1*) (Boyce *et al*., 2017, Eckmann & Ruelle, 1985, Mease *et al*., 2003, Strogatz, 2018). Following results from dynamical systems theory (Arnoldi *et al*., 2018, Boyce *et al*., 2017, Strogatz, 2018), we propose two complementary approaches to rank species according to their sensitivity to perturbations (Box 1 and 2). These approaches are based on the assumption that if **J** is nearly constant from time *t* to *t* + *k*, the solution **p**(*t* + *k*) of the linearized dynamics provides a good approximation to how perturbed abundances change over this time period. Thus, in addition to the challenge of approximating a nonlinear system (i.e., **f** (**N**)) by its linearized dynamics, here we explore the extent to which the linearized dynamics informs us about species sensitivities under non-equilibrium dynamics (i.e., when **J** is state-dependent). Importantly, note that we use information on species effects on each other contained in the Jacobian matrix (i.e., community-level information) to measure how individual species respond to perturbations.

#### Box 1

Expected sensitivity ranking

##### Rationale

This approach is based on analytically computing an expected value for the sensitivity of species *i* to perturbations (𝔼(*s*_*i*_)) using the solution **p**(*t* + *k*) = *ϵ*^**J***k*^**p**(*t*) of the linearized dynamics (*SI Section 2*) (Boyce *et al*., 2017). Note that, for sufficiently small perturbations under equilibrium dynamics, this solution is exact because **J** is constant when evaluated at an equilibrium point (**N**^*∗*^ for which **f** (**N**^*∗*^) = **0**). By assuming that **p**(*t*) follows a distribution with mean zero, we can obtain 𝔼(*s*_*i*_) at time *t* from the covariance matrix of **p**(*t* + *k*): **Σ**_*t*+*k*_ = *e*^**J***k*^**Σ**_*t*_(*e*^**J***k*^)^⊤^, where **Σ**_*t*_ is the covariance matrix of **p**(*t*). That is, the distribution of perturbed abundances (**Ñ**) described by **Σ**_*t*_ will approximate **Σ**_*t*+*k*_ after *k* time steps (Fig. 2A). Then, we can compute the expected sensitivity of species *i* as: 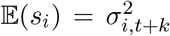, where 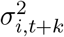 is the *i*th diagonal element of **Σ**_*t*+*k*_ (i.e., the variance of *p*_*i*_(*t* + *k*)). We define the order of 𝔼(*s*_*i*_) values across species as the expected sensitivity ranking and use it to predict the order of species sensitivities to perturbations (⟨*s*_*i*_⟩; Fig. 2A).

##### Application

Three ingredients are required to apply the expected sensitivity ranking. First, we need the Jacobian matrix of the community (**J**) evaluated using the abundances (**N**) at time *t*. This matrix can be computed directly from a parameterized population dynamics model (*SI Section 1*) or, as we focus here, inferred from the abundance time series without assuming a specific model (*SI Section 5*). Second, we need to define an initial covariance matrix of perturbations at time *t* (**Σ**_*t*_). Without any knowledge of perturbations, we suggest an uninformative approach by setting **Σ**_*t*_ = **I**, where **I** is the identity matrix (i.e., perturbations to each species are independent from each other). Finally, we need to specify the time for which perturbations evolve (*k*). Because information on the local time scale of the dynamics can be challenging to obtain, we suggest setting *k* equal to a small constant (e.g., *k* = 1). Alternatively, it is possible to set *k* to be inversely proportional to the local rate of change calculated from the time series (*SI Section 4*). In conclusion, although this approach has the advantage of using information on the entire Jacobian matrix to compute 𝔼(*s*_*i*_), it has the disadvantage of requiring additional information that may be hard to obtain in natural communities (**Σ**_*t*_ and *k*).

**Figure 2.**
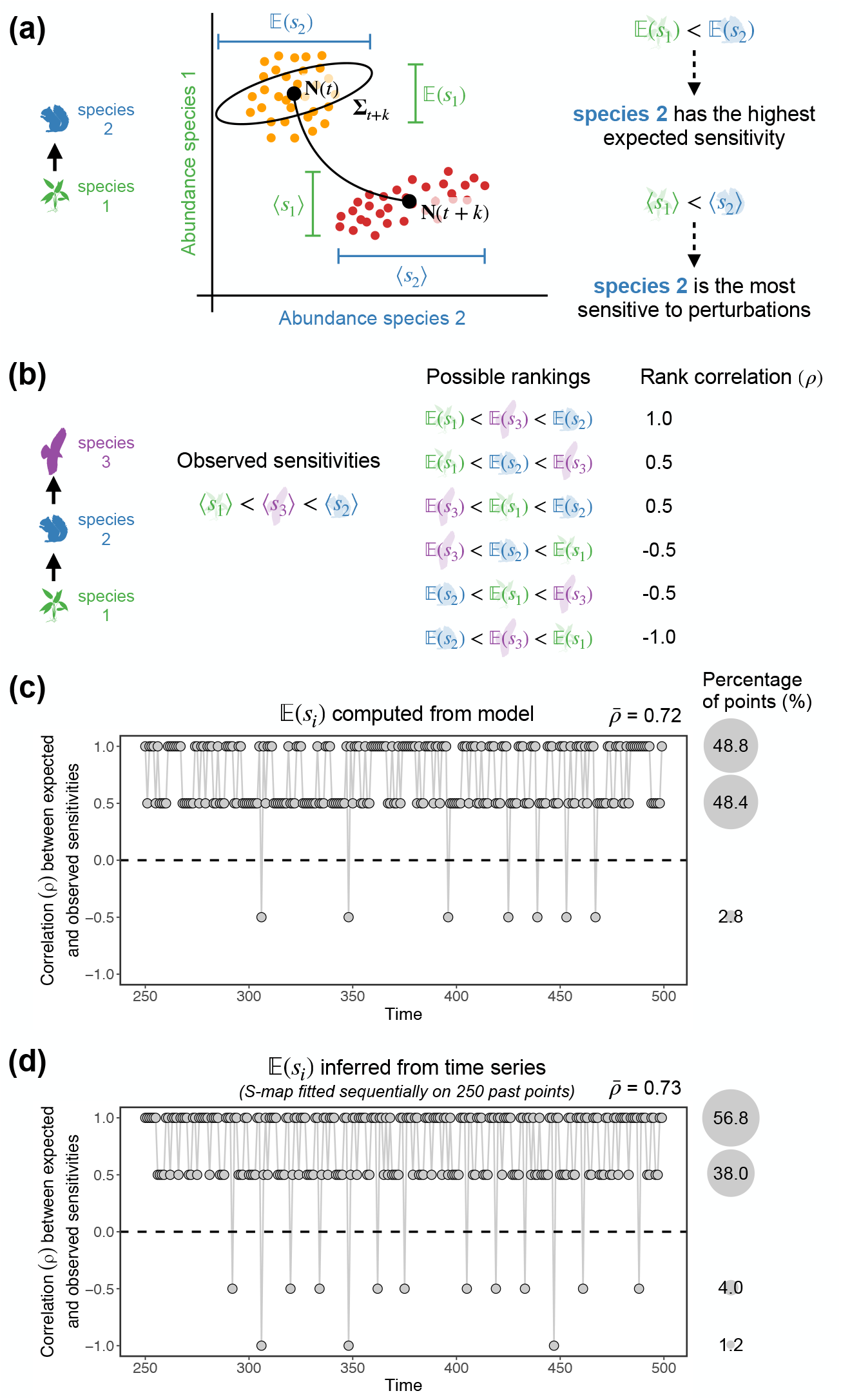
Ranking species sensitivities to perturbations. (a) Illustration with two species showing the expansion of perturbed abundances (**Ñ**(*t*), orange points) after *k* time steps (**Ñ**(*t* + *k*), red points). The expected sensitivity of species *i* to perturbations (𝔼(*s*_*i*_)) can be computed as the corresponding variance of the predicted distribution of perturbations (i.e., *i*th diagonal element of covariance matrix **Σ**_*t*+*k*_ depicted in gray). Note that **Σ**_*t*+*k*_ is shown at time *t* as it is computed using only information at that time point. We propose that the order of 𝔼(*s*_*i*_) values can be used to predict the order of species sensitivities to perturbations (⟨*s*_*i*_⟩ values). Alternatively, the order of species alignments with the leading eigenvector of the Jacobian matrix (|**v**_1*i*_| values) can be used to predict the order of ⟨*s*_*i*_⟩ values. (b) For the 3-species food chain model (equation (3)) at a given time, there are six possible ways to rank ⟨*s*_*i*_⟩ values, each one giving a Spearman’s rank correlation value (*ρ*). (c) Rank correlation (*ρ*) between 𝔼(*s*_*i*_) (computed analytically from the model) and ⟨*s*_*i*_⟩ over time quantified for a synthetic time series generated from the 3-species food chain model. The vast majority of points (97.2%) show a positive *ρ*. (d) Same as (c) but with the Jacobian matrix used to compute 𝔼(*s*_*i*_) inferred with the S-map using only past time-series data. Again, the great majority of points (94.8%) show a positive *ρ*.

#### Box 2

Eigenvector ranking

##### Rationale

This approach is based on the alignment of species *i* with the leading eigenvector (**v**_1_) of **J** (|**v**_1*i*_|). The solution of the linearized dynamics can also be written as 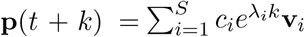, where **v**_*i*_ is the *i*th eigenvector of **J** associated with eigenvalue *λ*_*i*_ (*λ*_*S*_ ≤ … *≤ λ*_1_) and *c*_*i*_ are constants determined by the initial condition **p**(*t*) (*SI Section 9*) (Boyce *et al*., 2017, Strogatz, 2018). After a sufficient amount of time *k, λ*_1_ will dominate over other eigenvalues and the solution can be approximated by 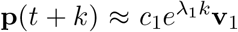. Thus, **v**_1_ dictates the local direction of greatest expansion (or smallest contraction) of perturbations. That is, the distribution of perturbed abundances (**Ñ**) will expand over time approximately along the direction of **v**_1_ and at a rate given by *λ*_1_ (positive values lead to expansion, whereas negative values lead to contraction). We also show that **v**_1_ serves as a proxy for the local leading Lyapunov vector, which provides the exact direction of perturbation growth under non-equilibrium dynamics (Kuptsov & Parlitz, 2012, Mease *et al*., 2003, Vallejo *et al*., 2017) (Fig. S4; *SI Section 10*). Specifically, we compute the alignment of species *i* with **v**_1_ as the absolute value of its *i*th element (|**v**_1*i*_|), where ||**v**_1_|| = 1. We define the order of |**v**_1*i*_| values across species as the eigenvector ranking and use it to predict the order of ⟨*s*_*i*_⟩ (*SI Section 11*). Note that we use the absolute value because only the line spanned by **v**_1_ and not its direction determines how perturbed abundances change over time. Also note that we use **v**_*i*_ and *λ*_*i*_ to denote the real parts of the *i*th eigenvector and eigenvalue, respectively. We show that our approach also holds for complex eigenvectors and eigenvalues, as long as their imaginary parts are small compared to real parts (*SI Section 9*).

##### Application

Similarly to the expected sensitivity ranking, the Jacobian matrix of the community (**J**) evaluated using the abundances (**N**) at time *t* is also required to apply the eigenvector ranking. The main advantage of the eigenvector ranking is that **J** is the only ingredient required to compute |**v**_1*i*_| and we do not need to specify the initial covariance matrix of perturbations (**Σ**_*t*_) nor the time for which perturbations evolve (*k*). Nevertheless, by using a single eigenvector instead of the entire Jacobian matrix, the eigenvector ranking uses less information than the expected sensitivity ranking. Importantly, we show that 𝔼(*s*_*i*_) and |**v**_1*i*_| are related in the special case of a symmetric Jacobian matrix (*SI Section 12*).

We illustrate how 𝔼(*s*_*i*_) (Box 1) and |**v**_1*i*_| (Box 2) allow us to predict the order of ⟨*s*_*i*_⟩ values under three simple scenarios of Lotka-Volterra dynamics at equilibrium (*SI Section 13*). We show that the order of 𝔼(*s*_*i*_) values is exactly the same as the order of (*s*_*i*_) values, whereas the order of |**v**_1*i*_| is similar to the order of ⟨*s*_*i*_⟩ values for all three scenarios (Figs. S1, S2, and S3). A potential limitation of these approaches to rank species sensitivities, however, is that they rely on knowledge of the parameterized population dynamics model **f** to obtain the Jacobian matrix **J**, which we rarely have. Therefore, in addition to computing 𝔼(*s*_*i*_) and |**v**_1*i*_| using the analytical **J**, we show that we can accurately rank ⟨*s*_*i*_⟩ by inferring **J** using the S-map, a data-driven method. The S-map is a locally weighted state-space regression method that has been shown to provide accurate inferences of the time-varying Jacobian matrix from abundance time series (Cenci *et al*., 2019, Deyle *et al*., 2016, Sugihara, 1994) (*SI Section 5*).

### Testing ranking approaches with synthetic time series

To test whether the order of expected sensitivities (𝔼(*s*_*i*_); Box 1) and eigenvector alignments (|**v**_1*i*_|; Box 2) can predict the order of species sensitivities (⟨*s*_*i*_⟩), we perform perturbation analyses using synthetic time series. Specifically, we generate multivariate synthetic time series with 500 points ({**N**(*t*)}, *t* = 1, …, 500) using five population dynamics models that produce non-equilibrium dynamics (Fig. S5; *SI Section 3*). Then, for half of each time series (*t* = 250, …, 500), we perform *n* = 300 random perturbations at each time *t*: **Ñ**= **N** + **p**, where **p** *∼* 𝒩 (**0, Σ**_*t*_) with **Σ**_*t*_ being a diagonal matrix with diagonal element *i* given by 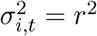 (i.e., independent normally distributed perturbations for each species with the same variance *r*^2^). We set *r* to be 15% of the mean standard deviation of species abundances, but relax this assumption in additional analyses (*SI Section 4*). Next, we numerically integrate the model **f** for a time *k* using each **Ñ**as an initial condition. Because the response of communities to perturbations can be highly dependent on the time scale of the dynamics (Hastings *et al*., 2018, Rinaldi & Scheffer, 2000, Sugihara, 1994), we set *k* to be inversely proportional to the mean rate of change of the dynamics (*SI Section 4*). Then, we compute ⟨*s*_*i*_⟩ from time *t* to *t* + *k* as well as 𝔼(*s*_*i*_) and |**v**_1*i*_| using **J** at time *t* as described in Box 1 and 2. Note that we compute 𝔼(*s*_*i*_) and |**v**_1*i*_| both analytically from the true Jacobian matrix of **f** evaluated at **N** and by sequentially inferring the Jacobian matrix with the S-map on 250 past points.

To assess how well the order of 𝔼(*s*_*i*_) and |**v**_1*i*_| predicts the order of ⟨*s*_*i*_⟩, we compute the Spearman’s rank correlation (*ρ*) between each ranking and ⟨*s*_*i*_⟩ at each time *t*. We focus on predicting the order instead of the exact values of (*s*_*i*_) for two important reasons. First, the exact values of ⟨*s*_*i*_⟩ depend on the initial covariance matrix **Σ**_*t*_ and on the time step *k*, which we rarely know for natural communities. Second, we can only infer an approximation of the Jacobian matrix with the S-map even from ideal time-series data (*SI Section 5* ; Cenci & Saavedra (2019)). We can illustrate our ranking procedure for 𝔼(*s*_*i*_) by considering the 3-species food chain model (Fig. 1; equation (3)). Note that we obtain similar results when using |**v**_1*i*_|. With 3 species, there are 6 possible ways to rank a given set of ⟨*s*_*i*_⟩ at any given time, resulting in 4 different *ρ* values (Fig. 2B). If the order of 𝔼(*s*_*i*_) matches the order of ⟨*s*_*i*_⟩ exactly, we obtain *ρ* = 1 (Fig. 2B). Otherwise, *ρ* decreases depending on the mismatch between the order of 𝔼(*s*_*i*_) and the order of ⟨*s*_*i*_⟩. Note that using *ρ* allows us to penalize prediction mistakes consistently, irrespective of whether the mistaken species are among the most or least sensitive ones. For example, in Fig. 2B, the second and third rankings have *ρ* = 0.5 because both contain one correct prediction, which is the least sensitive species in the second ranking and the most sensitive species in the third ranking. Under the 3-species food chain model, we find that the order of 𝔼(*s*_*i*_) matches the order of ⟨*s*_*i*_⟩ exactly (i.e., *ρ* = 1) for 48.8% of points and reasonably well (i.e., *ρ* = 0.5) for 48.4% of points in the time series (Fig. 2C). For 2.8% of points, however, the order of 𝔼(*s*_*i*_) is not a good predictor of the order of ⟨*s*_*i*_⟩ (i.e., *ρ* < 0). But most strikingly, we obtain very similar results when inferring 𝔼(*s*_*i*_) directly from the synthetic time series using the S-map (Fig. 2D), indicating that our approaches can be applied using only time-series data without any knowledge of the underlying model.

To benchmark our approaches, we use two simple single-species demographic indicators to predict the order of ⟨*s*_*i*_⟩ values. First, we use abundances absolute percent change between *t* − 1 and 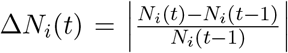. The rationale for this indicator is that a species will be more sensitive when its abundance is changing more rapidly. Second, we use abundances at time *t* after a sign reversal: *−N*_*i*_(*t*). This indicator is based on the notion that a species will be more sensitive when it has a low abundance, for example, due to strong density dependence effects. For both alternative indicators, we compute the rank correlation *ρ* between ⟨*s*_*i*_⟩ and the indicator at each time *t*. Note that computing Δ*N*_*i*_(*t*) or *−N*_*i*_(*t*) for a given species *i* only requires species-level information (i.e., abundance time series of species *i*) and not community-level information (i.e., Jacobian matrix) as 𝔼(*s*_*i*_) and |**v**_1*i*_| require.

We demonstrate the generality of our ranking approaches using a set of five synthetic non-equilibrium time series (*SI Section 3*). Although we find a high variation in *ρ* over time (gray points in Fig. 3A), the mean correlation 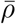 between ⟨*s*_*i*_⟩ and 𝔼(*s*_*i*_) as well as between ⟨*s*_*i*_⟩ and |**v**_1*i*_| is positive and high for all five models when we compute these rankings from the model (horizontal lines in Fig. 3A). In particular, we find that 𝔼(*s*_*i*_) shows the higher accuracy in ranking ⟨*s*_*i*_⟩, followed by |**v**_1*i*_|, Δ*N*_*i*_(*t*), and *−N*_*i*_(*t*). Note that we focus on 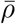 given that *ρ* is expected to vary over time due to changes in nonlinearity and rate of change of the Jacobian matrix. Importantly, we obtain very similar results for all models when inferring 𝔼(*s*_*i*_) and |**v**_1*i*_| using the S-map (horizontal lines in Fig. 3B). In addition to quantifying prediction accuracy, we can visualize how the value of ⟨*s*_*i*_⟩, 𝔼(*s*_*i*_), and |**v**_1*i*_| of each species changes over time (Fig. S6). We find that, even when inferring 𝔼(*s*_*i*_) and |**v**_1*i*_| with the S-map, we are able to detect shifts in ⟨*s*_*i*_⟩ across species (Fig. S6).

**Figure 3.**
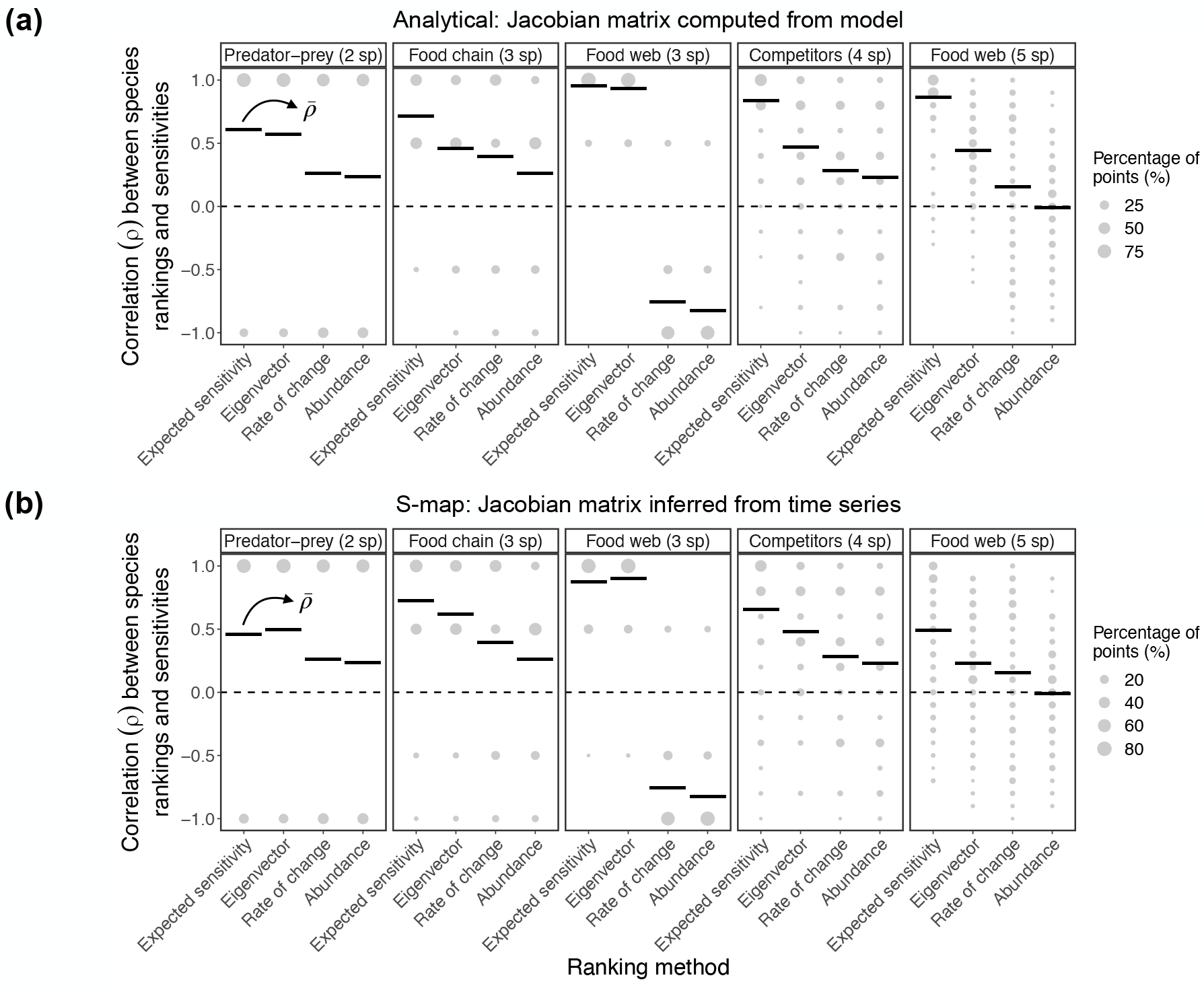
Expected sensitivity and eigenvector approaches allow us to accurately rank species sensitivities to perturbations under several population dynamics models. (a) Rank correlation (*ρ*) between species sensitivities to perturbations (⟨*s*_*i*_⟩) and four different approaches (expected sensitivity, 𝔼(*s*_*i*_); eigenvector, |**v**_1*i*_| ; rate of change, Δ*N*_*i*_(*t*); and abundance, *−N*_*i*_(*t*)). Note that the Jacobian matrix and, therefore, 𝔼(*s*_*i*_) and |**v**_1*i*_| are computed analytically from the model. Each panel shows the percentage of points with a given *ρ* value (size of gray points) and the average of these values across time (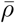, black horizontal lines) for a synthetic time series generated from the corresponding population dynamics model. (b) Same as (a) but with the Jacobian matrix and, therefore, 𝔼(*s*_*i*_) and **v**_1*i*_ inferred with the S-map using only past time-series data. In (a), the expected sensitivity approach shows a higher 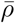 than the other three ranking approaches under all models. In (b), the expected sensitivity approach outperforms the eigenvector approach for three out of five models.

Although 𝔼(*s*_*i*_) is in general more accurate than |**v**_1*i*_|, we can use information on the leading eigenvalue (*λ*_1_) to increase the accuracy of the latter approach. Specifically, we expect that the higher *λ*_1_, the greater the local growth rate of perturbations in the direction of **v**_1_, which should improve our ability to rank ⟨*s*_*i*_⟩ values using |**v**_1*i*_|. Indeed, we find that 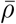 generally increases for the eigenvector approach when using only a subset of points with a high value of *λ*_1_ (Fig. S7). We find a positive correlation between the analytical and inferred *λ*_1_ (predator-prey (2 sp): 0.52; food chain (3 sp): 0.70; food web (3 sp): 0.57; competitors (4 sp): 0.23; and food web (5 sp): 0.70) and a high alignment between the analytical and inferred **v**_1_ for all models (Fig. S8). Note that *λ*_1_ is generally positive, whereas subsequent eigenvalues are negative or close to zero for the non-equilibrium attractors used here, implying that *λ*_1_ alone carries enough information to improve the eigenvector approach.

For most models, the expected sensitivity (𝔼(*s*_*i*_)) and eigenvector (|**v**_1*i*_|) approaches computed from the analytical Jacobian matrix show a high accuracy in ranking species sensitivities (⟨*s*_*i*_⟩) when using different perturbation distributions (Figs. S9 and S10) or time steps (*k*) to evolve perturbations (Figs. S11 and S12; *SI Section 4*). In particular, 𝔼(*s*_*i*_) shows an extremely high accuracy when *k* is small and fixed over time (e.g., *k* = 1; Fig. S11), given that the solution for the linearized dynamics (**p**(*t* + *k*)) is more precise for smaller values of *k*. In contrast, we find that |**v**_1*i*_| performs best when *k* depends on the time scale of the dynamics (Fig. 3), given that the eigenvector approach depends on the convergence of **p**(*t* + *k*) to the line spanned by **v**_1_, which requires a large *k* when dynamics are slower. We also find that the accuracy of 𝔼(*s*_*i*_) computed using wrong values of *k* and **Σ**_*t*_ remains high (Figs. S10 and S13), except when these values are greatly misspecified (Fig. S14). Finally, although the accuracy decreases in some cases, we find that 𝔼(*s*_*i*_) and |**v**_1*i*_| inferred with the S-map remain accurate when normalizing species abundances (Fig. S15), using shorter time series (Fig. S16), adding observational noise to the time series (Fig. S17), or adding process noise to the model (Fig. S18; *SI Section 6*).

### Detecting sensitive species in empirical time series

To illustrate the implementation of our data-driven framework, we apply our approaches to rank species sensitivities (Box 1 and 2) to two empirical time series. Each time series depicts a different marine community with four interacting variables that has been shown to exhibit non-equilibrium dynamics over long periods of time (*SI Section 7* ; Benincà *et al*. (2015, 2009)). Note that some variables represent physical attributes (e.g., bare rock) and others consist of species aggregations (e.g., barnacles) but we use the term species to refer to all variables. We first fit the S-map sequentially to both time series to infer 𝔼(*s*_*i*_) and |**v**_1*i*_| (*SI Section 7*). Because we do not know the laws governing population dynamics in these communities (i.e., **f** (**N**)), we cannot compute species sensitivities (⟨*s*_*i*_⟩). Instead, we perform abundance forecasts using a Long Short-Term Memory (LSTM) neural network (James *et al*., 2021) and test the hypothesis that species that are more sensitive to perturbations (i.e., have a higher value of 𝔼(*s*_*i*_) or |**v**_1*i*_|) at a given time will be harder to forecast (Cenci *et al*., 2020). The rationale behind this hypothesis is that, for a community under perturbations, the LSTM neural network will not be able to accurately forecast the abundance of a given species at a point in time when that species is highly sensitive to perturbations (Cenci *et al*., 2020). Both empirical communities described above are thought to be under perturbations triggered by changes in environmental conditions (Benincà *et al*., 2015, 2009).

For both the Jacobian matrix inference (i.e., S-map) and the abundance forecasts (i.e., LSTM neural network), we assign 70% of the data as a training set and use the remaining 30% as a test set. For each time *t* in the test set, we compute a standardized forecast root-mean-square error (RMSE) for each species *i* as (Perretti *et al*., 2013):

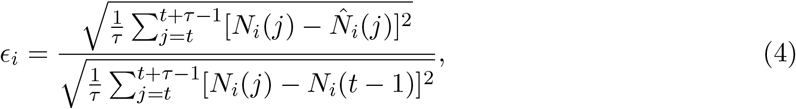

where *τ* = 3 is the number of forecasts, the numerator is the RMSE for the LSTM neural network forecast 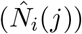 and the denominator is the RMSE for a naive forecast using the last point in the current training set (*N*_*i*_(*t −* 1)). We then compute the rank correlation (*ρ*) between 𝔼(*s*_*i*_) and *ϵ*_*i*_ as well as between |**v**_1*i*_| and *ϵ*_*i*_ at each point in the test set to test the hypothesis that our rankings can predict the order of forecast errors. We also compute the rank correlation between each of the two alternative indicators previously described (Δ*N*_*i*_(*t*) and *−N*_*i*_(*t*)) and *ϵ*_*i*_ to verify whether these simple indicators can predict the order of forecast errors. Because we do not know how perturbations affect these communities, we set **Σ**_*t*_ = **I** and *k* = *τ* to compute 𝔼(*s*_*i*_) (Box 1). We confirm the rationale behind our hypothesis by performing abundance forecasts under perturbations with the five synthetic time series used in our theoretical analyses (Fig. S19; *SI Section 8*).

We first illustrate the application of our approaches to detect sensitive species with the rocky intertidal community. We find that barnacles have the highest expected sensitivity (𝔼(*s*_*i*_)) followed by either algae or mussels depending on the point in time (Figs. 4A and S20). Interestingly, barnacles also show the highest alignment with the leading eigenvector (|**v**_1*i*_|) for the majority of points in time (Fig. S20). We also find consistent results for 𝔼(*s*_*i*_) and |**v**_1*i*_| with the marine plankton community (Figs. S20 and S21). Thus, although the values of 𝔼(*s*_*i*_) over time are different from those of |**v**_1*i*_|, these two rankings suggest some general patterns in how species sensitivities change over time in these communities. Importantly, we find that the mean rank correlation 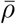 (computed over the test set) between 𝔼(*s*_*i*_) and *ϵ*_*i*_ is positive for both time series, but only significant for one of them (rocky intertidal community: 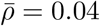, *p*-value = 0.287, 1,000 randomizations; marine plankton community: 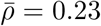, *p*-value < 0.001; Fig. 4B). However, we find that the mean rank correlation 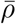 between |**v**_1*i*_| and *ϵ*_*i*_ is positive and significant for both time series (rocky intertidal community: 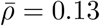, *p*-value = 0.023; marine plankton community: 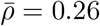, *p*-value < 0.001; Fig. 4B).

**Figure 4.**
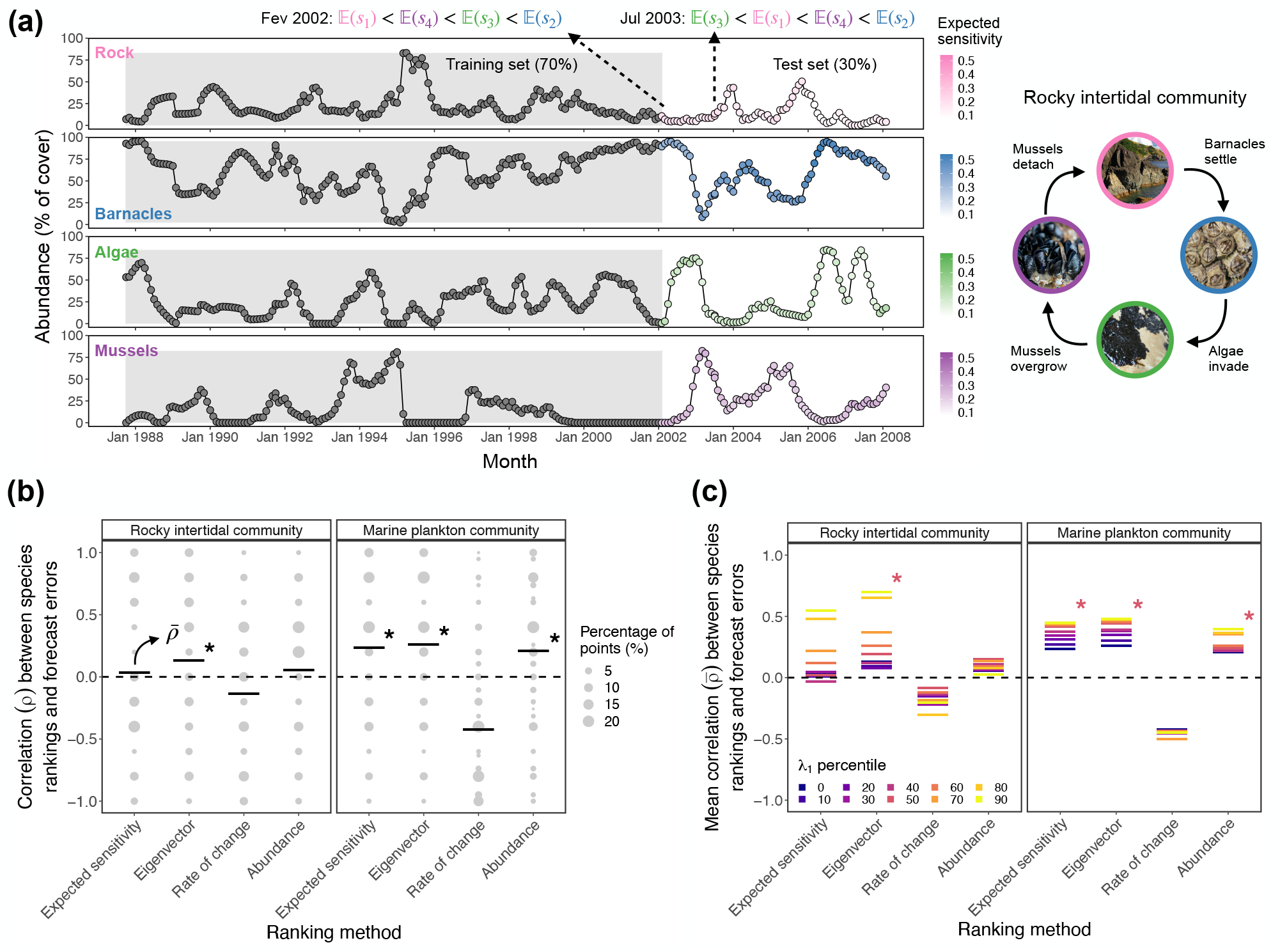
Species abundance forecast errors are associated with species sensitivities to perturbations. (a) Time series of a rocky intertidal community containing four variables (bare rock, barnacles, algae, and mussels). The diagram on the right depicts the cyclic succession in this community (adapted from Benincà *et al*. (2015)). Note that percentage of cover does not necessarily sum to 100% as individuals of different species may overlap on top of the rock. We use a moving training set (gray region) to train the S-map and compute expected sensitivities (𝔼(*s*_*i*_)) as well as species alignments with the leading eigenvector (|**v**_1*i*_|) at the last point in the training set. Simultaneously, we train an LSTM neural network to forecast species abundances and compute species forecast errors (*ϵ*_*i*_). Barnacles (blue) show the highest value of 𝔼(*s*_*i*_) followed by either algae (green) or mussels (purple) depending on the point in time. Note that 𝔼(*s*_*i*_) values across species sum to 1 for each point in time (darker points denote higher E(*s*_*i*_)). (b) Rank correlation (*ρ*) between *ϵ*_*i*_ and four different approaches (expected sensitivity, 𝔼(*s*_*i*_); eigenvector, |**v**_1*i*_ |; rate of change, Δ*N*_*i*_(*t*); and abundance, *−N*_*i*_(*t*)). Each panel shows the percentage of points with a given *ρ* value (size of gray points) and the average of these values across the test set (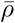, black horizontal lines) for a given empirical time series (asterisks denote a *p*-value less than 0.05 for 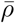 according to a randomization test). (c) Average correlation 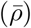 between *ϵ*_*i*_ and the different ranking approaches computed for points in the test set with a *λ*_1_ value higher than a given percentile of the *λ*_1_ distribution. For the expected sensitivity and eigenvector approaches, 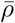 increases as we only use points with successively higher values of *λ*_1_ for both time series (asterisks denote a *p*-value less than 0.05 for 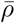 using the 50th percentile). Pictures are under the Creative Commons License: rock by Piotr Zurek, barnacles by tangatawhenua, algae by redrovertracy, and mussels by Wayne Martin.

We find further evidence that species with higher 𝔼(*s*_*i*_) or |**v**_1*i*_| are harder to forecast by computing 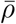 only for points in the test set with successively higher values of *λ*_1_ (i.e., higher local growth rate of perturbations; Fig. 4C). For example, as expected from our analyses with synthetic time series (Fig. S7), 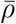 between |**v**_1*i*_| and *ϵ*_*i*_ increases for both time series when we only use points for which *λ*_1_ is higher than its 50th percentile (rocky intertidal community: 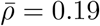, *p*-value = 0.021; marine plankton community: 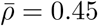, *p*-value < 0.001; Fig. 4C). The fact that 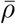 does not increase in general by using only points with a high *λ*_1_ for the alternative indicators (Δ*N*_*i*_(*t*) and *−N*_*i*_(*t*)) supports our ranking approaches in linking species forecast errors to their sensitivities to perturbation (Fig. 4C). We find these results to remain similar when changing the size of the training set and the number of steps ahead (*τ*) to forecast (Figs. S22, S23, and S24) as well as to normalizing species abundances before performing the S-map (Fig. S25).

## Discussion

Understanding how individual species affect the response to perturbations of the whole community and, in turn, how species interactions at the community level affect responses of individual species is paramount to ecological management and conservation (Beauchesne *et al*., 2021, Clark *et al*., 2021, Kéfi *et al*., 2019, Levin & Lubchenco, 2008). Yet, the traditional focus of ecology on recovery to equilibrium using parameterized models has been hampering efforts to understand how species respond to perturbations when community dynamics are out of equilibrium. Here we introduce a data-driven framework to solve a previously unexplored problem: how to rank the species that compose a community according to their sensitivity to perturbations under non-equilibrium dynamics? Our findings provide three main insights into how communities and their constituent species respond to perturbations.

First, we show that information on the time-varying local effects between interacting species (i.e., Jacobian matrix) can be used to determine which species will be most affected by perturbations at a given time. In particular, using dynamical systems theory (Arnoldi *et al*., 2018, Mease *et al*., 2003, Strogatz, 2018) and nonlinear time series methods (Cenci *et al*., 2019, Deyle *et al*., 2016, Sugihara, 1994), we develop two complementary approaches that can accurately rank species from most to least sensitive to small perturbations on abundances under non-equilibrium dynamics. Both the expected sensitivity and the eigenvector ranking allow us to detect which of the species that compose a natural community are the most and least sensitive in real time if a long time series is available. Hence, it may be possible to inform management and conservation programs regarding which species are currently the most sensitive ones. Our measure of sensitivity uses community-level information to quantify the likelihood of large changes (either decreases or increases) in the abundance of a given species. Therefore, species sensitivities may complement current demographic indicators that estimate single-species vulnerability to perturbations (Caswell, 2000, Mace *et al*., 2008, Morris *et al*., 2002). That is, while some species obviously require constant monitoring due to a high extinction risk (Dirzo *et al*., 2014, Estes *et al*., 2011), other species may require more attention during periods of time when they have a high sensitivity, irrespective of their abundance. It is worth noting, however, that our results cannot be extrapolated beyond a given studied community as our framework uses information on that specific community to rank species sensitivities.

It is important to note that the expected sensitivity ranking is more accurate than the eigenvector ranking for most of our perturbation analyses with synthetic time series. In particular, the expected sensitivity ranking has its best performance when the time over which perturbations evolve (*k*) is small and fixed, and its worse performance when the covariance matrix of perturbations (**Σ**_*t*_) and *k* are greatly misspecified. In contrast, the eigenvector ranking has the advantage of not depending on **Σ**_*t*_ and *k* for its computation and has its best performance when *k* depends on the local time scale of the dynamics. Indeed, in the non-equilibrium communities found in nature, large differences in time scale and, therefore, in the time it takes for perturbation effects to appear are widespread (Hastings *et al*., 2018, Rinaldi & Scheffer, 2000, Strogatz, 2018). Thus, it is reasonable to expect that as a practical tool the heuristic eigenvector ranking may be as useful as the more theoretically complete but assumption bound expected sensitivity ranking.

Second, we find support for our hypothesis that the abundance forecast errors for the different species in a community are associated with their sensitivity to perturbations. The predictability of ecological dynamics is known to change across communities (Dietze, 2017, Pennekamp *et al*., 2019) and, for a single community, across time (Cenci *et al*., 2020). In particular, it has been shown that at points in time when a community is more sensitive to perturbations its local predictability can be lower and, therefore, the average abundance forecast error can be higher (Cenci *et al*., 2020). Here, we have extended this result for individual species by showing that the local predictability of a given species is associated with how sensitive this species is to perturbations, which we infer through its expected sensitivity and its alignment with the leading eigenvector. The fact that the correlation between species forecast errors and our ranking approaches strengthens when the leading eigenvalue is high (i.e., perturbations grow rapidly along a given direction in state space) further supports our hypothesis that species forecast errors are associated with their sensitivities. In addition, the better performance with empirical data of the eigenvector approach in relation to the expected sensitivity approach suggests that we may be misspecifying the information required to compute expected sensitivities. These results provide empirical support to the eigenvector approach as a way to detect sensitive species using minimal information inferred from time-series data. Overall, our findings suggest that sensitivity to perturbations is an additional factor influencing the intrinsic predictability of different species in ecological communities (Dietze, 2017, Pennekamp *et al*., 2019).

Applying our ranking approaches to empirical data requires an accurate inference of the time-varying Jacobian matrix with the S-map. Although the S-map has been shown to provide accurate inferences when time series are noisy (Cenci *et al*., 2019, Deyle *et al*., 2016), several limitations remain. Because information on the shape of the attractor is required to fit the S-map, longer time series with smaller amounts of noise improve inference quality, all else being equal. In our analyses with synthetic time series, we show that our ranking approaches remain accurate when using shorter time series (Fig. S16) or under small amounts of noise (Figs. S17 and S18). Long time series with a strong signal of non-equilibrium deterministic dynamics, such as the rocky intertidal or plankton community investigated here (Benincà *et al*., 2015, 2009), are examples of ideal data sets to apply our ranking approaches. Although here we focused on small communities and small amounts of noise, future work may combine our ranking approaches with recent improvements of the S-map (e.g., regularization and multiview distance; Cenci *et al*. (2019), Chang *et al*. (2021)) to detect sensitive species under more challenging settings.

Finally, we show that approaches based on linear dynamical systems that are typically used for communities close to equilibrium can also provide information for communities under non-equilibrium dynamics (Cenci & Saavedra, 2019, Ushio *et al*., 2018). Although the methodology may be similar in both cases, it is important to note that its interpretation is completely different. For example, whereas the linearized dynamics can be used to compute a recovery rate under equilibrium (Arnoldi *et al*., 2018, Medeiros *et al*., 2021, Strogatz, 2018), we show that they can be used to derive the time-varying expected sensitivity of different species to perturbations under non-equilibrium dynamics. Similarly to previous studies, our results confirm the utility of tracking variances and covariances of perturbed abundances to anticipate changes in ecological communities (Baruah *et al*., 2022, Chen *et al*., 2019, Dakos, 2018). In addition, we use the leading eigenvector, which has been previously employed to decompose community responses into species responses to perturbations under equilibrium dynamics (Dakos, 2018, Ghadami *et al*., 2020, Patterson *et al*., 2021, Weinans *et al*., 2019). Thus, the approaches introduced here increase our understanding of how communities and their constituent species respond to perturbations when there is no stable equilibrium. Both approaches are based on a linearization of the dynamics and, thereby, only provide an assessment of responses to small perturbations. Moreover, both approaches assume that the Jacobian matrix does not change much over the time period for which perturbations evolve. Improving our framework to deal with strong nonlinearities, fast changes in the Jacobian matrix, and local oscillations due to complex eigenvalues are promising avenues for future research. Overall, our findings illustrate how integrating well-known results of equilibrium dynamics with data-driven methods for non-equilibrium dynamics provides a fruitful avenue for future development and new insights on how to understand the response of single species and entire communities to perturbations.

## Acknowledgments

We thank D. Rothman, T. Lieberman, S. Cenci, C. Song, J. Deng, and M. AlAdwani for insightful discussions and suggestions regarding this work. L.P.M. was supported by MIT ESI and the Martin Family Society of Fellows for Sustainability. G.S. was supported by DoD-Strategic Environmental Research and Development Program 15 RC-2509, NSF DEB-1655203, NSF ABI-1667584, and DOI USDI-NPS P20AC00527. S.S. was supported by NSF DEB-2024349.

