## Supporting Information for "Ranking species based on sensitivity to perturbations under non-equilibrium community dynamics"

### Contents

**1 Derivation of the dynamics of small perturbations 2**

**2 Derivation of analytical expected sensitivity 2**

**3 Synthetic time series from population dynamics models 4**

**4 Perturbation analyses 6**

**5 Inference of Jacobian matrix with the S-map 6**

**6 Analyses with short and noisy synthetic time series 7**

**7 Forecast analyses with empirical time series 8**

**8 Forecast analyses with synthetic time series 9**

**9 Leading eigenvector and direction of greatest perturbation expansion under equilibrium dynamics 10**

**10 Leading Lyapunov vector and direction of greatest perturbation expansion under non-equilibrium dynamics 12**

**11 From direction of greatest perturbation expansion to ranking species sensitivities 13**

**12 Connection between expected sensitivity and eigenvector approaches 14**

**13 Illustrations with Lotka-Volterra dynamics at equilibrium 15**

### 1 Derivation of the dynamics of small perturbations

In this section, we provide a derivation of the linear dynamics of small perturbations, which is the foundation of our approaches to rank species sensitivities to perturbations. Let us consider the most general form of a population dynamics model for a given species $i$ within a community with $S$ species (Case, 2000):

$$\frac{dN_i}{dt} = f_i(\mathbf{N}), \tag{S1}$$

where $N_i$ is the abundance of species $i$, $\mathbf{N} = [N_1, ..., N_S]^\top$ is the vector of abundances of all species, and $f_i$ ($f_i$: $\mathbb{R}^S \to \mathbb{R}$) is the function describing how the growth rate of species $i$ depends on the abundances of all species. Note that $f_i$ also depends on a set of parameters, which we consider to be fixed over time. We can write equation (S1) for all species in the community as $\frac{d\mathbf{N}}{dt} = \mathbf{f}(\mathbf{N})$, where $\frac{d\mathbf{N}}{dt} = [\frac{dN_1}{dt}, ..., \frac{dN_S}{dt}]^\top$ and $\mathbf{f}$: $\mathbb{R}^S \to \mathbb{R}^S$. See below (*Section 3*) for some examples of population dynamics models of this form.

In this study, we are interested in ranking species according to their sensitivity to perturbations, that is, how much their abundance trajectories are expected to change after some time following a small random wiggle on abundances. Then, let us consider a random pulse perturbation $\mathbf{p}$ that changes $\mathbf{N}$ into $\tilde{\mathbf{N}}$ (i.e., $\tilde{\mathbf{N}} = \mathbf{N} + \mathbf{p}$). Now, we can write the Taylor expansion of $\frac{d\tilde{\mathbf{N}}}{dt}$ around $\mathbf{N}$ (Strogatz, 2018):

$$\frac{d\tilde{\mathbf{N}}}{dt} = \mathbf{f}(\mathbf{N}) + \frac{\partial \mathbf{f}}{\partial \tilde{\mathbf{N}}}\Big|_{\tilde{\mathbf{N}}=\mathbf{N}} \cdot (\tilde{\mathbf{N}} - \mathbf{N}) + O(\mathbf{p}^\top \mathbf{p}), \tag{S2}$$

where $\frac{\partial \mathbf{f}}{\partial \mathbf{N}} = \mathbf{J}$ is the Jacobian matrix of partial derivatives with $j_{ij} = \frac{\partial f_i}{\partial N_j}$. If $\mathbf{p}$ is small, we can approximate its dynamics by taking just the linear term (i.e., ignoring higher-order terms):

$$\begin{aligned} \frac{d\tilde{\mathbf{N}}}{dt} &= \mathbf{f}(\mathbf{N}) + \frac{\partial \mathbf{f}}{\partial \tilde{\mathbf{N}}}\Big|_{\tilde{\mathbf{N}}=\mathbf{N}} \cdot (\tilde{\mathbf{N}} - \mathbf{N}) \\ \frac{d\mathbf{N}}{dt} + \frac{d\mathbf{p}}{dt} &= \frac{d\mathbf{N}}{dt} + \mathbf{J}|_{\tilde{\mathbf{N}}=\mathbf{N}} \cdot \mathbf{p} \\ \frac{d\mathbf{p}}{dt} &= \mathbf{J}|_{\tilde{\mathbf{N}}=\mathbf{N}} \cdot \mathbf{p}. \end{aligned} \tag{S3}$$

Thus, as it is known (Boyce *et al.*, 2017, Kuptsov & Parlitz, 2012, Mease *et al.*, 2003, Strogatz, 2018, Vallejo *et al.*, 2017), the dynamics of a small perturbation $\mathbf{p}$ can be approximated by the linear equation above called the tangent dynamics of $\frac{d\mathbf{N}}{dt}$. Note that we have not assumed the existence of an equilibrium here (i.e., $\mathbf{N}^*$ for which $\mathbf{f}(\mathbf{N}^*) = \mathbf{0}$) and, therefore, equation (S3) is valid irrespective of whether $\mathbf{N}$ is close to equilibrium or not.

### 2 Derivation of analytical expected sensitivity

Here we derive the expected value ($\mathbb{E}(s_i)$; Box 1 in the main text) of the sensitivity $s_i$ (equation (1) in the main text) of species $i$ to small perturbations ($\mathbf{p}$) affecting species abundances ($\mathbf{N}$). We assume that $\mathbf{p}(t)$ follows a distribution with mean zero and covariance matrix $\mathbf{\Sigma}_t$. We assume a distribution with mean zero because unbiased perturbations are the most uninformative way to consider how perturbations may impact a community. In most of our perturbation analyses, we assume that $\mathbf{p}(t)$ follows a multivariate normal distribution (i.e., $\mathbf{p}(t) \sim \mathcal{N}(\mathbf{0}, \mathbf{\Sigma}_t)$), but this assumption is not necessary for the derivation below. The linearized dynamics of a small perturbation is given by $\frac{d\mathbf{p}}{dt} = \mathbf{J}\mathbf{p}$ (see *Section 1*) (Boyce *et al.*, 2017, Eckmann & Ruelle, 1985, Mease *et al.*, 2003, Strogatz, 2018). We can obtain the solution for this linear system as $\mathbf{p}(t+k) = e^{\mathbf{J}k}\mathbf{p}(t)$, where $e^{\mathbf{A}} = \sum_{i=1}^{\infty} \frac{1}{i!}\mathbf{A}^i$ is the exponential of matrix $\mathbf{A}$ (Arnoldi *et al.*, 2018, Boyce *et al.*, 2017). By defining $\mathbf{M} = e^{\mathbf{J}k}$, we can compute the expected value of $\mathbf{p}(t+k)$:

$$\begin{aligned} \mathbb{E}[\mathbf{p}(t+k)] &= \mathbb{E}[\mathbf{M}\mathbf{p}(t)] \\ &= \mathbf{M}\mathbb{E}[\mathbf{p}(t)] \\ &= \mathbf{0}. \end{aligned} \tag{S4}$$

Thus, $\mathbf{p}(t+k)$ also follows a distribution with mean zero. In the special case where $\mathbf{p}(t)$ follows a normal distribution, $\mathbf{p}(t+k)$ also follows a normal distribution because $\mathbf{M}\mathbf{p}(t)$ is a weighted sum of normal distributions.

Because $p_i(t)$ and $p_i(t+k)$ have mean zero, the sensitivity of species $i$ can be approximated by the ratio of the variance of $p_i(t+k)$ and the variance of $p_i(t)$:

$$\langle s_i \rangle = \frac{\frac{1}{n}\sum_{j=1}^{n} p_i^{(j)}(t+k)^2}{\frac{1}{n}\sum_{j=1}^{n} p_i^{(j)}(t)^2} = \frac{\mathrm{Var}[p_i(t+k)]}{\mathrm{Var}[p_i(t)]}, \tag{S5}$$

where $\mathrm{Var}[p_i(t)] = \sigma_{i,t}^2$ is the $i$th diagonal element of $\mathbf{\Sigma}_t$. Assuming that $\sigma_{i,t}^2$ is the same for every species $i$, we can ignore it for the purpose of ranking species sensitivities and focus only on $\mathrm{Var}[p_i(t+k)]$. We can obtain $\mathrm{Var}[p_i(t+k)]$ by computing the covariance matrix of $\mathbf{p}(t+k)$:

$$\begin{aligned} \mathbf{\Sigma}_{t+k} &= \mathbb{E}[\mathbf{p}(t+k)\mathbf{p}(t+k)^\top] \\ &= \mathbb{E}[(\mathbf{M}\mathbf{p}(t))(\mathbf{M}\mathbf{p}(t))^\top] \\ &= \mathbf{M}\mathbb{E}[\mathbf{p}(t)\mathbf{p}(t)^\top]\mathbf{M}^\top \\ &= \mathbf{M}\mathbf{\Sigma}_t\mathbf{M}^\top. \end{aligned} \tag{S6}$$

Therefore, we define the expected sensitivity of species $i$ at time $t$ as: $\mathbb{E}(s_i) = \mathrm{Var}[p_i(t+k)] = \sigma_{i,t+k}^2$, where $\sigma_{i,t+k}^2$ is the $i$th diagonal element of $\mathbf{\Sigma}_{t+k}$. Note that we can normalize $\mathbb{E}(s_i)$ by dividing it by $\sum_{i=1}^{S} \sigma_{i,t+k}^2$, which has been shown to correspond to the expected magnitude of $\mathbf{p}(t+k)$ (i.e., $\mathbb{E}[||\mathbf{p}(t+k)||^2]$) (Arnoldi *et al.*, 2018). Although this normalization does not change the order of $\mathbb{E}(s_i)$ values, it allows us to interpret the normalized $\mathbb{E}(s_i)$ as the relative contribution of species $i$ to the expected magnitude of $\mathbf{p}(t+k)$.

In addition to knowing $\mathbf{J}$, knowledge of $\mathbf{\Sigma}_t$ and $k$ is required to compute $\mathbb{E}(s_i)$. In our main set of perturbation analyses, we compute $\mathbb{E}(s_i)$ using the true value of $k$ used to evolve perturbed abundances but do not use the true value of $\mathbf{\Sigma}_t$. Specifically, we set $\mathbf{\Sigma}_t = \mathbf{I}$, where $\mathbf{I}$ is the identity matrix. We test the robustness of the expected sensitivity ranking under uncertainty in $k$ and $\mathbf{\Sigma}_t$ in three different ways. First, we compute $\mathbb{E}(s_i)$ using $\mathbf{\Sigma}_t = \mathbf{I}$ when $\sigma^2_{i,t}$ varies over time and across species (i.e., normally distributed perturbations with a variance proportional to relative species abundances; Fig. S10). Second, we compute $\mathbb{E}(s_i)$ using $k = 1$ when $k$ varies over time (i.e., $k$ inversely proportional to the time scale of the dynamics; Fig. S13). Third, we compute $\mathbb{E}(s_i)$ as described above for our main set of analyses but add 100% of normally distributed noise to $\mathbf{\Sigma}_t$ and $k$ at each point in time (Fig. S14).

### 3 Synthetic time series from population dynamics models

To test whether expected sensitivities ($\mathbb{E}(s_i)$; Box 1 in the main text) and species alignments with the leading eigenvector ($|\mathbf{v}_{1i}|$; Box 2 in the main text) can accurately rank species sensitivities to perturbations ($\langle s_i \rangle$, equation (2) in the main text), we perform perturbation analyses using synthetic time series. We generate synthetic time series using five different population dynamics models with the generic form: $\frac{d\mathbf{N}}{dt} = \mathbf{f}(\mathbf{N})$, where $\mathbf{f}\colon \mathbb{R}^S \to \mathbb{R}^S$ is a nonlinear function. Here, we present the equations, parameter values and references for each model.

The first model contains two species and depicts the interactions between a predator (species 1) and its prey (species 2), producing a limit cycle (Yodzis, 1989) (Fig. S5):

$$\begin{aligned} \frac{dN_1}{dt} &= kN_1\Big(\frac{aN_2^2}{1+ahN_2^2}\Big) - dN_1 \\ \frac{dN_2}{dt} &= rN_2\Big(1-\frac{N_2}{K}\Big) - N_1\Big(\frac{aN_2^2}{1+ahN_2^2}\Big), \end{aligned} \tag{S7}$$

where $k = 0.5$ $a = 0.002$, $h = 4$, $d = 0.1$ $r = 0.5$, and $K = 100$.

The second model contains three species and depicts a food chain with a primary producer (species 1), a primary consumer (species 2), and a secondary consumer (species 3), producing chaotic dynamics (Hastings & Powell, 1991, Upadhyay, 2000) (Fig. 1 in the main text and S5):

$$\begin{aligned} \frac{dN_1}{dt} &= rN_1\Big(1-\frac{N_1}{K}\Big) - \frac{a_1N_1N_2}{1+b_1N_1} \\ \frac{dN_2}{dt} &= -sN_2 + hN_1N_2 - \frac{a_2N_2N_3}{1+b_2N_2} \\ \frac{dN_3}{dt} &= -lN_3 + nN_2N_3, \end{aligned} \tag{S8}$$

where $r = 4.3$, $K = 50$, $a_1 = 0.1$, $b_1 = 0.1$, $a_2 = 0.1$, $b_2 = 0.1$, $s = 1$, $h = 0.05$, $l = 1$, and $n = 0.03$.

The third and fourth models have the general form of the classic Lotka-Volterra model (Case, 2000):

$$\frac{dN_i}{dt} = N_i\Big(r_i + \sum_{j=1}^{S} a_{ij}N_j\Big) \tag{S9}$$

where $r_i$ is an element of the vector $\mathbf{r}$ representing the intrinsic growth rate of species $i$ and $a_{ij}$ is an element of the interaction matrix $\mathbf{A}$ representing the interaction effect of species $j$ on species $i$. The third model contains three species ($S = 3$) and produces chaotic dynamics between two prey and one predator (Gilpin, 1979) (Fig. S5) with the following values for $r_i$ and $a_{ij}$:

$$\mathbf{r} = \begin{bmatrix} 1 \\ 1 \\ -1 \end{bmatrix}, \mathbf{A} = \begin{bmatrix} -0.1 & -0.1 & -1 \\ -0.15 & -0.1 & -0.1 \\ 0.5 & 0.05 & 0 \end{bmatrix}$$

The fourth model contains four competitor species ($S = 4$) and also produces chaotic dynamics (Vano *et al.*, 2006) (Fig. S5) with the following values for $r_i$ and $a_{ij}$:

$$\mathbf{r} = \begin{bmatrix} 1 \\ 0.72 \\ 1.53 \\ 1.27 \end{bmatrix}, \mathbf{A} = \begin{bmatrix} -1 & -1.09 & -1.52 & 0 \\ 0 & -1 & -0.44 & -1.36 \\ -2.33 & 0 & -1 & -0.47 \\ -1.21 & -0.51 & -0.35 & -1 \end{bmatrix}$$

Finally, the fifth model depicts a 5-species food web with two secondary consumers (species 1 and 2), two primary consumers (species 3 and 4), and one primary producer (species 5) also generating chaotic dynamics (Deyle *et al.*, 2016) (Fig. S5):

$$\begin{aligned} \frac{dN_1}{dt} &= \nu_1\lambda_1\frac{N_1N_3}{N_3+N_3^*} - \nu_1N_1 \\ \frac{dN_2}{dt} &= \nu_2\lambda_2\frac{N_2N_4}{N_4+N_4^*} - \nu_2N_2 \\ \frac{dN_3}{dt} &= \mu_1\kappa_1\frac{N_3N_5}{N_5+N_5^*} - \nu_1\lambda_1\frac{N_1N_3}{N_3+N_3^*} - \mu_1N_3 \\ \frac{dN_4}{dt} &= \mu_2\kappa_2\frac{N_4N_5}{N_5+N_5^*} - \nu_2\lambda_2\frac{N_2N_4}{N_4+N_4^*} - \mu_2N_4 \\ \frac{dN_5}{dt} &= N_5\left(1-\frac{N_5}{K}\right) - \mu_1\kappa_1\frac{N_3N_5}{N_5+N_5^*} - \mu_2\kappa_2\frac{N_4N_5}{N_5+N_5^*}, \end{aligned} \tag{S10}$$

where $\nu_1 = 0.1$, $\nu_2 = 0.07$, $\lambda_1 = 3.2$, $\lambda_2 = 2.9$, $N_3^* = 0.5$, $N_4^* = 0.5$, $\mu_1 = 0.15$, $\mu_2 = 0.15$, $\kappa_1 = 2.5$, $\kappa_2 = 2$, $N_5^* = 0.3$, and $K = 1.2$.

For each model, we numerically integrate the dynamics using a Runge-Kutta method with a time step of 0.05 and obtain a time series with 10,000 points. Then, we sample equidistant points obtaining a final multivariate time series with 500 points ($\{\mathbf{N}(t)\}$, $t = 1, ..., 500$). Note that with this protocol we obtain time series that fully sample the attractor of each model and have a size similar to the empirical time series used here (Fig. S5). Also note that by sampling equidistant points we test the robustness of the S-map to infer $\mathbb{E}(s_i)$ and $|\mathbf{v}_{1i}|$ under the typical low sampling frequency of empirical time series.

### 4 Perturbation analyses

For each synthetic time series, we perform random perturbations on abundances to compute species sensitivities ($\langle s_i \rangle$; equation (2) in the main text). We apply $n = 300$ random pulse perturbations $\mathbf{p}$ to the abundance vector $\mathbf{N}$ at each point in time: $\tilde{\mathbf{N}} = \mathbf{N} + \mathbf{p}$. We perform these perturbations in three different ways. First, we assume perturbations are normally distributed around $\mathbf{N}$ and use $p_i(t) \sim \mathcal{N}(\mu = 0, \sigma^2 = r^2)$ (Fig. 1C, D in the main text). Second, we assume perturbations are uniformly distributed around $\mathbf{N}$ and apply $\mathbf{p}(t)$ such that $\tilde{\mathbf{N}}$ is uniformly distributed inside a hypersphere of radius $r$ centered in $\mathbf{N}$. Third, we assume normally distributed perturbations with a variance proportional to relative species abundances, such that: $p_i(t) \sim \mathcal{N}(\mu = 0, \sigma^2 = N_i'(t) r^2)$, where $N_i'(t) = \frac{N_i(t)}{\sum_{i=1}^{S} N_i(t)}$. Note that in this last scenario we relax our assumption that the variance of $p_i(t)$ is fixed over time and equal for every species. For all types of perturbation, we set $r$ to be 15% of the mean standard deviation of species abundances: $r = 0.15 \frac{1}{S} \sum_{i=1}^{S} \sigma_{N_i}$, where $\sigma_{N_i}$ is the standard deviation of $N_i$ for the whole time series. The results for normally distributed perturbations are presented in the main text, whereas the results for the other perturbation types are shown in Figs. S9 and S10.

After applying perturbations, we numerically integrate model $\mathbf{f}$ for $k$ time steps using each perturbed abundance vector $\tilde{\mathbf{N}}$ as well as the unperturbed abundance vector $\mathbf{N}$ as initial conditions. Then, we compute $\langle s_i \rangle$ using the initial (i.e., time $t$) and final (i.e., time $t + k$) perturbed and unperturbed abundances (equation (2) in the main text). Because $\frac{d\mathbf{N}}{dt}$ (i.e., time scale) can greatly vary across state space, impacting how perturbations grow over time, we set $k$ to be inversely proportional to the mean absolute percent change between $N_i(t+1)$ and $N_i(t)$. Specifically, we use $k = \left[ \frac{1}{S} \sum_{i=1}^{S} \left| \frac{N_i(t+1) - N_i(t)}{N_i(t)} \right| \right]^{-\frac{1}{2}}$. Thus, $k$ increases as the percent change decreases and we use a square root to damp the large variability in time scale found for most models. We also perform these analyses using a fixed value of $k$ ($k = 1$ or $k = 3$) for all points in the time series (Figs. S11 and S12). Note that $k = 3$ can be considered a long time period for some models, allowing us to test the robustness of our approaches for longer periods of time.

### 5 Inference of Jacobian matrix with the S-map

We perform the S-map using the `rEDM` package in R to sequentially infer the Jacobian matrix ($\mathbf{J}$) through time using only past time-series data in order to compute expected sensitivities ($\mathbb{E}(s_i)$) and species alignments with the leading eigenvector ($|\mathbf{v}_{1i}|$). The S-map is a locally weighted state-space regression method that can be used to infer the time-varying Jacobian matrix of a dynamical system (Cenci *et al.*, 2019, Deyle *et al.*, 2016, Sugihara, 1994). Given a time series ($\{\mathbf{N}(t_k)\}$, $k = 1, ..., T$), we can fit a linear regression of the following form to each point: $N_i(t_k + 1) = c_{i0} + \sum_{j=1}^{S} c_{ij} N_j(t_k)$. Note that $c_{ij} = \frac{\partial N_i(t_k+1)}{\partial N_j(t_k)}$ is an approximation of the Jacobian element $j_{ij}$ at time $t_k$. The S-map consists of performing this linear regression locally for a given target point $\mathbf{N}(t^*)$ by giving a stronger weight to points that are closer to it in state space. This is done by finding a solution for $\mathbf{c}$ in $\mathbf{b} = \mathbf{A}\mathbf{c}$, where $b_k = w_k N_i(t_k + 1)$, $a_{kj} = w_k N_j(t_k)$, $w_k = \exp\left[ -\theta \frac{||\mathbf{N}(t_k) - \mathbf{N}(t^*)||}{\bar{d}} \right]$, and $\overline{d} = \frac{1}{T}\sum_{k=1}^{T} ||\mathbf{N}(t_k) - \mathbf{N}(t^*)||$. Thus, $\mathbf{b} \in \mathbb{R}^T$ contains the abundances at $t_k + 1$ weighted by the relative distance of each point to the target point, $\mathbf{A} \in \mathbb{R}^{T\times(S+1)}$ is the weighted data matrix of abundances at $t_k$, and $\mathbf{c} \in \mathbb{R}^{S+1}$ estimates the $i$th row of the Jacobian matrix at time $t_k$ as well as an intercept term. We obtain the solution for $\mathbf{c}$ via singular value decomposition (Deyle *et al.*, 2016), which is equivalent to the ordinary least squares solution (Cenci *et al.*, 2019). Importantly, the parameter $\theta$ tunes how strongly the regression is localized around the target point and is typically selected using leave-one-out cross-validation (LOOCV) (Cenci *et al.*, 2019).

For each of the five synthetic time series, we fit the S-map sequentially to infer $\mathbf{J}$ for each point in time, which is then used to compute $\mathbb{E}(s_i)$ and $|\mathbf{v}_{1i}|$. To do so, we assign half of the time series (i.e., $\{\mathbf{N}(t)\}$, $t = 1, ..., 250$) as a training set to select the optimal $\theta$ ($\hat{\theta}$) via LOOCV by using the S-map to forecast species abundances (Cenci *et al.*, 2019). Then, we use $\hat{\theta}$ to fit the S-map over the whole training set and infer $\mathbb{E}(s_i)$ and $|\mathbf{v}_{1i}|$ at the last point in the training set (i.e., $t = 250$) to rank $\langle s_i \rangle$ values (computed via the perturbation analyses). Next, we add a new point to the training set, remove its first point, and repeat the LOOCV and ranking procedures until the end of the time series. Note that we keep the size of the training set fixed after each update (e.g., $t = 2, ..., 251$ for the first update), controlling for the effects of time series length on the performance of the S-map. Also note that we can only infer the coefficients of $\mathbf{J}$ up to a constant (Cenci & Saavedra, 2019), so we only consider the direction of $\mathbf{v}_1$ and the relative value of $\lambda_1$ through time.

Recent improvements of the S-map have been developed to deal with observational and process noise as well as with communities with a large number of species (Cenci *et al.*, 2019, Chang *et al.*, 2021). Here, we find that the classic S-map as described above (Deyle *et al.*, 2016, Sugihara, 1994) already provides a very good inference of expected sensitivities ($\mathbb{E}(s_i)$; Box 1 in the main text) and eigenvector alignments ($|\mathbf{v}_{1i}|$; Box 2 in the main text). In addition to the performance shown in Fig. 3, we show that the classic S-map allows us to accurately predict the order of species sensitivities ($\langle s_i \rangle$) when normalizing species abundances (Fig. S15), when using shorter time series (Fig. S16), when adding observational noise to the time series (Fig. S17), or when the model has a stochastic component (i.e., process noise; Fig. S18). Our analyses with short and noisy time series are described in the next section (see *Section 6*). We believe that combining our ranking approaches with recent developments of the S-map (Cenci *et al.*, 2019, Chang *et al.*, 2021) to deal with large amounts of noise or with communities with a large number of species is an exciting direction for future research.

### 6 Analyses with short and noisy synthetic time series

In our analyses with synthetic time series reported in the main text, we infer the Jacobian matrix ($\mathbf{J}$) and, therefore, expected sensitivities ($\mathbb{E}(s_i)$) and eigenvector alignments ($|\mathbf{v}_{1i}|$) using time series with 250 points and without noise. These conditions, however, are rarely observed in empirical time series, which are typically much shorter and contaminated with noise (Cenci *et al.*, 2019, Sugihara, 1994). In this section, we describe additional analyses with short and noisy synthetic time series.

To test the robustness of our ranking approaches (i.e., using $\mathbb{E}(s_i)$ or $|\mathbf{v}_{1i}|$ to rank $\langle s_i \rangle$ over time) with shorter time series, we perform the S-map using a smaller training set. Instead of using 250 points (e.g., $t = 1, ..., 250$ in the first training set) as described in the previous section, we use only 100 points (e.g., $t = 1, ..., 100$ in the first training set) to train the S-map and infer $\mathbb{E}(s_i)$ and $|\mathbf{v}_{1i}|$ at the last point in the training set to predict species sensitivities ($\langle s_i \rangle$). Fig. S16 shows that our results remain similar to the results in Fig. 3B, which use 250 points.

We also verify the performance of our ranking approaches inferred with the S-map under scenarios of observational noise. To do so, we use the same synthetic time series and perturbation analyses as reported in the main text but add normally distributed noise to the time series used to train the S-map. That is, for each species $i$ and time $t$ in the training set, we transform $N_i(t)$ into $N_i(t) + \mathcal{N}(\mu = 0, \sigma^2 = [\delta N_i(t)]^2)$, where $\delta = 0.1$ (i.e., 10% of observational noise). Then, we use the noisy time series to infer $\mathbb{E}(s_i)$ and $|\mathbf{v}_{1i}|$ with the S-map and predict the order of $\langle s_i \rangle$ at each point in time. The middle column in Fig. S5 shows the attractors for each population dynamics model with observational noise. Fig. S17 shows that, although the mean rank correlation ($\bar{\rho}$) can decrease for some models, our results remain similar to the results in Fig. 3B, which do not contain noise.

We also perform analyses with synthetic time series with process noise. To do so, we generate synthetic time series using a modified version of our population dynamics models (equations (S7), (S8), (S9), and (S10)). In particular, we transform each deterministic population dynamics model $\frac{d\mathbf{N}}{dt} = \mathbf{f}(\mathbf{N})$ into a model with a stochastic component: $d\mathbf{N} = \mathbf{f}(\mathbf{N})dt + \mathbf{g}(\mathbf{N})dW$, where $\mathbf{f}(\mathbf{N})$ is the original deterministic part, $\mathbf{g}(\mathbf{N})$ is the stochastic part, and $W$ is a Wiener process. We use the simplest form of stochasticity, which consists of independent process noise for each species. That is, $\mathbf{g}(\mathbf{N})$ is a diagonal matrix with $N_i\delta$ as the diagonal elements, where $\delta = 0.03$. We then use the stochastic version of the models to generate the synthetic time series but use the deterministic version (i.e., $\delta = 0$) to evolve perturbed points over time in our perturbation analyses (see *Section 4*). Finally, we inferred our ranking approaches with the S-map using the synthetic time series with process noise to predict the order of $\langle s_i \rangle$ over time. The right column in Fig. S5 shows the attractors for each population dynamics model with process noise. Fig. S18 shows that, although the mean rank correlation ($\bar{\rho}$) can decrease for some models, our results remain similar to the results in Fig. 3B.

### 7 Forecast analyses with empirical time series

We apply our ranking approaches to two empirical time series. Both time series contain four interacting variables (hereafter species) and have been shown to exhibit non-equilibrium dynamics for long periods of time (Benincà *et al.*, 2015, 2009). The first time series has 251 points and reports the percentage of cover of barnacles, mussels, crustose algae, and bare rock in a pristine rocky intertidal site sampled monthly for 20 years (Benincà *et al.*, 2015) (Fig. 4A in the main text). The second time series has 794 points and reports the abundance of picocyanobacteria, nanoflagellates, rotifers, and calanoid copepods in an experimental mesocosm sampled twice a week for 7 years (Benincà *et al.*, 2009) (Fig. S21). Because both time series report species abundances on the same scale and unit, we do not normalize species abundances before performing the S-map in order to preserve properties of the Jacobian matrix (e.g., sign of Jacobian coefficients (Song & Saavedra, 2021); but see Fig. S25).

For each time series, we test the hypothesis that the order of species sensitivities ($\mathbb{E}(s_i)$) and species alignments with the leading eigenvector ($|\mathbf{v}_{1i}|$) should predict the order of species standardized forecast errors ($\epsilon_i$; equation (4) in the main text). To do so, we fit the S-map to compute both rankings and use a Long Short-Term Memory (LSTM) neural network (James *et al.*, 2021) to forecast species abundances. Specifically, for each time series, we assign 70% of the data as the training set and sequentially infer the Jacobian matrix with the S-map by moving the training set forward while keeping its size fixed as described in the previous section. In addition, we independently train the LSTM neural network on the training set and forecast the abundances of all species for $\tau = 3$ steps ahead (Cenci *et al.*, 2020). Note that we normalize species abundances to mean zero and unit standard deviation before training the LSTM neural network. Then, we move the training set forward keeping its size fixed, fit the S-map and train the LSTM neural network in the new training set, and forecast abundances for $\tau = 3$ steps ahead until we reach the end of the time series. Thus, for each time $t$ in the test set (i.e., last 30% of points in the time series), we obtain $\mathbb{E}(s_i)$, $|\mathbf{v}_{1i}|$ and $\epsilon_i$ for each species and compute the rank correlation $\rho$ between them. Note that neither the S-map nor the LSTM neural network use information from abundances outside the current training set for inference and forecasting, respectively. Finally, we perform a randomization test to verify whether the mean rank correlation over the test set ($\bar{\rho}$) is significantly greater than zero. For each empirical time series and for each ranking approach, we shuffle $\epsilon_i$ values across species for each point in the test set and compute $\bar{\rho}$ 1,000 times to obtain a $p$-value. We also perform these analyses using $\tau = 2$ (Fig. S22) as well as using 60% and 50% of points in the training set (Figs. S23 and S24).

### 8 Forecast analyses with synthetic time series

In the previous section, we describe our analyses using empirical time series to test the hypothesis that species showing higher forecast errors ($\epsilon_i$) at a given time are also more sensitive to perturbations (i.e., have a higher value of $\mathbb{E}(s_i)$ or $|\mathbf{v}_{1i}|$). Here, we describe similar analyses using the five synthetic time series generated from population dynamics models (see *Section 3*). In these analyses, we compute an average forecast error ($\bar{\epsilon}_i$) for each species by trying to forecast species abundances with the LSTM neural network under perturbations (Cenci *et al.*, 2020, James *et al.*, 2021). First, we separate a given synthetic time series into a training set (first half of the time series) and a test set (second half of the time series). Then, we add 10% of observational noise to the training set (see *Section 6*) and use it to infer the Jacobian matrix with the S-map and to forecast species abundances with the LSTM at the last point in the training set. Following the analyses of the previous section, we forecast species abundances for $\tau = 3$ steps ahead and then move the training set forward by keeping its size fixed and repeat the inference and forecast procedures until the end of the time series. For each time $t$ in the test set, we compute an average forecast root-mean-square error (RMSE) under perturbations for each species $i$ as:

$$\bar{\epsilon}_i = \frac{1}{n}\sum_{j=1}^{n}\sqrt{[\tilde{N}_i^{(j)}(t+\tau-1) - \hat{N}_i(t+\tau-1)]^2}, \tag{S11}$$

where $n$ is the number of perturbed abundances ($n = 300$), $\tilde{N_i}^{(j)}(t+\tau-1)$ is the $j$th perturbed abundance of species $i$ at time $t+\tau-1$, and $\hat{N}_i(t+\tau-1)$ is the forecast of the abundance of species $i$ at time $t+\tau-1$. Thus, we compute the average forecast error of each species for $n$ potential perturbed abundances that could have been observed at a given point in time. Note that perturbed abundances are obtained from our perturbation analyses (see *Section 4*).

We then use the inferred expected sensitivity ($\mathbb{E}(s_i)$) and eigenvector ($|\mathbf{v}_{1i}|$) rankings as well as our alternative indicators (i.e., $\Delta N_i(t)$ or $-N_i(t)$) to predict the order of average forecast errors ($\bar{\epsilon}_i$) over the test set. Note that this analysis follows closely our analyses of predicting the order of species sensitivities ($\langle s_i \rangle$) described in the main text. Fig. S19 shows the results for these analyses as the Spearman's rank correlation ($\rho$) between a given ranking ($\mathbb{E}(s_i)$, $|\mathbf{v}_{1i}|$, $\Delta N_i(t)$, or $-N_i(t)$) and $\bar{\epsilon}_i$ over the test set. The figure shows that, for all models expected for the model with 4 competitor species, $\mathbb{E}(s_i)$ and $|\mathbf{v}_{1i}|$ show, on average, a positive rank correlation with $\bar{\epsilon}_i$ (Fig. S19). Furthermore, the figure shows that this is not the case for $\Delta N_i(t)$ and $-N_i(t)$ (Fig. S19). Therefore, this analysis illustrates that species forecast errors can be related to our measures of sensitivity to perturbations under synthetic time series.

### 9 Leading eigenvector and direction of greatest perturbation expansion under equilibrium dynamics

We now explain how the leading eigenvector of the Jacobian matrix $\mathbf{J}$ (see *Section 1*) points in the direction of greatest expansion of small perturbations under equilibrium dynamics. Under equilibrium dynamics and for sufficiently small perturbations, $\mathbf{J}$ evaluated at the equilibrium $\mathbf{N}^*$ is constant. Thus, we can obtain the general solution of the linear differential equation (S3) as (Boyce *et al.*, 2017, Strogatz, 2018):

$$\mathbf{p}(t+k) = \sum_{i=1}^{S} c_i e^{\lambda_i k}\mathbf{v}_i, \tag{S12}$$

where $\lambda_i$ is the $i$th eigenvalue of $\mathbf{J}$ ($\lambda_S \le \ldots \le \lambda_1$) associated with eigenvector $\mathbf{v}_i$, and the constants $c_i$ depend on the initial condition $\mathbf{p}(t) = \sum_{i=1}^{S} c_i\mathbf{v}_i$. We use $\lambda_i$ and $\mathbf{v}_i$ to denote the real parts of the $i$th eigenvalue and eigenvector, respectively. Under equilibrium dynamics, $\lambda_i < 0$ for all $i$ implies a stable equilibrium, whereas $\lambda_i > 0$ for any $i$ implies an unstable equilibrium

(Strogatz, 2018). Note that, without loss of generality, we can set $t = 0$ for the initial condition. Also note that the solution for $\mathbf{p}(t + k)$ can only be described by equation (S12) if $\mathbf{J}$ has $S$ distinct eigenvalues and, therefore, a set of $S$ linearly independent eigenvectors. We propose that given a sufficient amount of time $k$, $e^{\lambda_1 k}$ will become much larger than subsequent terms (i.e., $e^{\lambda_2 k}, ..., e^{\lambda_S k}$) and, therefore, equation (S12) can be approximated using only the leading eigenvalue and its associated leading eigenvector:

$$\mathbf{p}(t + k) \approx c_1 e^{\lambda_1 k}\mathbf{v}_1. \tag{S13}$$

Therefore, after a sufficient time $k$, perturbed abundances $\mathbf{p}$ will be located closely to the line spanned by $\mathbf{v}_1$.

It is important to note that the time $k$ required for $c_1 e^{\lambda_1 k}\mathbf{v}_1$ to approximate $\mathbf{p}(t+k)$ depends on all eigenvalues and eigenvectors. For example, if $\lambda_S < ... < \lambda_2 < 0 < \lambda_1$ and eigenvectors are orthogonal to each other, then the time $k$ for $c_1 e^{\lambda_1 k}\mathbf{v}_1$ to approximate $\mathbf{p}(t+k)$ is expected to be be small (see first scenario in *Section 13* and Fig. S1). Importantly, this is the scenario we expect to observe in chaotic non-equilibrium dynamical systems that typically have directions of expansion (i.e., unstable manifold) and contraction (i.e., stable manifold) at each point along an attractor (Eckmann & Ruelle, 1985, Strogatz, 2018). On the other hand, if more than one eigenvalue is positive or if eigenvectors are not orthogonal, then the time $k$ for $c_1 e^{\lambda_1 k}\mathbf{v}_1$ to approximate $\mathbf{p}(t+k)$ is expected to be large (see second scenario in *Section 13* and Fig. S2).

In addition, it is also important to consider the case of complex eigenvalues and eigenvectors. In this case, the real solution approximated using only $\lambda_1$ and $\mathbf{v}_1$ is given by (Boyce *et al.*, 2017):

$$\mathbf{p}(t + k) \approx c_1\mathbf{p}_1 + c_2\mathbf{p}_2, \tag{S14}$$

where $c_1$ and $c_2$ are constants and $\mathbf{p}_1$ and $\mathbf{p}_2$ are the two linearly independent real solutions given by:

$$\begin{aligned} \mathbf{p}_1 &= e^{ak}[\mathbf{u}\cos(bk) - \mathbf{z}\sin(bk)] \\ \mathbf{p}_2 &= e^{ak}[\mathbf{u}\sin(bk) + \mathbf{z}\cos(bk)], \end{aligned} \tag{S15}$$

where $\lambda_1 = a + ib$, $\lambda_2 = a - ib$ is the pair of leading complex eigenvalues and $\mathbf{v}_1 = \mathbf{u} + i\mathbf{z}$, $\mathbf{v}_2 = \mathbf{u} - i\mathbf{z}$ is the pair of leading complex eigenvectors. Thus, in this case the solution $\mathbf{p}(t + k)$ is oscillatory. However, we can see that if the imaginary parts of the leading eigenvalue and leading eigenvector ($b$ and $\mathbf{z}$) are not too strong, then their real parts ($a$ and $\mathbf{u}$) still inform us about the magnitude and direction of greatest expansion of perturbations, respectively (see third scenario in *Section 13* and Fig. S3). Finally, we note that $b$ (and therefore $\mathbf{z}$) is zero for the majority of points in three out of five synthetic time series that we analyze (predator-prey (2 sp): 47.7%; food chain (3 sp): 69.1%; food web (3 sp): 81.6%; competitors (4 sp): 26.4%; and food web (5 sp): 95.2%). To keep a simple notation, we use $\lambda_i$ and $\mathbf{v}_i$ throughout the text to refer to the real part of the $i$th eigenvalue and eigenvector, respectively.

#### 10 Leading Lyapunov vector and direction of greatest perturbation expansion under non-equilibrium dynamics

In this study, we focus on non-equilibrium attractors such as limit cycles or chaotic attractors (Fig. S5). By "non-equilibrium dynamics" we refer to trajectories of a deterministic dynamical system (e.g., population dynamics model) that do not settle to an equilibrium point. A large literature on nonlinear dynamics has shown that local Lyapunov exponents and their associated Lyapunov vectors determine how a (hyper)sphere of small perturbations at a given state $\mathbf{N}$ deforms into a (hyper)ellipsoid after sufficient time (Eckmann & Ruelle, 1985, Kuptsov & Parlitz, 2012, Mease *et al.*, 2003, Strogatz, 2018, Vallejo *et al.*, 2017). Let $l_i$ ($l_S \leq ... \leq l_1$) and $\mathbf{w}_i$ denote the $i$th local Lyapunov exponent and vector, respectively. If at time $t$ we apply $S$ perturbations with a small norm $||\mathbf{p}_i(t)|| = \delta$ ($i = 1, ..., S$) in the directions of $\mathbf{w}_i$ (i.e., $\frac{\mathbf{p}_i(t)}{\delta} = \mathbf{w}_i$), then after some time $k$, $||\mathbf{p}_i(t+k)|| \approx ||\mathbf{p}_i(t)||e^{l_i k}$ denotes the length of the $i$th principal axis of the ellipsoid (Kuptsov & Parlitz, 2012, Mease *et al.*, 2003, Strogatz, 2018, Vallejo *et al.*, 2017). As we have mentioned in *Section 1*, under non-equilibrium dynamics small perturbations evolve according to $\frac{d\mathbf{p}}{dt} = \mathbf{J}\mathbf{p}$. If $\mathbf{J}$ is constant through time, as is the case when it is evaluated at an equilibrium point, it has been shown that Lyapunov vectors ($\mathbf{w}_1, ..., \mathbf{w}_S$) are equivalent to the eigenvectors of $\mathbf{J}$ ($\mathbf{v}_1, ..., \mathbf{v}_S$) and Lyapunov exponents ($l_S \leq ... \leq l_1$) are equivalent to the eigenvalues of $\mathbf{J}$ ($\lambda_S \leq ... \leq \lambda_1$) (Kuptsov & Parlitz, 2012, Mease *et al.*, 2003). Nevertheless, when $\mathbf{J}$ is not constant through time, it is necessary to incorporate information on all $\mathbf{J}$ matrices along a trajectory to estimate $l_i$ and $\mathbf{w}_i$. The problem with this approach, however, is that it requires information beyond time $t$ in order to detect the directions of perturbation expansion/contraction at time $t$ and therefore is not useful for real-world applications. Thus, the question is whether the leading eigenvector can be used as a proxy for the leading Lyapunov vector to detect the direction of greatest expansion of small perturbations under non-equilibrium dynamics.

Here we specify the conditions under which the leading eigenvector $\mathbf{v}_1$ is a good approximation to the leading Lyapunov vector $\mathbf{w}_1$. On the one hand, we hypothesize that when the rate of change of the system ($\frac{d\mathbf{N}}{dt}$) is large, the Jacobian matrix $\mathbf{J}$ changes rapidly and $\mathbf{v}_1$ approximates $\mathbf{w}_1$ only for a small time $k$. Note that, under these circumstances, only a small amount of time is required for $c_1 e^{\lambda_1 k}\mathbf{v}_1$ to approximate $\mathbf{p}(t+k)$ (equation (S13)). On the other hand, we hypothesize that when $\frac{d\mathbf{N}}{dt}$ is small, $\mathbf{J}$ changes slowly and $\mathbf{v}_1$ approximates $\mathbf{w}_1$ for a larger time $k$. Note that, under this scenario, a larger amount of time is required for $c_1 e^{\lambda_1 k}\mathbf{v}_1$ to approximate $\mathbf{p}(t+k)$. Therefore, the leading eigenvector must show a higher accuracy in detecting the direction of greatest perturbation expansion when the amount of time $k$ for which perturbations evolve is inversely proportional to the current rate of change of the system. Note that we set $k$ to be inversely proportional to the rate of change of the system in our main perturbation analyses (see *Section 4*; Fig. 3 in the main text), but also perform perturbation analyses using fixed values of

$k$ (see *Section 4*; Figs. S11 and S12).

To verify how well the leading eigenvector $\mathbf{v}_1$ approximates the leading Lyapunov vector $\mathbf{w}_1$, we compute $\mathbf{w}_1$ for all points in each time series generated by each of the five population dynamics models used in this study (see *Section 3*). Although computing the complete set of Lyapunov vectors is a more complicated procedure (Ginelli *et al.*, 2007, Kuptsov & Parlitz, 2012), computing just $\mathbf{w}_1$ (i.e., direction of greatest perturbation expansion) is straightforward (Vallejo *et al.*, 2017). Specifically, we compute $\mathbf{w}_1$ by applying a small perturbation $\mathbf{p}$ at time $t$ and evolving the original dynamics ($\frac{d\mathbf{N}}{dt} = \mathbf{f}(\mathbf{N})$) and the tangent dynamics ($\frac{d\mathbf{p}}{dt} = \mathbf{J}\mathbf{p}$) simultaneously for $k$ time steps. Then, $\mathbf{p}$ will rotate over time to the direction of $\mathbf{w}_1$ while expanding at a rate given by the leading Lyapunov exponent ($l_1$) (Kuptsov & Parlitz, 2012, Mease *et al.*, 2003, Vallejo *et al.*, 2017). For the convergence of $\mathbf{p}$ to $\mathbf{w}_1$ to be faster, we follow standard methods (Vallejo *et al.*, 2017) and choose $\mathbf{p}$ to be a vector with a small norm $r$ in the direction of $\mathbf{v}_1$. Specifically $\mathbf{p} = r\frac{\mathbf{v}_1}{||\mathbf{v}_1||}$, where we set $r$ to be 5% of the mean standard deviation of species abundances: $r = 0.05\frac{1}{S}\sum_{i=1}^{S}\sigma_{N_i}$. For each point in time, we use the same value of $k$ as used in our perturbation analyses (i.e., $k$ is inversely proportional to the local rate of change of the dynamics as described in *Section 4*. The leading Lyapunov vector at time $t$ can then be estimated as $\mathbf{w}_1 = \mathbf{p}(t+k)$, whereas the leading Lyapunov exponent can be calculated as $l_1 = \frac{1}{k}\log\left(\frac{||\mathbf{p}(t+k)||}{||\mathbf{p}(t)||}\right)$. To verify how aligned $\mathbf{v}_1$ is with $\mathbf{w}_1$, we compute the absolute value of the cosine of the angle between $\mathbf{v}_1$ and $\mathbf{w}_1$ at each point in time. Thus, if $\mathbf{v}_1$ indeed points in the direction of $\mathbf{w}_1$, we expect that only the magnitude and not the direction of $\mathbf{p}$ will change after $k$ time steps. In this case, the growth rate of the magnitude of $\mathbf{p}$ is given by $l_1$. To benchmark the observed alignment between $\mathbf{v}_1$ and $\mathbf{w}_1$, we repeat the procedure above but choose $\mathbf{p}$ to be a vector with norm $r$ and a random direction at each point in time. We use this analysis to compare the alignment between $\mathbf{v}_1$ and $\mathbf{w}_1$ (expected to be high) with the alignment of a randomly chosen vector $\mathbf{p}(t)$ and $\mathbf{p}(t+k)$ (expected to be low). We find $\mathbf{v}_1$ to be highly aligned with $\mathbf{w}_1$ (i.e., absolute value of cosine close to 1) for all five synthetic time series (left boxplots in Fig. S4). In contrast, when the initial perturbation ($\mathbf{p}(t)$) has a random direction instead of the direction of $\mathbf{v}_1$, we find it to be poorly aligned with $\mathbf{p}(t+k)$ (right boxplots in Fig. S4).

### 11 From direction of greatest perturbation expansion to ranking species sensitivities

Now, we show how we can rank species sensitivities based on the direction of greatest perturbation expansion approximated by the leading eigenvector. We define the sensitivity of species $i$ to a single perturbation $\mathbf{p}$ from time $t$ to $t+k$ as the squared difference between its perturbed and unperturbed abundance in relation to the initial squared difference caused by the perturbation (equation (1) in the main text):

$$s_i = \frac{[\tilde{N}_i(t+k) - N_i(t+k)]^2}{[\tilde{N}_i(t) - N_i(t)]^2} = \frac{p_i(t+k)^2}{p_i(t)^2}. \tag{S16}$$

Note that under equilibrium dynamics, we can just change $N_i$ to $N_i^*$ and the derivation below remains the same. Let us first consider the numerator of the equation above by substituting the approximated solution of the linearized dynamics (equation (S13)) into it:

$$\begin{aligned} p_i(t+k)^2 &\approx [c_1 e^{\lambda_1 k}\mathbf{v}_{1i}]^2 \\ &\approx c_1^2 e^{2\lambda_1 k}\mathbf{v}_{1i}^2, \end{aligned} \tag{S17}$$

where $\mathbf{v}_{1i}^2$ corresponds to the square of the $i$th element of $\mathbf{v}_1$. Thus, $c_1^2 e^{2\lambda_1 k}$ represents the total amount of expansion, which depends on $\lambda_1$, $k$, and $c_1$ via the initial condition. Note, however, that this term is the same for every species $i$. Therefore, the values of $p_i(t+k)^2$ across species can be ranked using $|\mathbf{v}_{1i}|$, which follows the same order as $\mathbf{v}_{1i}^2$. We use $|\mathbf{v}_{1i}|$ instead of $\mathbf{v}_{1i}^2$ because it has a clear geometric interpretation as the alignment of $\mathbf{v}_1$ with the coordinate axis corresponding to species $i$ in state space. That is, if $||\mathbf{v}_1|| = 1$, then $|\mathbf{v}_{1i}|$ is equivalent to the absolute value of the cosine of the angle $\alpha_i$ between $\mathbf{v}_1$ and $\mathbf{e}_i$: $|\mathbf{v}_{1i}| = |\cos\alpha_i| = |\mathbf{v}_1\mathbf{e}_i|$, where $\mathbf{e}_i$ is the $i$th standard basis vector.

So far, we have only considered species sensitivities to a single perturbation $\mathbf{p}$. We now consider multiple perturbations at time $t$ ($\mathbf{p}(t)$), which follow a given distribution with mean zero and covariance matrix $\mathbf{\Sigma}_t$. For a set of $n$ randomly perturbed abundances, we can define the sensitivity of species $i$ from time $t$ to $t+k$ as the average squared difference between a set of $n$ randomly perturbed abundances and its unperturbed abundance in relation to the initial average squared difference (equation (2) in the main text):

$$\langle s_i \rangle = \frac{\frac{1}{n}\sum_{j=1}^{n}[\tilde{N}_i^{(j)}(t+k) - N_i(t+k)]^2}{\frac{1}{n}\sum_{j=1}^{n}[\tilde{N}_i^{(j)}(t) - N_i(t)]^2} = \frac{\frac{1}{n}\sum_{j=1}^{n} p_i^{(j)}(t+k)^2}{\frac{1}{n}\sum_{j=1}^{n} p_i^{(j)}(t)^2}. \tag{S18}$$

By focusing on the numerator, we can see that $\frac{1}{n}\sum_{j=1}^{n} p_i^{(j)}(t+k)^2 = \mathbb{E}[c_1^2 e^{2\lambda_1 k}\mathbf{v}_{1i}^2] = e^{2\lambda_1 k}\mathbf{v}_{1i}^2\mathbb{E}[c_1^2]$, since $e^{2\lambda_1 k}$ and $\mathbf{v}_{1i}^2$ are constants. The expectation $\mathbb{E}[c_1^2]$ will depend on the distribution of initial conditions, but will affect the sensitivity of all species by the same amount. Finally, we note that because $p_i(t)$ has mean zero, the denominator of equation (S18) is a constant given by the variance of $p_i(t)$ (i.e., the $i$th diagonal element $\sigma_{i,t}^2$ of $\mathbf{\Sigma}_t$; *Section 2*). Thus, if $\sigma_{i,t}^2$ is the same for every species $i$, the denominator of equation (S18) will not affect the order of $\langle s_i \rangle$ values and we can use $|\mathbf{v}_{1i}|$ to rank species sensitivities. However, we keep this denominator in our definition of $\langle s_i \rangle$ to control for distinct variances across species in one of our perturbation analyses (see *Section 4*).

### 12 Connection between expected sensitivity and eigenvector approaches

Here, we show a connection between our measures of expected sensitivity ($\mathbb{E}(s_i)$; Box 1 in the main text) and alignment with the leading eigenvector ($|\mathbf{v}_{1i}|$; Box 2 in the main text) under two simplifying assumptions. First, we assume that all species are affected by perturbations with the same variance and there is no covariance among species pairs (i.e., the covariance matrix of perturbations $\mathbf{\Sigma}_t$ is the identity matrix $\mathbf{I}$). Second, we assume that the Jacobian matrix $\mathbf{J}$ at time $t$ is symmetric (i.e., $\mathbf{J} = \mathbf{J}^\top$). Although these assumptions may not be fulfilled in natural communities, they allow us to obtain a first insight into the connections between $\mathbb{E}(s_i)$ and $|\mathbf{v}_{1i}|$. Using these assumptions, we can write the following equation for the covariance matrix of perturbations at time $t+k$ (*Section 2*):

$$\begin{aligned}
\mathbf{\Sigma}_{t+k} &= e^{\mathbf{J}k}\mathbf{\Sigma}_t e^{\mathbf{J}^\top k} \\
&= e^{\mathbf{J}k} e^{\mathbf{J}k} \\
&= e^{\mathbf{J}k+\mathbf{J}k} \\
&= e^{\mathbf{J}2k},
\end{aligned} \tag{S19}$$

where $e^{\mathbf{A}}$ is the exponential of a given matrix $\mathbf{A}$ and we have used the fact that if $\mathbf{A}$ and $\mathbf{B}$ commute then $e^{\mathbf{A}}e^{\mathbf{B}} = e^{\mathbf{A}+\mathbf{B}}$. Now, we can write the eigendecomposition of $\mathbf{\Sigma}_{t+k}$ as:

$$\mathbf{\Sigma}_{t+k} = \mathbf{V} e^{\mathbf{\Lambda}2k} \mathbf{V}^\top, \tag{S20}$$

where $\mathbf{V}$ is the matrix containing the eigenvectors of $\mathbf{J}$ ($\mathbf{v}_i$) as column vectors and $\mathbf{\Lambda}$ is the diagonal matrix containing the eigenvalues of $\mathbf{J}$ ($\lambda_i$). Note that we have used the property that $\mathbf{A}$ and $e^{\mathbf{A}}$ share the same eigenvectors and that if $\lambda_i$ is an eigenvalue of $\mathbf{A}$, then $e^{\lambda_i}$ is the corresponding eigenvalue of $e^{\mathbf{A}}$. The expected sensitivity of species $i$ is defined as the $i$th diagonal element of $\mathbf{\Sigma}_{t+k}$ ($\sigma^2_{i,t+k}$; *Section 2*), which gives us:

$$\begin{aligned}
\mathbb{E}(s_i) &= \sigma^2_{i,t+k} \\
&= \sum_{j=1}^{S} \mathbf{v}^2_{ji} e^{\lambda_j 2k} \\
&\approx \mathbf{v}^2_{1i} e^{\lambda_1 2k},
\end{aligned} \tag{S21}$$

where $\mathbf{v}_{ji}$ is the $j$th element of $\mathbf{v}_i$ and in the last step we used the fact that, given a sufficient amount of time $k$, $e^{\lambda_1 2k}$ will become much larger than $e^{\lambda_2 2k}, ..., e^{\lambda_S 2k}$ and will dominate the expression. Thus, the order of $\mathbb{E}(s_i)$ values will follow closely the order of $|\mathbf{v}_{1i}|$ values under the assumptions considered here. Finally, note that the final expression in equation (S21) is very similar to what we obtained in equation (S17) as an explanation of how we can use $|\mathbf{v}_{1i}|$ to rank species sensitivities to a given perturbation ($s_i$).

### 13 Illustrations with Lotka-Volterra dynamics at equilibrium

To illustrate how expected sensitivities ($\mathbb{E}(s_i)$; Box 1 in the main text) and alignments with the leading eigenvector ($|\mathbf{v}_{1i}|$; Box 2 in the main text) are able to rank species according to their sensitivity to perturbations ($\langle s_i \rangle$), we use the classic Lotka-Volterra model (equation (S9)) under equilibrium dynamics. For this model, the vector of species abundances at equilibrium is given by: $\mathbf{N}^* = -\mathbf{A}^{-1}\mathbf{r}$. While the focus of our study is on non-equilibrium dynamics, our goal here is simply to show the performance of these two proposed methods under three simple scenarios of equilibrium dynamics. Our results for non-equilibrium dynamics are described in the main text.

We use three different scenarios of the Lotka-Volterra dynamics with $S = 3$ species. For all scenarios we choose a combination of $\mathbf{r}$ and $\mathbf{A}$ giving the following feasible (i.e., positive abundances for all species) equilibrium: $\mathbf{N}^* = [1, 1, 1]^\top$. Note that for this feasible equilibrium, the Jacobian matrix evaluated at $\mathbf{N}^*$ is given by: $\mathbf{J} = \text{diag}(\mathbf{N}^*)\mathbf{A} = \mathbf{A}$. For each scenario, we compute the eigenvalues ($\lambda_i$) and eigenvectors ($\mathbf{v}_i$) of $\mathbf{J}$ as well as expected sensitivities ($\mathbb{E}(s_i)$) using $k = 0.1$, 0.2, 0.3, 0.4, and 0.5. We then perform 2,000 normally distributed perturbations $\mathbf{p}$ to $\mathbf{N}^*$ (i.e., $p_i \sim \mathcal{N}(\mu = 0, \sigma^2 = r^2)$ with $r = 0.05$) and evolve each perturbed abundance over time according to equation (S9) for $k = 0.5$ time steps. Finally, we compute species sensitivities ($\langle s_i \rangle$) at $t = 0.1$, 0.2, 0.3, 0.4, and 0.5 using all perturbed abundances at those time points.

The first scenario (Fig. S1) consists of the following parameter values of the Lotka-Volterra model:

$$\mathbf{r} = \begin{bmatrix} 1 \\ 1 \\ 1 \end{bmatrix}, \mathbf{A} = \begin{bmatrix} 1 & -2 & 0 \\ 0 & -1 & 0 \\ 0 & 2 & -3 \end{bmatrix}$$

The eigenvalues of $\mathbf{J}$ show that the feasible equilibrium for this system is a saddle point: $\lambda_1 = 1$ (unstable manifold), $\lambda_2 = -1$, and $\lambda_3 = -3$ (stable manifolds). The order of expected sensitivities is given by $\mathbb{E}(s_3) < \mathbb{E}(s_2) < \mathbb{E}(s_1)$, which corresponds exactly to the order of species sensitivities ($\langle s_i \rangle$) for all times (Fig. S1B, C). The order of eigenvector alignments is given by $|\mathbf{v}_{13}|, |\mathbf{v}_{12}| < |\mathbf{v}_{11}|$ and corresponds closely to the order of species sensitivities, but cannot distinguish species 2 and 3 (Fig. S1B, C). Note that expected sensitivities depend on the time step $k$, whereas eigenvector alignments do not.

The second scenario (Fig. S2) consists of the following parameter values:

$$\mathbf{r} = \begin{bmatrix} -4.5 \\ 17.5 \\ 7 \end{bmatrix}, \mathbf{A} = \begin{bmatrix} 4 & 0.5 & 0 \\ 0.5 & -10 & -8 \\ 0 & -8 & 1 \end{bmatrix}$$

The eigenvalues of $\mathbf{J}$ show that the feasible equilibrium is again a saddle point: $\lambda_1 = 5.2$, $\lambda_2 = 4.0$ (unstable manifolds), and $\lambda_3 = -14.2$ (stable manifold). However, this scenario is more challenging than the previous one for our ranking approaches because there are two (instead of one) directions of perturbation expansion. The order of expected sensitivities is given by $\mathbb{E}(s_2) < \mathbb{E}(s_1) < \mathbb{E}(s_3)$, which corresponds exactly to the order of species sensitivities from $k = 0.2$ to $k = 0.5$ (Fig. S2B, C). The order of eigenvector alignments is given by $|\mathbf{v}_{11}| < |\mathbf{v}_{12}| < |\mathbf{v}_{13}|$ and provides a reasonable match to the order of species sensitivities (Fig. S2B, C).

Finally, the third scenario (Fig. S3) consists of the following parameter values:

$$\mathbf{r} = \begin{bmatrix} 5 \\ -1 \\ -7 \end{bmatrix}, \mathbf{A} = \begin{bmatrix} -4 & -3 & 2 \\ -2 & 1 & 2 \\ 5 & 2 & 0 \end{bmatrix}$$

For this scenario, the leading eigenvalue of $\mathbf{J}$ is complex and therefore indicate oscillatory dynamics: $\lambda_1 = 2.0 + 0.7i$, $\lambda_2 = 2.0 - 0.7i$, and $\lambda_3 = -7.0 + 0i$. This scenario is also challenging for our ranking approaches due to this oscillatory behavior. Note, however, that the imaginary part of the leading eigenvalue is small compared to the real part. The order of expected sensitivities is given by $\mathbb{E}(s_1) < \mathbb{E}(s_3) < \mathbb{E}(s_2)$, which corresponds exactly to the order of species sensitivities from $k = 0.3$ to $k = 0.5$ (Fig. S3B, C). The order of eigenvector alignments is given by $|\mathbf{v}_{13}| < |\mathbf{v}_{11}| < |\mathbf{v}_{12}|$ and provides a reasonable match to the order of species sensitivities (Fig. S3B, C).

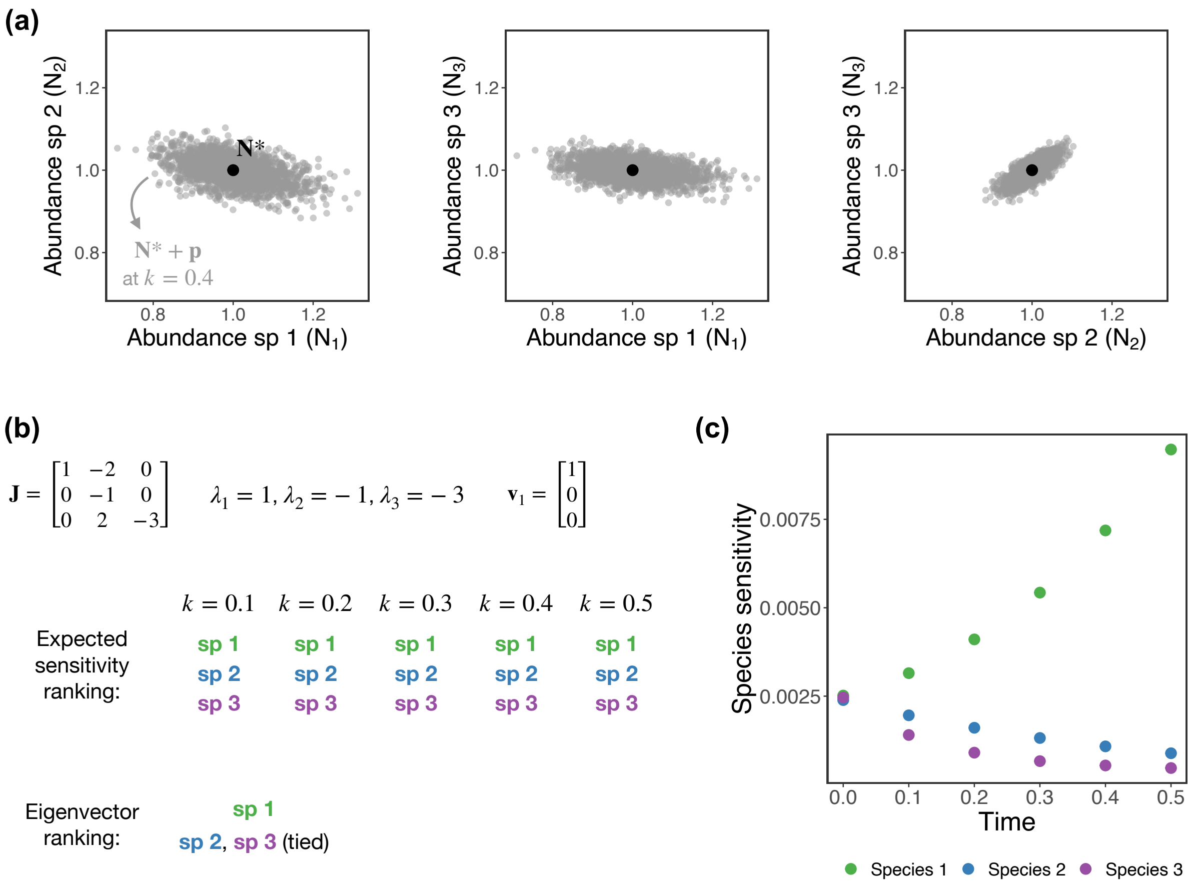


**Figure S1.** First scenario of Lotka-Volterra dynamics at equilibrium (see *Section 13*) showing how expected sensitivities ($\mathbb{E}(s_i)$) and alignments with the leading eigenvector ($|\mathbf{v}_{1i}|$) can rank species sensitivities to perturbations ($\langle s_i \rangle$). (a) Perturbed abundances ($\tilde{\mathbf{N}} = \mathbf{N}^* + \mathbf{p}$; 2,000 gray points) at time $k = 0.4$ projected onto the planes of species 1 and 2 (left), species 1 and 3 (middle), and species 2 and 3 (right). (b) Jacobian matrix ($\mathbf{J}$) and its eigenvalues ($\lambda_i$) and leading eigenvector ($\mathbf{v}_1$) for this Lotka-Volterra system (top). The order of expected sensitivities (computed using different values of $k$) and eigenvector alignments (bottom). (c) Species sensitivities computed using the perturbed abundances (gray points in (a)) at different points in time (i.e., for different values of $k$). In this scenario, the expected sensitivity ranking is more accurate than the eigenvector ranking.

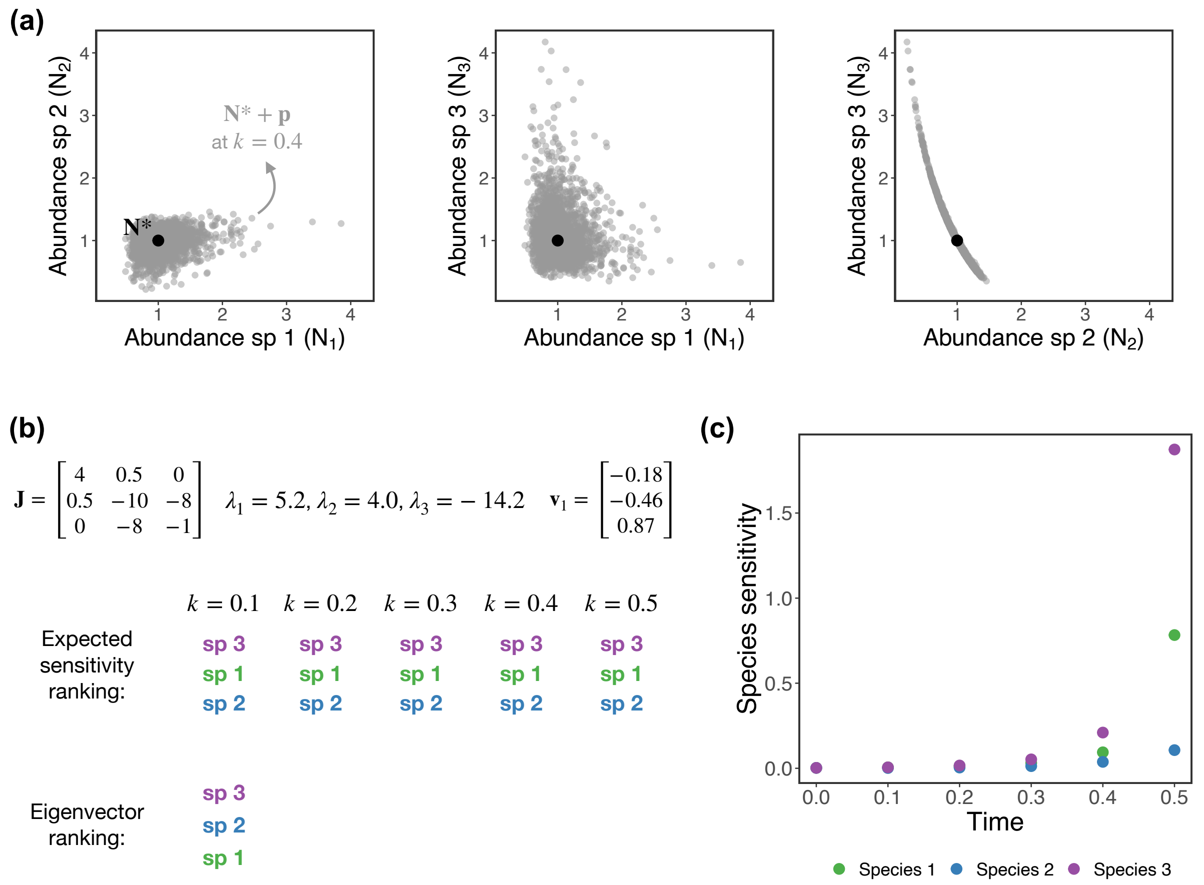


**Figure S2.** Second scenario of Lotka-Volterra dynamics at equilibrium (see *Section 13*) showing how expected sensitivities ($\mathbb{E}(s_i)$) and alignments with the leading eigenvector ($|\mathbf{v}_{1i}|$) can rank species sensitivities to perturbations ($\langle s_i \rangle$). (a) Perturbed abundances ($\tilde{\mathbf{N}} = \mathbf{N}^* + \mathbf{p}$; 2,000 gray points) at time $k = 0.4$ projected onto the planes of species 1 and 2 (left), species 1 and 3 (middle), and species 2 and 3 (right). (b) Jacobian matrix ($\mathbf{J}$) and its eigenvalues ($\lambda_i$) and leading eigenvector ($\mathbf{v}_1$) for this Lotka-Volterra system (top). The order of expected sensitivities (computed using different values of $k$) and eigenvector alignments (bottom). (c) Species sensitivities computed using the perturbed abundances (gray points in (a)) at different points in time (i.e., for different values of $k$). In this scenario, the expected sensitivity ranking is more accurate than the eigenvector ranking.

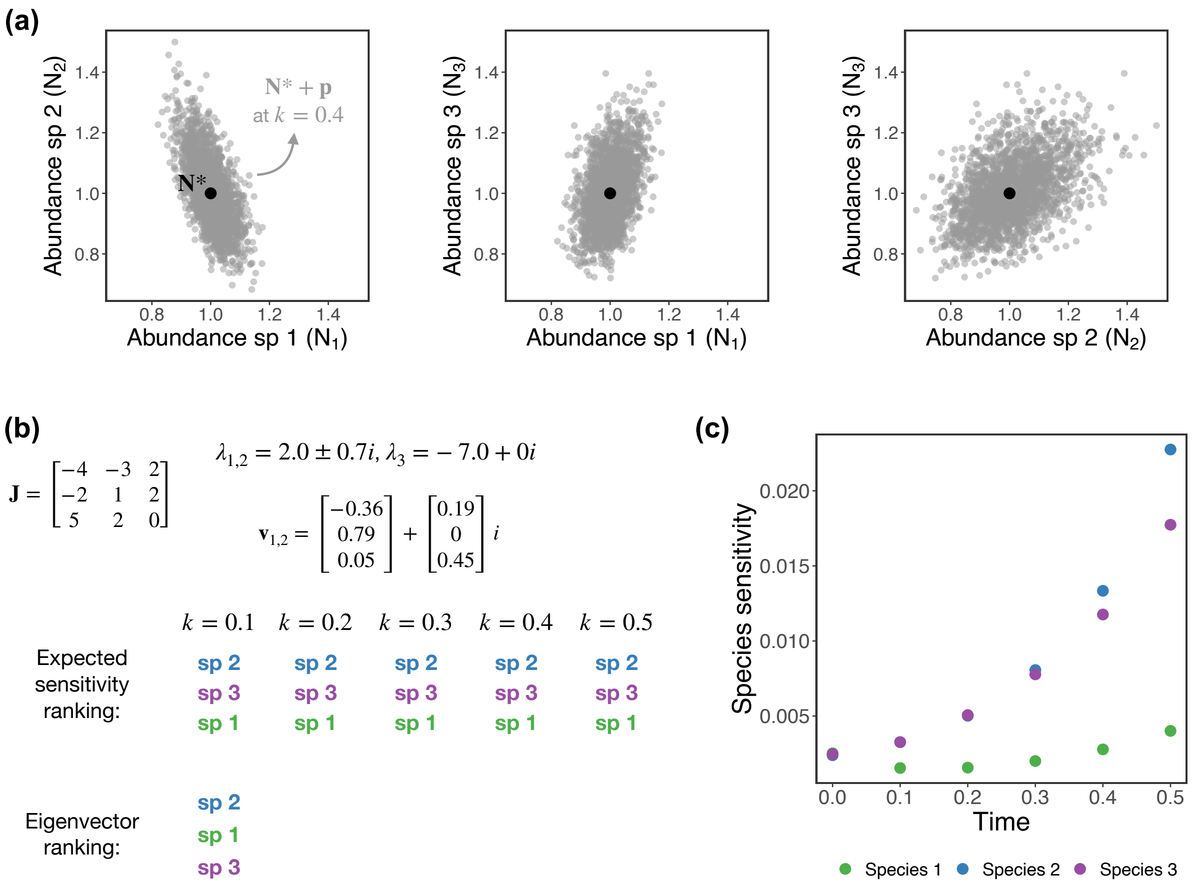


**Figure S3.** Third scenario of Lotka-Volterra dynamics at equilibrium (see *Section 13*) showing how expected sensitivities ($\mathbb{E}(s_i)$) and alignments with the leading eigenvector ($|\mathbf{v}_{1i}|$) can rank species sensitivities to perturbations ($\langle s_i \rangle$). (a) Perturbed abundances ($\tilde{\mathbf{N}} = \mathbf{N}^* + \mathbf{p}$; 2,000 gray points) at time $k = 0.4$ projected onto the planes of species 1 and 2 (left), species 1 and 3 (middle), and species 2 and 3 (right). (b) Jacobian matrix ($\mathbf{J}$) and its eigenvalues ($\lambda_i$) and leading eigenvector ($\mathbf{v}_1$) for this Lotka-Volterra system (top). The order of expected sensitivities (computed using different values of $k$) and eigenvector alignments (bottom). (c) Species sensitivities computed using the perturbed abundances (gray points in (a)) at different points in time (i.e., for different values of $k$). In this scenario, the expected sensitivity ranking is more accurate than the eigenvector ranking.

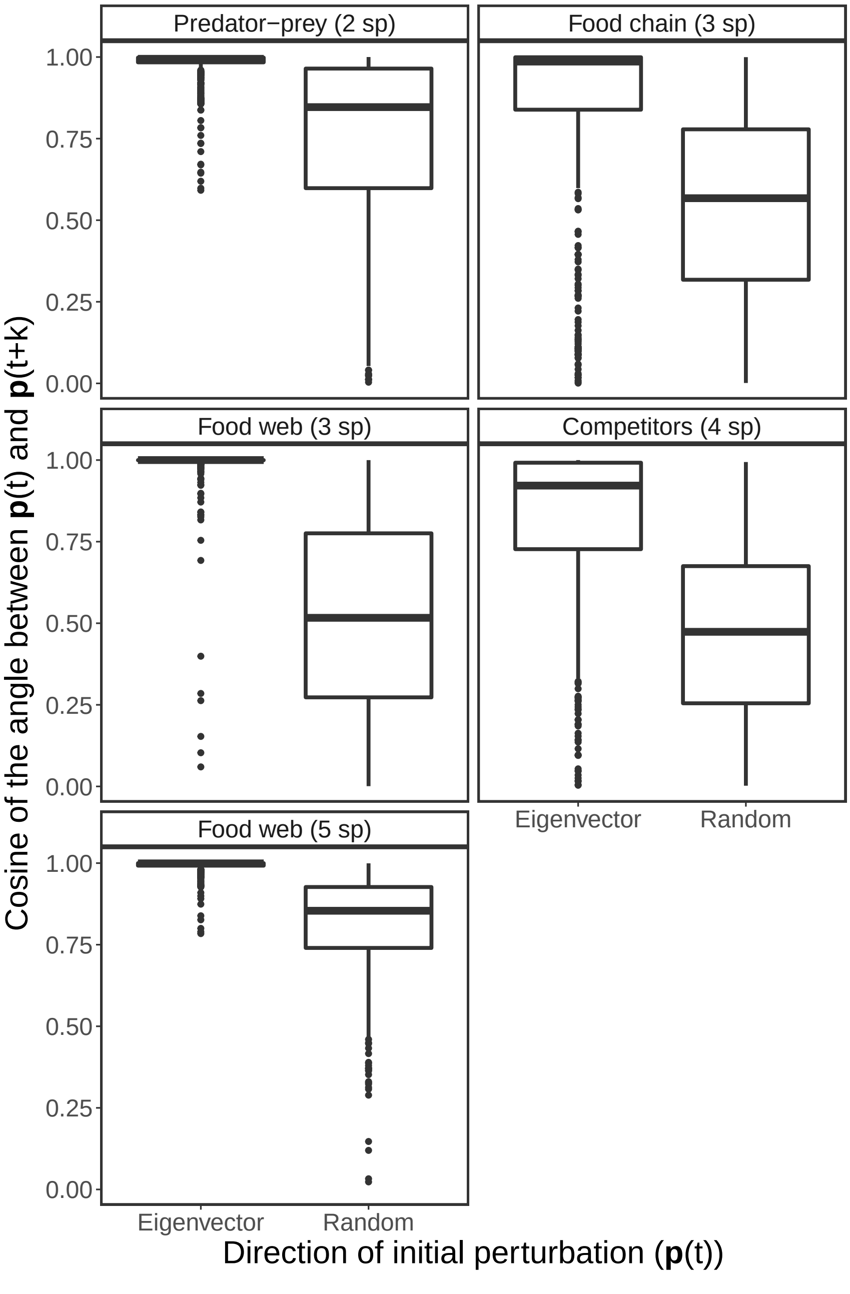


**Figure S4.** Alignments (i.e., absolute value of cosine of the angle) between the initial ($\mathbf{p}(t)$) and final ($\mathbf{p}(t+k)$) perturbation vector for two directions of $\mathbf{p}(t)$ for the five population dynamics models (see *Section 10*). Boxplots on the left correspond to $\mathbf{p}(t)$ in the direction of the leading eignevector ($\mathbf{v}_1$) whereas boxplots on the right correspond to $\mathbf{p}(t)$ in a random direction. Note that $\mathbf{p}(t+k)$ converges to the leading Lyapunov vector ($\mathbf{w}_1$) when $\mathbf{p}(t)$ is in the direction of $\mathbf{v}_1$. The figure shows that $\mathbf{v}_1$ is on average much more aligned with $\mathbf{w}_1$ (left boxplots) than what is expected at random (right boxplots).

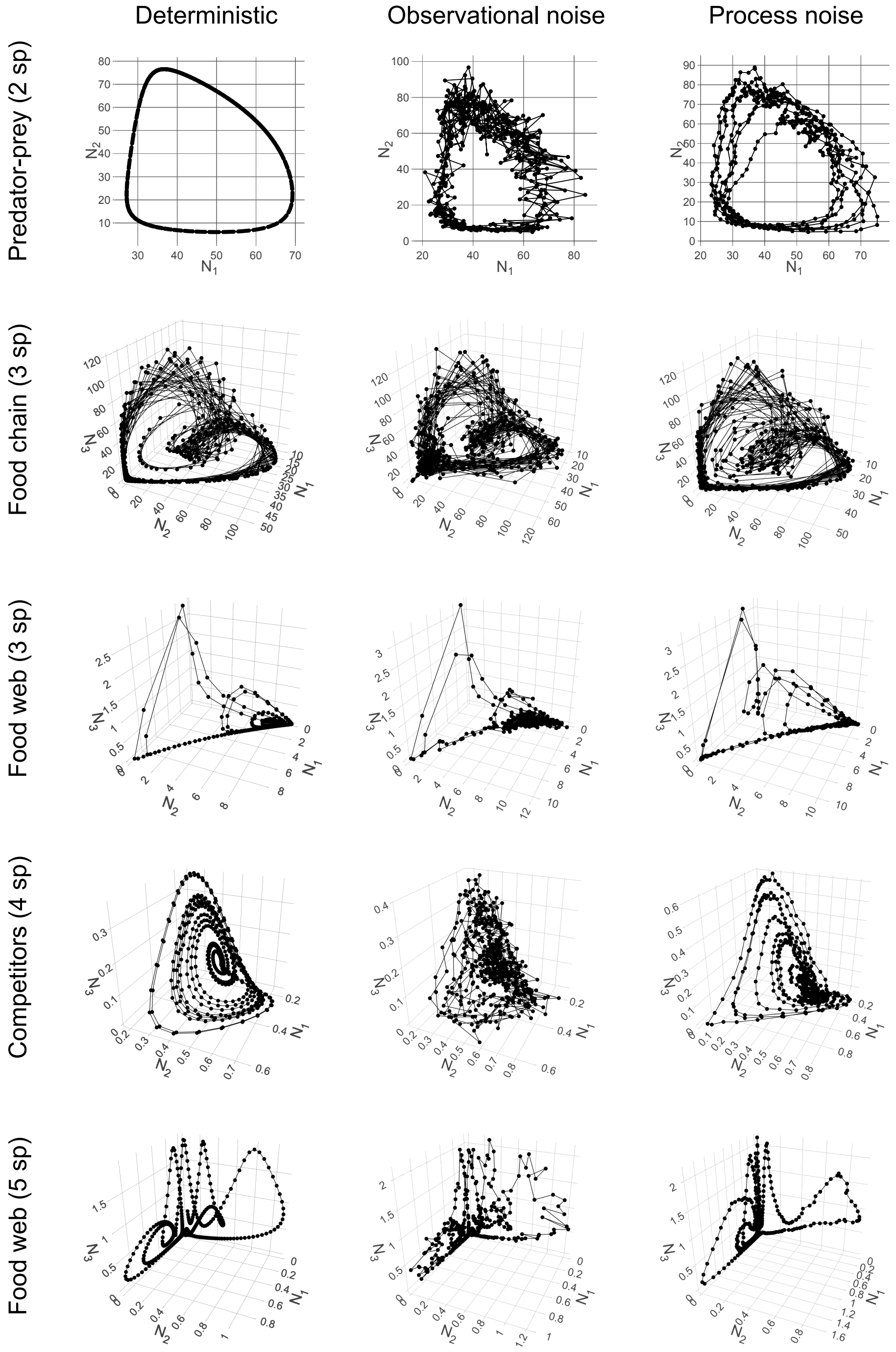


**Figure S5.** Attractors in state space corresponding to each multivariate synthetic time series generated from a population dynamics model (different rows; see *Section 3*) with a different type of noise (different columns; see *Section 6*). Each plot shows the 500 points ($\{\mathbf{N}(t)\}$, $t = 1, ..., 500$) generated by numerically integrating the indicated model and then sampling equidistant points. Note that we only show the abundances of species 1, 2, and 3 for models with more than 3 species.

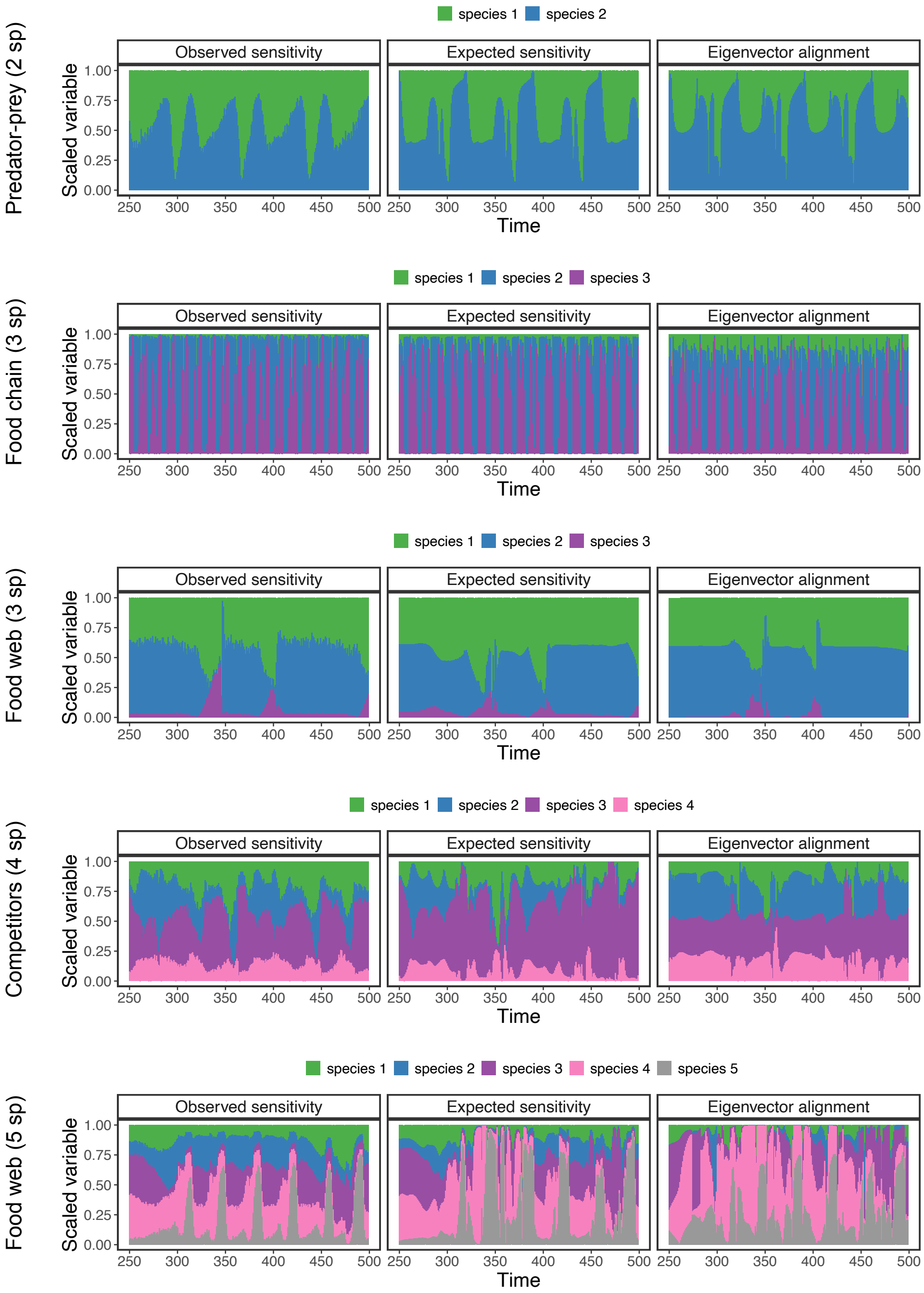


**Figure S6.** Species sensitivities computed from our perturbation analyses ($\langle s_i \rangle$; first column) as well as expected sensitivities ($\mathbb{E}(s_i)$; second column) and eigenvector alignments ($|\mathbf{v}_{1i}|$; third column) inferred from each synthetic time series (different rows) with the S-map over time. A bar in one of the plots shows the values of the corresponding variable (i.e., $\langle s_i \rangle$, $\mathbb{E}(s_i)$, or $|\mathbf{v}_{1i}|$) across species. Note that variables are rescaled to sum 1 across species to improve visualization but that this procedure does not change the rankings. These results correspond to our main set of analyses with synthetic time series shown in the main text (Fig. 3).

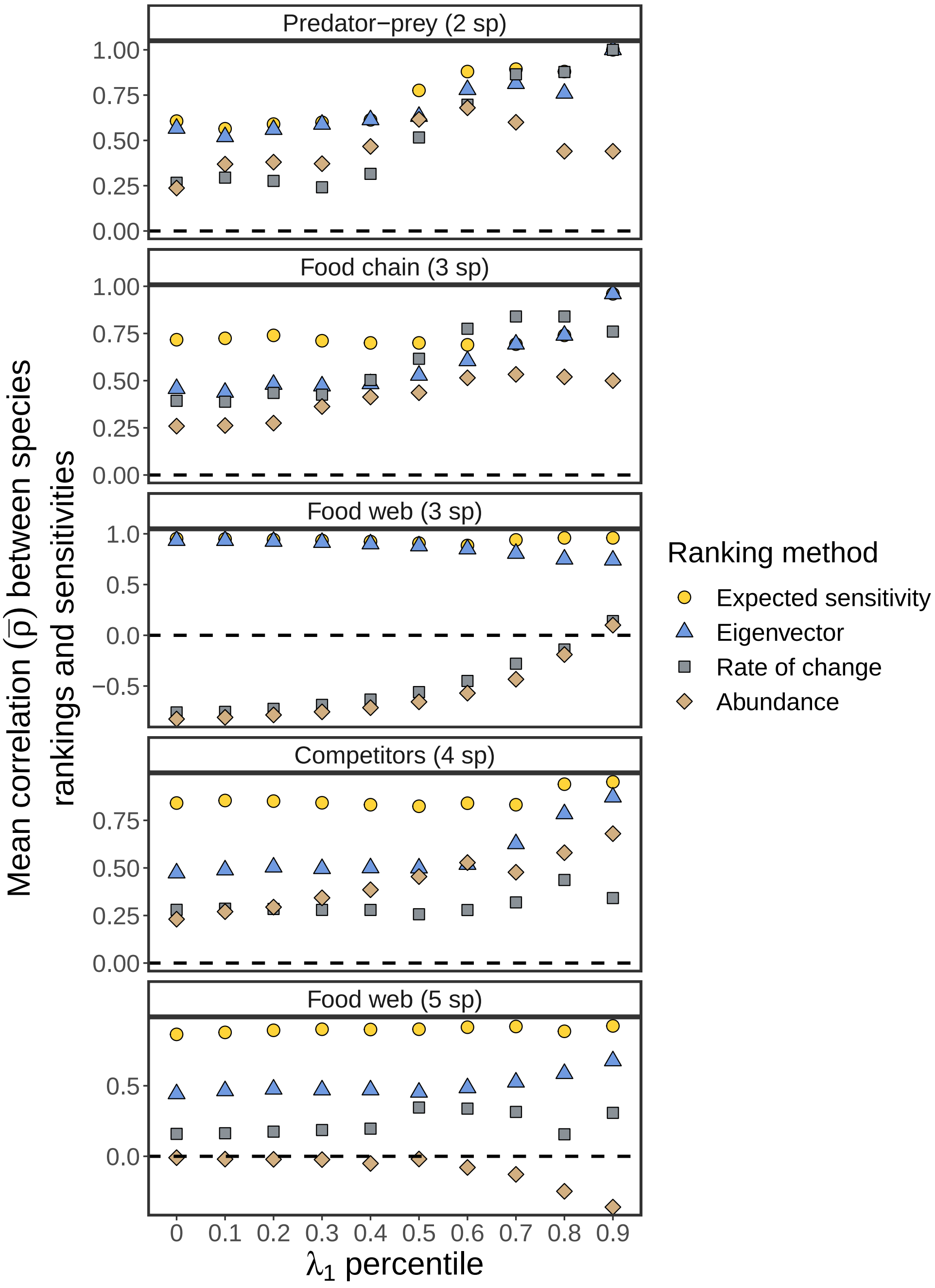


**Figure S7.** Mean Spearman's rank correlation over time ($\bar{\rho}$) between species sensitivities to perturbation ($\langle s_i \rangle$) and four different approaches (expected sensitivity, $\mathbb{E}(s_i)$; eigenvector, $|\mathbf{v}_{1i}|$; rate of change, $\Delta N_i(t)$; and abundance, $-N_i(t)$) as a function of the percentile of $\lambda_1$ used to filter the time series. Each point represents the $\bar{\rho}$ value obtained using a given ranking approach after removing time series points with a $\lambda_1$ value lower than the indicated percentile of the $\lambda_1$ distribution. The figure shows that, for most models, the expected sensitivity and eigenvector rankings (yellow circles and blue triangles) become more accurate (i.e., higher $\bar{\rho}$) when we only use points with a high $\lambda_1$. Note that we compute $\mathbb{E}(s_i)$, $|\mathbf{v}_{1i}|$, and $\lambda_1$ analytically for this figure. Also note that the values of $\bar{\rho}$ for the 0th percentile are exactly the same as the ones shown in Fig. 3A in the main text.

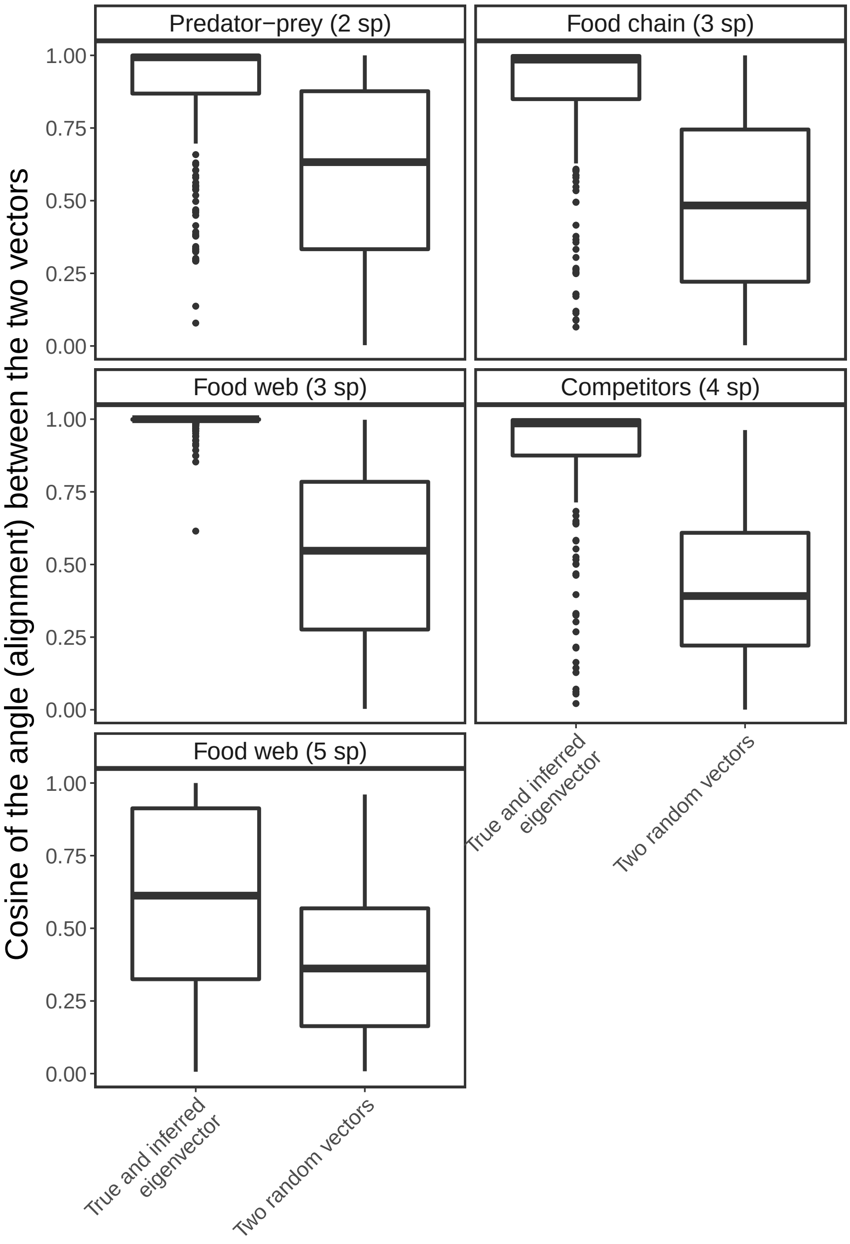


**Figure S8.** Alignments (i.e., absolute value of cosine of the angle) between $\mathbf{v}_1$ inferred with the S-map and $\mathbf{v}_1$ computed from the analytical Jacobian matrix (left boxplots) as well as alignments between two randomly sampled vectors (right boxplots) for each of the five population dynamics models. Each boxplot on the left shows the alignment values computed using the second half of each time series (i.e., last 250 points) for which the S-map was used to infer $\mathbf{v}_1$ (see *Section 5*). Each boxplot on the right shows the alignment values computed using 250 pairs of vectors with random directions. The figure shows that $\mathbf{v}_1$ inferred with the S-map is on average much more aligned with the analytical $\mathbf{v}_1$ than what is expected if their directions are sampled at random.

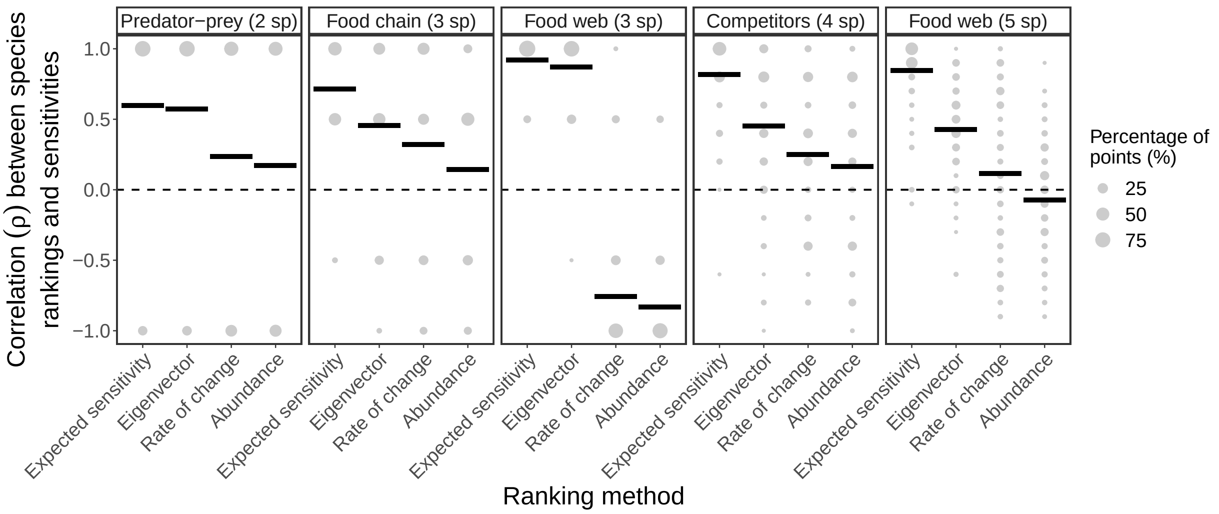


**Figure S9.** Same as Fig. 3A in the main text, but performing uniformly distributed perturbations instead of normally distributed perturbations (see *Section 4*). The figure shows the percentage of points with a given rank correlation value ($\rho$, size of gray points) and the average rank correlation ($\bar{\rho}$, horizontal lines) between species sensitivities to perturbations ($\langle s_i \rangle$) and four different approaches (expected sensitivity, $\mathbb{E}(s_i)$; eigenvector, $|\mathbf{v}_{1i}|$; rate of change, $\Delta N_i(t)$; and abundance, $-N_i(t)$). Note that we compute $\mathbb{E}(s_i)$ and $|\mathbf{v}_{1i}|$ analytically for this figure. For this figure, perturbed abundances ($\tilde{\mathbf{N}}$) are uniformly sampled inside a hypersphere of radius $r$ centered in $\mathbf{N}$, were $r$ corresponds to 15% of the mean standard deviation of species abundances.

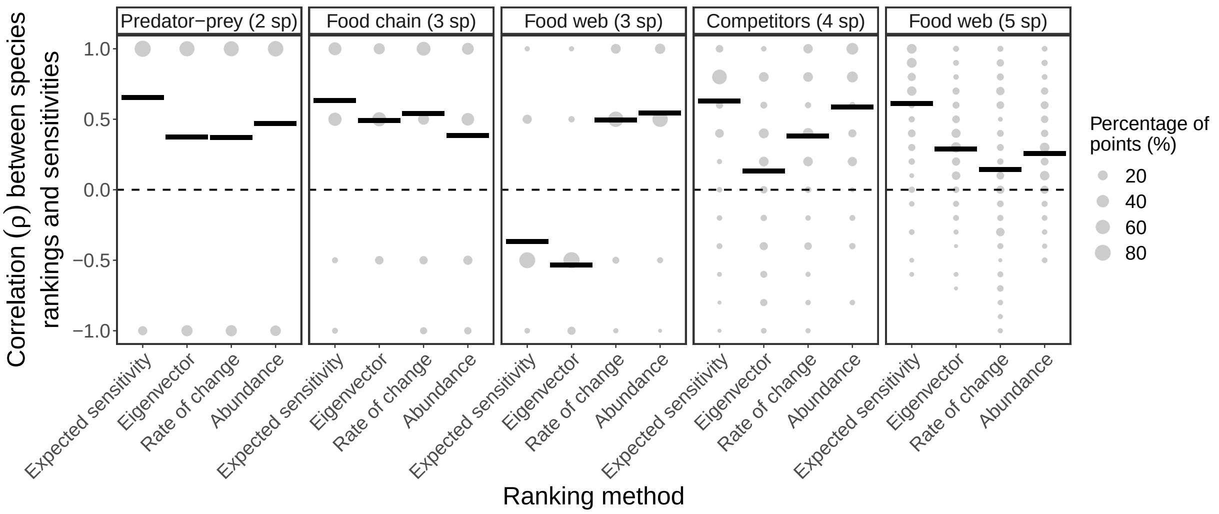


**Figure S10.** Same as Fig. 3A in the main text, but performing normally distributed perturbations with a variance proportional to relative species abundances instead of a fixed variance over time (see *Section 4*). The figure shows the percentage of points with a given rank correlation value ($\rho$, size of gray points) and the average rank correlation ($\bar{\rho}$, horizontal lines) between species sensitivities to perturbations ($\langle s_i \rangle$) and four different approaches (expected sensitivity, $\mathbb{E}(s_i)$; eigenvector, $|\mathbf{v}_{1i}|$; rate of change, $\Delta N_i(t)$; and abundance, $-N_i(t)$). Note that we compute $\mathbb{E}(s_i)$ and $|\mathbf{v}_{1i}|$ analytically for this figure. For this figure, we sample perturbations to $\mathbf{N}(t)$ as: $p_i(t) \sim \mathcal{N}(\mu = 0, \sigma^2 = N_i'(t) r^2)$, where $N_i'(t) = \frac{N_i(t)}{\sum_{i=1}^{S} N_i(t)}$ and were $r$ corresponds to 15% of the mean standard deviation of species abundances.

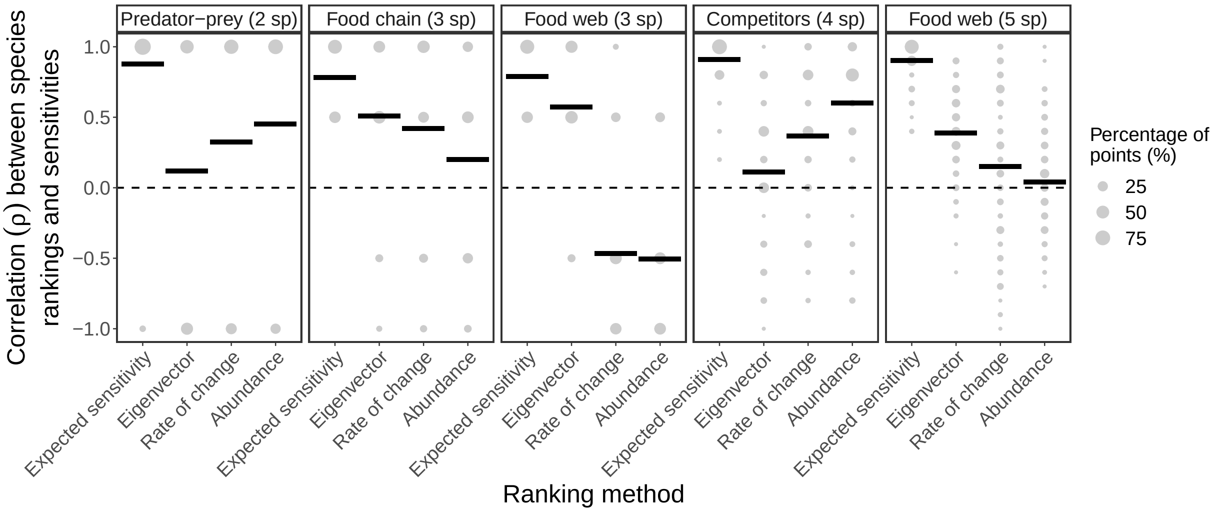


**Figure S11.** Same as Fig. 3A in the main text, but using $k = 1$ as the time step to integrate perturbed and unperturbed abundances instead of $k$ being inversely proportional to the mean absolute abundance percent change (see *Section 4*). The figure shows the percentage of points with a given rank correlation value ($\rho$, size of gray points) and the average rank correlation ($\bar{\rho}$, horizontal lines) between species sensitivities to perturbations ($\langle s_i \rangle$) and four different approaches (expected sensitivity, $\mathbb{E}(s_i)$; eigenvector, $|\mathbf{v}_{1i}|$; rate of change, $\Delta N_i(t)$; and abundance, $-N_i(t)$). Note that we compute $\mathbb{E}(s_i)$ and $|\mathbf{v}_{1i}|$ analytically for this figure. For this figure, we numerically integrate every perturbed ($\tilde{\mathbf{N}}(t)$) and unperturbed abundance ($\mathbf{N}(t)$) for $k = 1$ time step to compute $\langle s_i \rangle$.

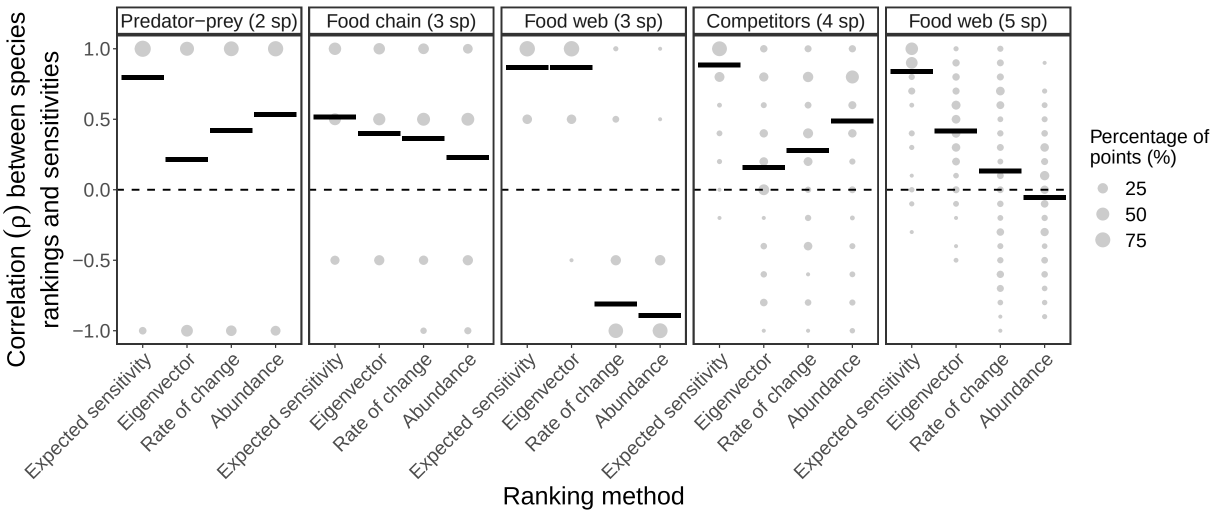


**Figure S12.** Same as Fig. 3A in the main text, but using $k = 3$ as the time step to integrate perturbed and unperturbed abundances instead of $k$ being inversely proportional to the mean absolute abundance percent change (see *Section 4*). The figure shows the percentage of points with a given rank correlation value ($\rho$, size of gray points) and the average rank correlation ($\bar{\rho}$, horizontal lines) between species sensitivities to perturbations ($\langle s_i \rangle$) and four different approaches (expected sensitivity, $\mathbb{E}(s_i)$; eigenvector, $|\mathbf{v}_{1i}|$; rate of change, $\Delta N_i(t)$; and abundance, $-N_i(t)$). Note that we compute $\mathbb{E}(s_i)$ and $|\mathbf{v}_{1i}|$ analytically for this figure. For this figure, we numerically integrate every perturbed ($\tilde{\mathbf{N}}(t)$) and unperturbed abundance ($\mathbf{N}(t)$) for $k = 3$ time steps to compute $\langle s_i \rangle$.

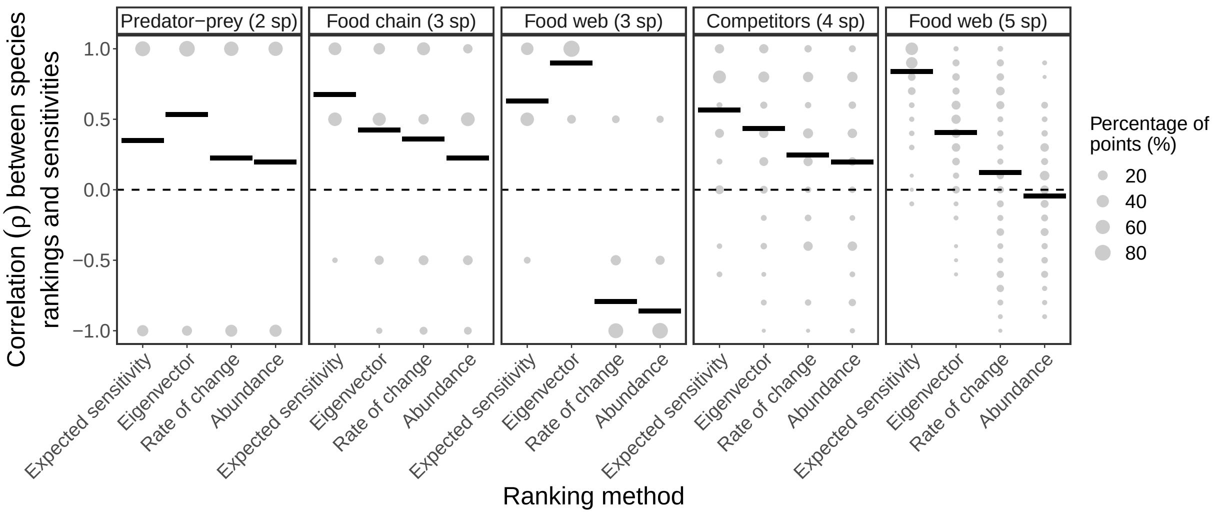


**Figure S13.** Same as Fig. 3A in the main text, but using $k = 1$ as the time step to compute expected sensitivities ($\mathbb{E}(s_i)$) when the true time step used to integrate perturbed and unperturbed abundances is inversely proportional to the mean absolute abundance percent change (see *Section 2*). The figure shows the percentage of points with a given rank correlation value ($\rho$, size of gray points) and the average rank correlation ($\bar{\rho}$, horizontal lines) between species sensitivities to perturbations ($\langle s_i \rangle$) and four different approaches (expected sensitivity, $\mathbb{E}(s_i)$; eigenvector, $|\mathbf{v}_{1i}|$; rate of change, $\Delta N_i(t)$; and abundance, $-N_i(t)$). Note that we compute $\mathbb{E}(s_i)$ and $|\mathbf{v}_{1i}|$ analytically for this figure. For this figure, we numerically integrate every perturbed ($\tilde{\mathbf{N}}(t)$) and unperturbed abundance ($\mathbf{N}(t)$) for a time step $k$ that depends on the local time scale of the dynamics, but always compute $\mathbb{E}(s_i)$ using $k = 1$.

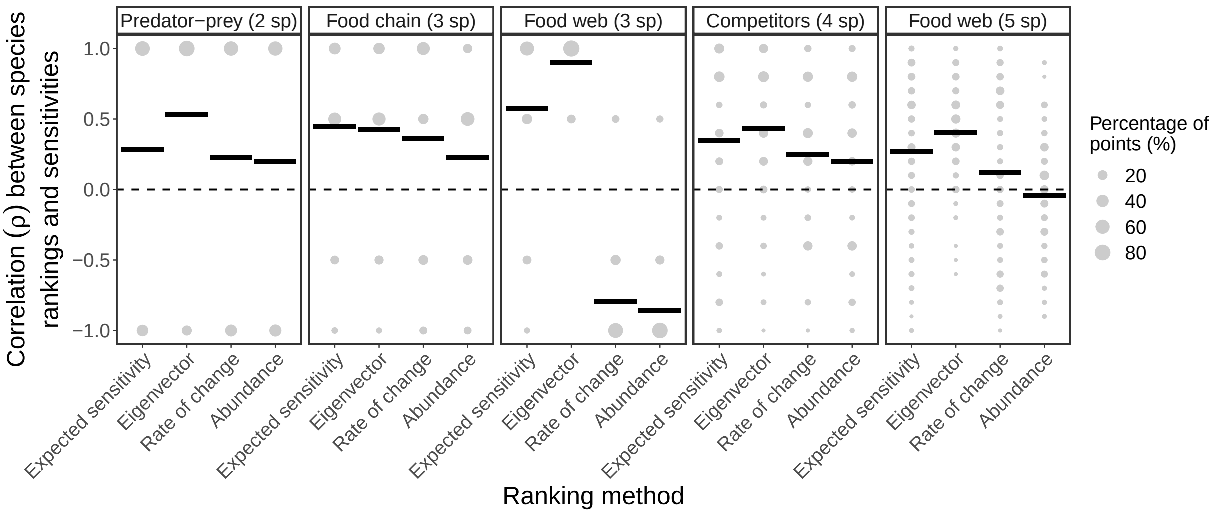


**Figure S14.** Same as Fig. 3A in the main text, but adding a normally distributed noise to $k$ and $\boldsymbol{\Sigma}_t$ at each point in time to compute expected sensitivities ($\mathbb{E}(s_i)$; see *Section 2*). The figure shows the percentage of points with a given rank correlation value ($\rho$, size of gray points) and the average rank correlation ($\bar{\rho}$, horizontal lines) between species sensitivities to perturbations ($\langle s_i \rangle$) and four different approaches (expected sensitivity, $\mathbb{E}(s_i)$; eigenvector, $|\mathbf{v}_{1i}|$; rate of change, $\Delta N_i(t)$; and abundance, $-N_i(t)$). Note that we compute $\mathbb{E}(s_i)$ and $|\mathbf{v}_{1i}|$ analytically for this figure. For this figure, we perform the same perturbation analyses as described for Fig. 3 (see *Section 4*), but add 100% of a normally distributed noise to the true value of $k$ and to $\boldsymbol{\Sigma}_t = \mathbf{I}$ before computing $\mathbb{E}(s_i)$.

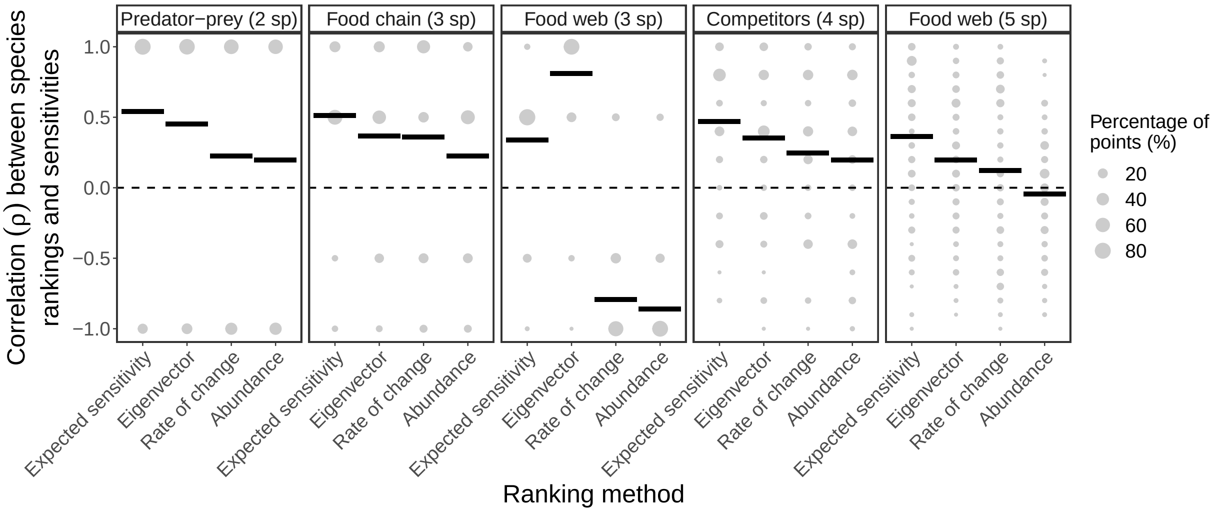


**Figure S15.** Same as Fig. 3B in the main text, but normalizing the abundances of each species $i$ ($N_i(t)$) in the training set to mean zero and unit standard deviation before performing the S-map. The figure shows the percentage of points with a given rank correlation value ($\rho$, size of gray points) and the average rank correlation ($\bar{\rho}$, horizontal lines) between species sensitivities to perturbations ($\langle s_i \rangle$) and four different approaches (expected sensitivity, $\mathbb{E}(s_i)$; eigenvector, $|\mathbf{v}_{1i}|$; rate of change, $\Delta N_i(t)$; and abundance, $-N_i(t)$). Note that we infer the Jacobian matrix with the S-map using a moving training set in order to compute $\mathbb{E}(s_i)$ and $|\mathbf{v}_{1i}|$ for this figure.

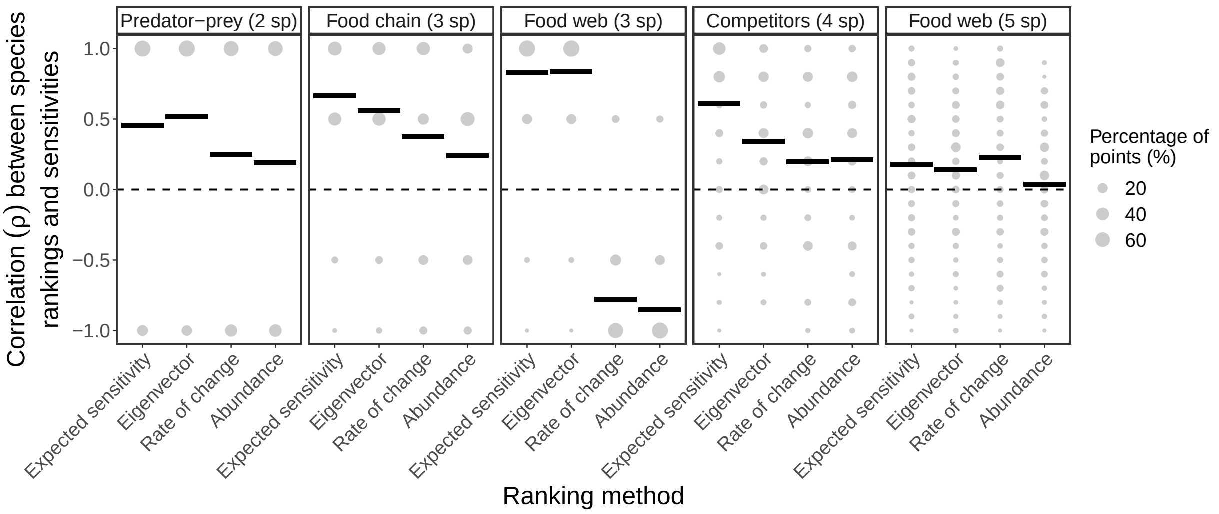


**Figure S16.** Same as Fig. 3B in the main text, but using a shorter training set with 100 instead of 250 points to perform the S-map (see *Section 6*). The figure shows the percentage of points with a given rank correlation value ($\rho$, size of gray points) and the average rank correlation ($\bar{\rho}$, horizontal lines) between species sensitivities to perturbations ($\langle s_i \rangle$) and four different approaches (expected sensitivity, $\mathbb{E}(s_i)$; eigenvector, $|\mathbf{v}_{1i}|$; rate of change, $\Delta N_i(t)$; and abundance, $-N_i(t)$). Note that we infer the Jacobian matrix with the S-map using a moving training set in order to compute $\mathbb{E}(s_i)$ and $|\mathbf{v}_{1i}|$ for this figure.

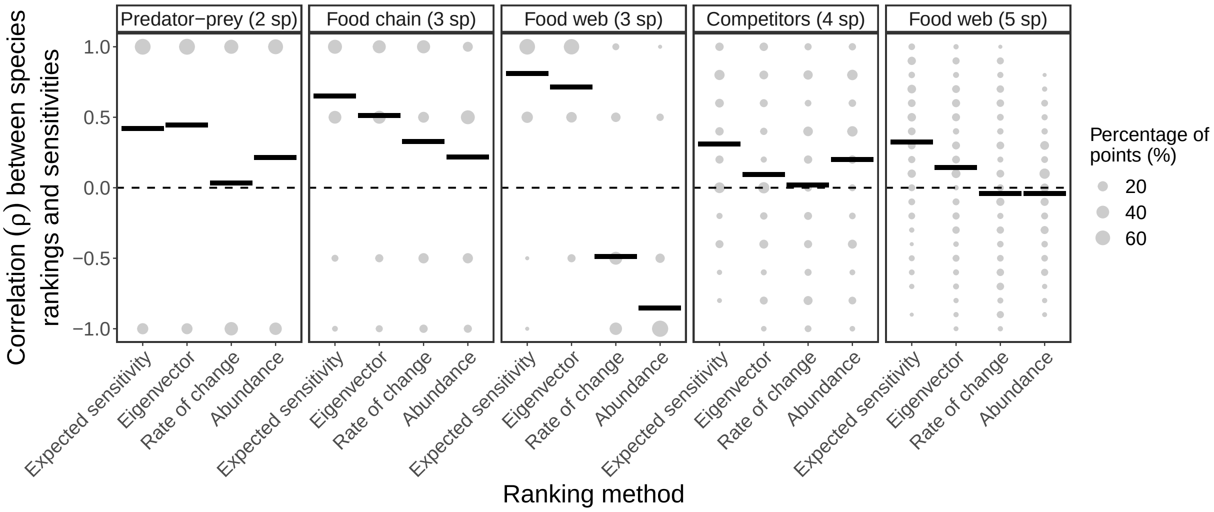


**Figure S17.** Same as Fig. 3B in the main text, but adding 10% of observational noise to the training set before performing the S-map (see *Section 6*). The figure shows the percentage of points with a given rank correlation value ($\rho$, size of gray points) and the average rank correlation ($\bar{\rho}$, horizontal lines) between species sensitivities to perturbations ($\langle s_i \rangle$) and four different approaches (expected sensitivity, $\mathbb{E}(s_i)$; eigenvector, $|\mathbf{v}_{1i}|$; rate of change, $\Delta N_i(t)$; and abundance, $-N_i(t)$). Note that we infer the Jacobian matrix with the S-map using a moving training set in order to compute $\mathbb{E}(s_i)$ and $|\mathbf{v}_{1i}|$ for this figure.

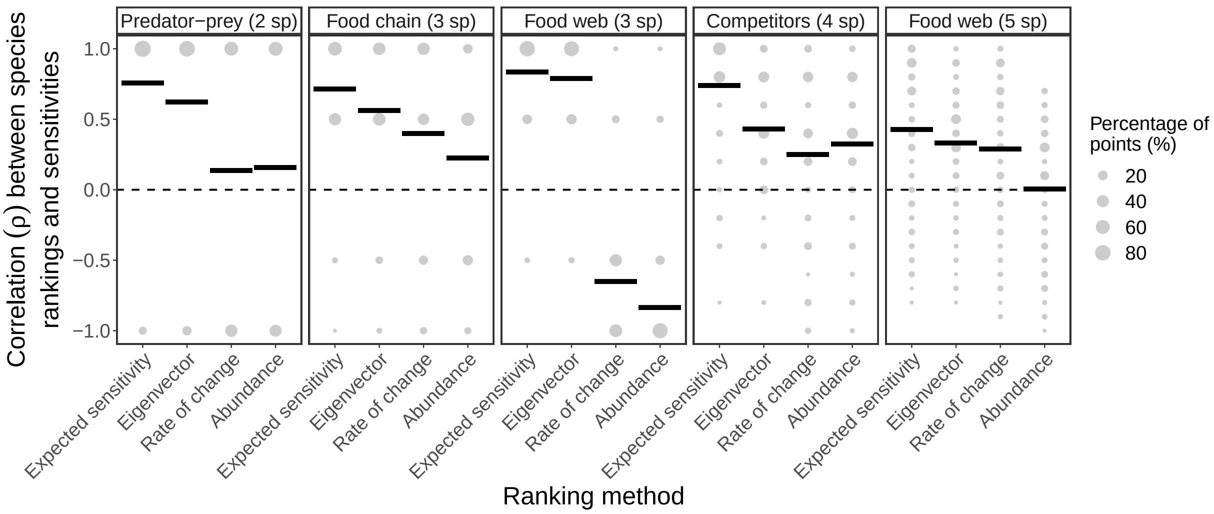


**Figure S18.** Same as Fig. 3B in the main text, but generating each synthetic time series with the population dynamics model containing stochasticity (i.e., process noise; see *Section 6*). The figure shows the percentage of points with a given rank correlation value ($\rho$, size of gray points) and the average rank correlation ($\bar{\rho}$, horizontal lines) between species sensitivities to perturbations ($\langle s_i \rangle$) and four different approaches (expected sensitivity, $\mathbb{E}(s_i)$; eigenvector, $|\mathbf{v}_{1i}|$; rate of change, $\Delta N_i(t)$; and abundance, $-N_i(t)$). Note that we infer the Jacobian matrix with the S-map using a moving training set in order to compute $\mathbb{E}(s_i)$ and $|\mathbf{v}_{1i}|$ for this figure.

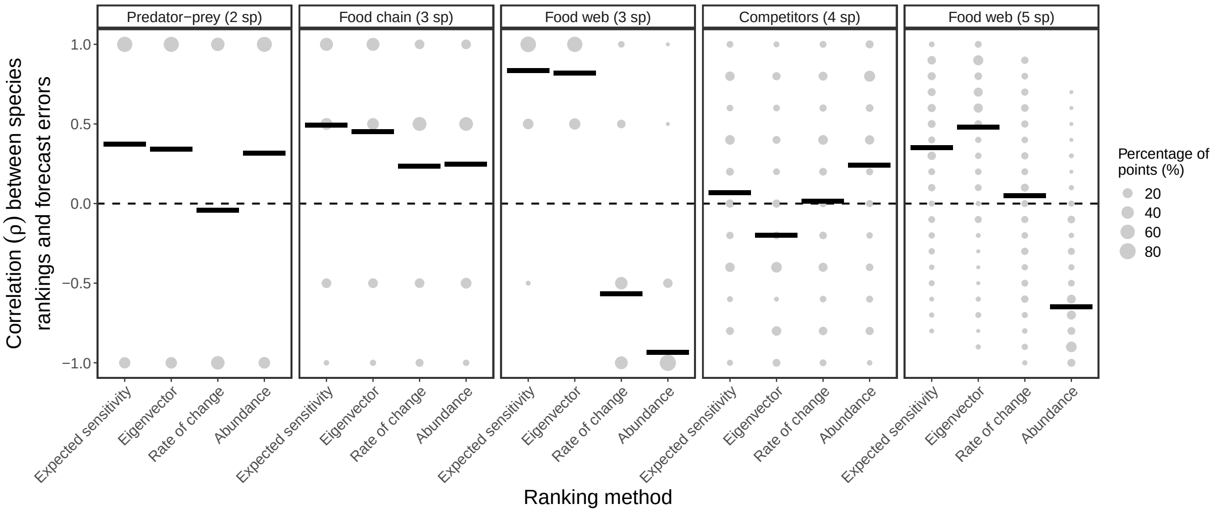


**Figure S19.** Similar to Fig. 3B in the main text, but here we compute the Spearman's rank correlation ($\rho$) between species average forecast errors under perturbations ($\bar{\epsilon}_i$; see *Section 8*) and the four ranking approaches (expected sensitivity, $\mathbb{E}(s_i)$; eigenvector, $|\mathbf{v}_{1i}|$; rate of change, $\Delta N_i(t)$; and abundance, $-N_i(t)$). The figure shows the percentage of points with a given $\rho$ value (size of gray points) and the average rank correlation ($\bar{\rho}$, horizontal lines). Note that we infer the Jacobian matrix with the S-map using a moving training set in order to compute $\mathbb{E}(s_i)$ and $|\mathbf{v}_{1i}|$ for this figure. This figure illustrates our hypothesis that species that are more sensitive to perturbations (i.e., high $\mathbb{E}(s_i)$ or $|\mathbf{v}_{1i}|$) tend to be harder to forecast under perturbations (i.e., high $\bar{\epsilon}_i$).

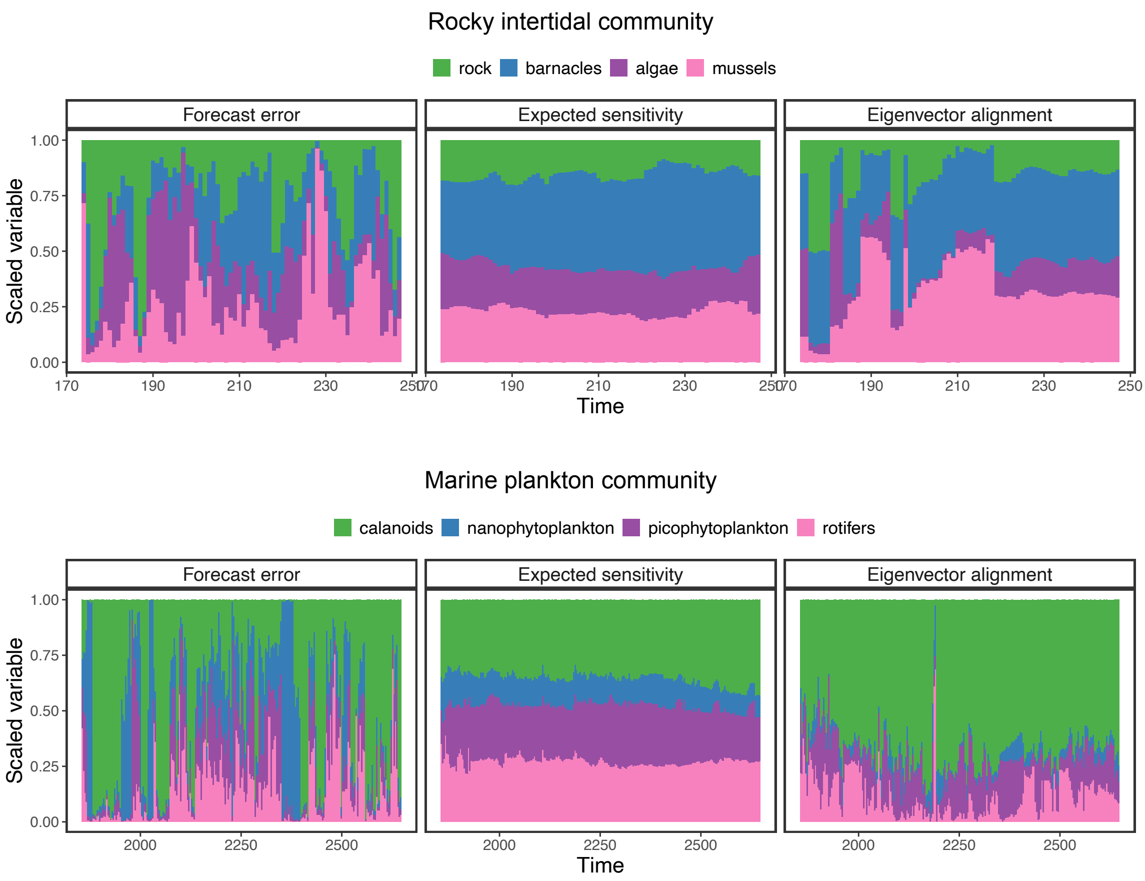


**Figure S20.** Species standardized forecast root-mean-square error computed from our forecast analyses ($\epsilon_i$; first column; see *Section 7*) as well as expected sensitivities ($\mathbb{E}(s_i)$; second column) and eigenvector alignments ($|\mathbf{v}_{1i}|$; third column) inferred from each empirical time series (different rows) with the S-map over time. A bar in one of the plots shows the values of the corresponding variable (i.e., $\epsilon_i$, $\mathbb{E}(s_i)$, or $|\mathbf{v}_{1i}|$) across species. Note that variables are rescaled to sum 1 across species to improve visualization but that this procedure does not change the rankings. These results correspond to our main set of analyses with empirical time series shown in the main text (Fig. 4).

(a)

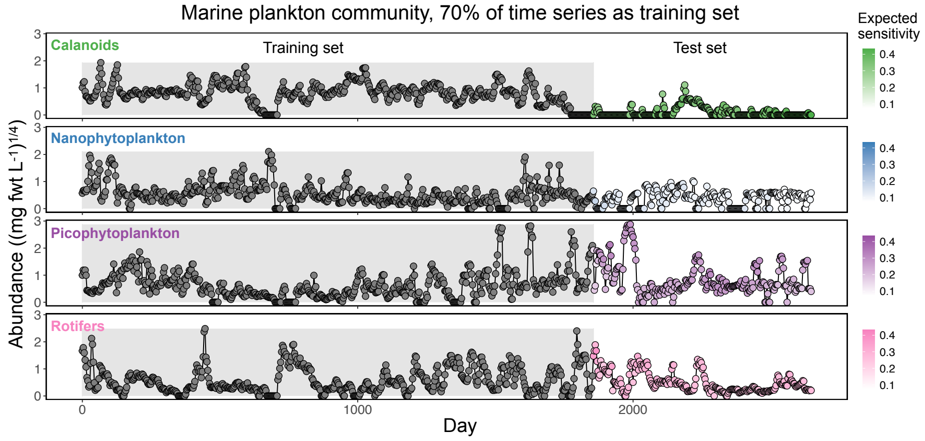


(b)

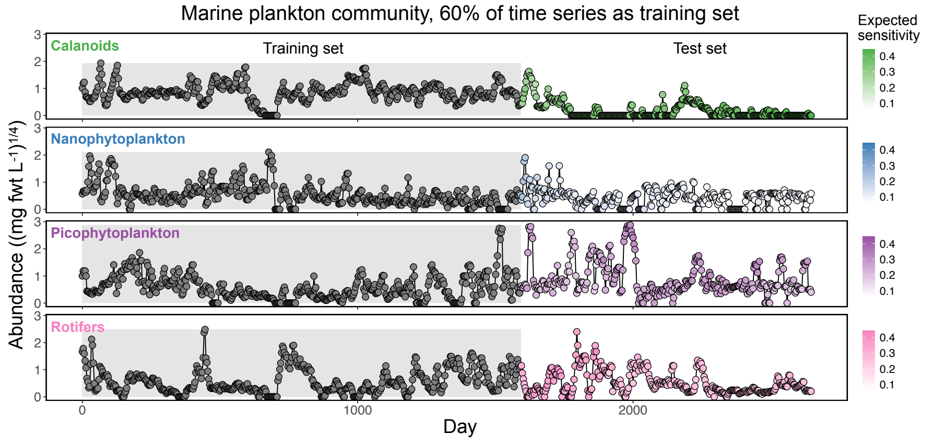


(c)

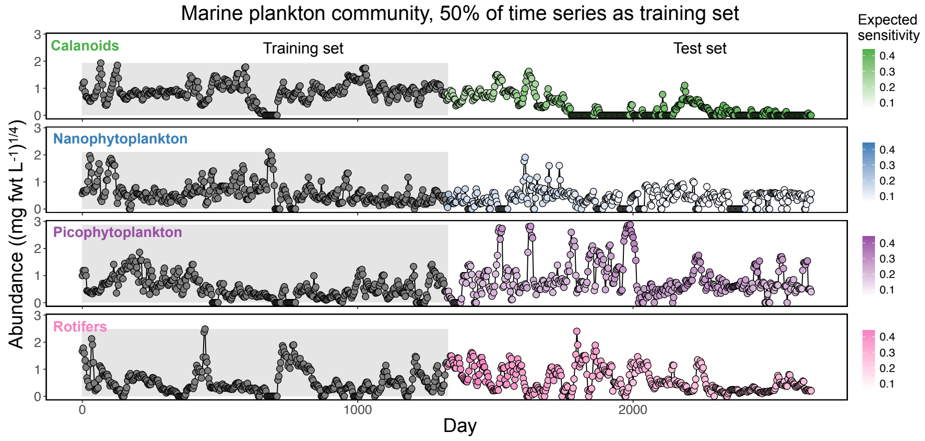


**Figure S21.** Same as Fig. 4A in the main text but for the empirical time series of marine plankton species (Benincà *et al.*, 2009) (see *Section 7*). Each panel shows the time series of the abundance of a given species with points colored according to their expected sensitivity value ($\mathbb{E}(s_i)$). We infer $\mathbb{E}(s_i)$ at the last point in the training set with the S-map trained on a moving training set (gray region) containing (a) 70%, (b) 60%, or (c) 50% of the whole time series. In general, calanoids are the most sensitive species followed by rotifers or picocyanobacteria depending on the point in time.

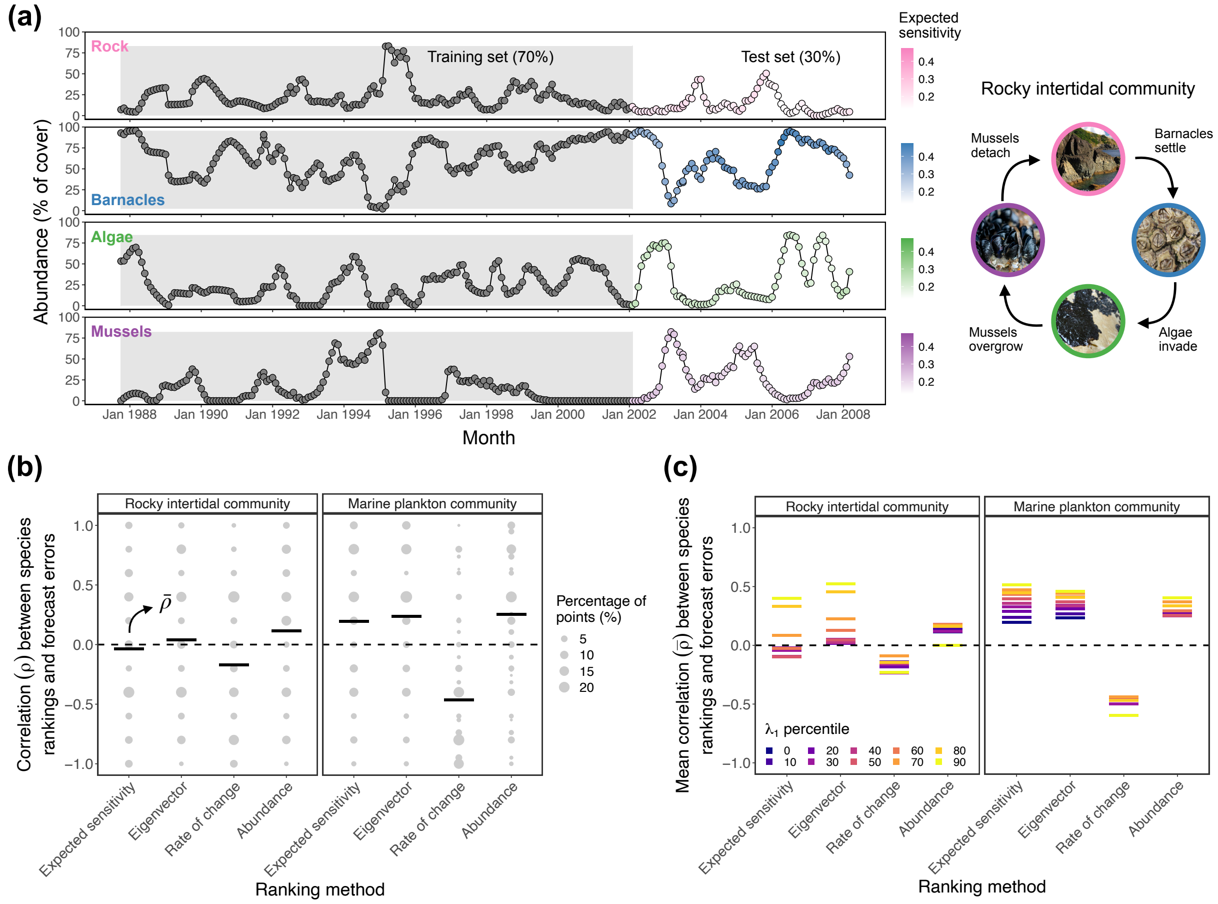


**Figure S22.** Same as Fig. 4 in the main text but using $\tau = 2$ steps ahead to forecast species abundances and compute forecast errors ($\epsilon_i$) instead of $\tau = 3$ (see *Section 7*). Note that here we use $k = 2$ instead of $k = 3$ to compute expected sensitivities ($\mathbb{E}(s_i)$). (a) Time series of a rocky intertidal community containing four species with point color depicting their expected sensitivity value. (b) Rank correlation ($\rho$) between $\epsilon_i$ and four different approaches (expected sensitivity, $\mathbb{E}(s_i)$; eigenvector, $|\mathbf{v}_{1i}|$; rate of change, $\Delta N_i(t)$; and abundance, $-N_i(t)$). Each panel shows the percentage of points with a given $\rho$ value (size of gray points) and the average of these values across the test set ($\bar{\rho}$, horizontal lines) for a given empirical time series. (c) Average correlation ($\bar{\rho}$) between $\epsilon_i$ and the different ranking approaches computed for points in the test set that have a $\lambda_1$ value higher than a given percentile of the $\lambda_1$ distribution.

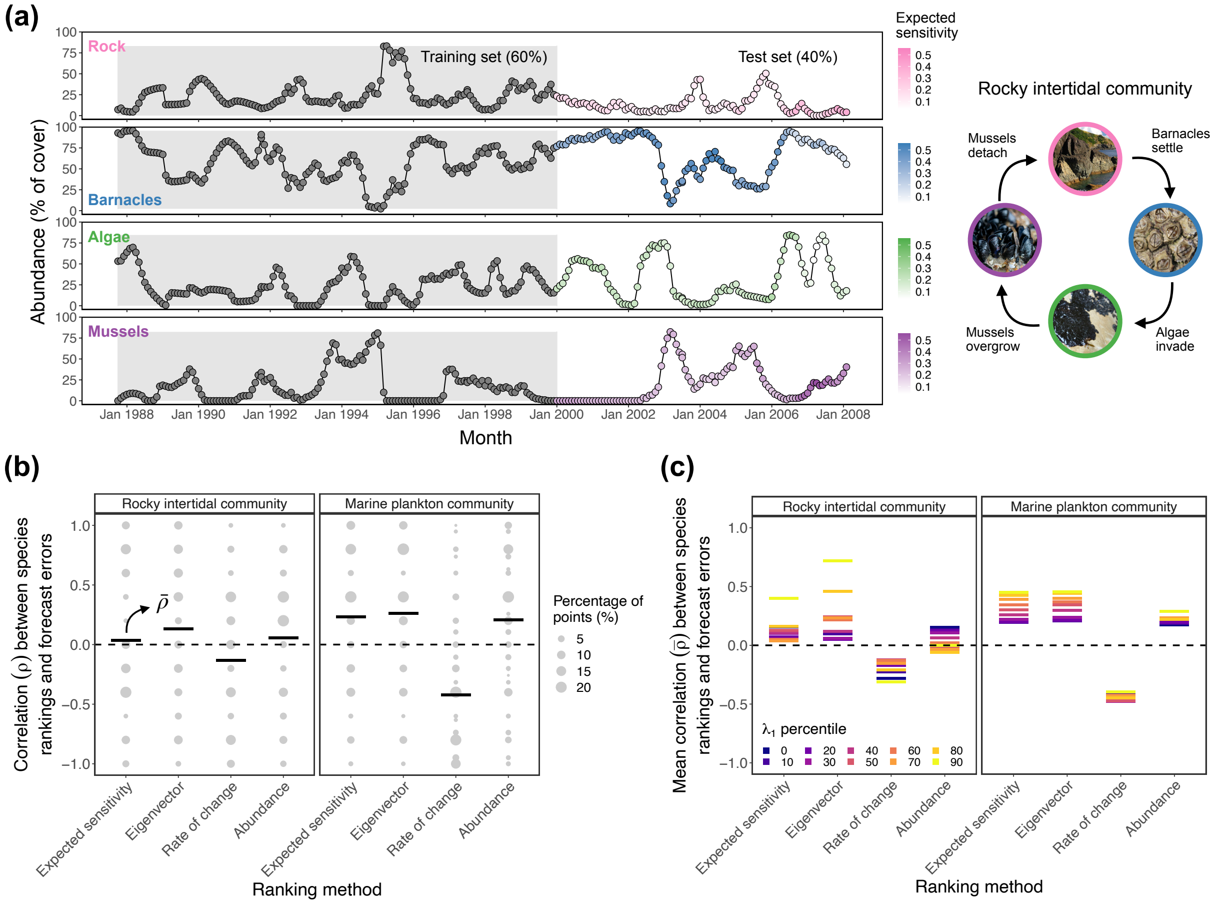


**Figure S23.** Same as Fig. 4 in the main text but using 60% instead of 70% of the each empirical time series as the moving training set (gray region in (a); see *Section 7*). (a) Time series of a rocky intertidal community containing four species with point color depicting their expected sensitivity value ($\mathbb{E}(s_i)$). (b) Rank correlation ($\rho$) between $\epsilon_i$ and four different approaches (expected sensitivity, $\mathbb{E}(s_i)$; eigenvector, $|\mathbf{v}_{1i}|$; rate of change, $\Delta N_i(t)$; and abundance, $-N_i(t)$). Each panel shows the percentage of points with a given $\rho$ value (size of gray points) and the average of these values across the test set ($\bar{\rho}$, horizontal lines) for a given empirical time series. (c) Average correlation ($\bar{\rho}$) between $\epsilon_i$ and the different ranking approaches computed for points in the test set that have a $\lambda_1$ value higher than a given percentile of the $\lambda_1$ distribution.

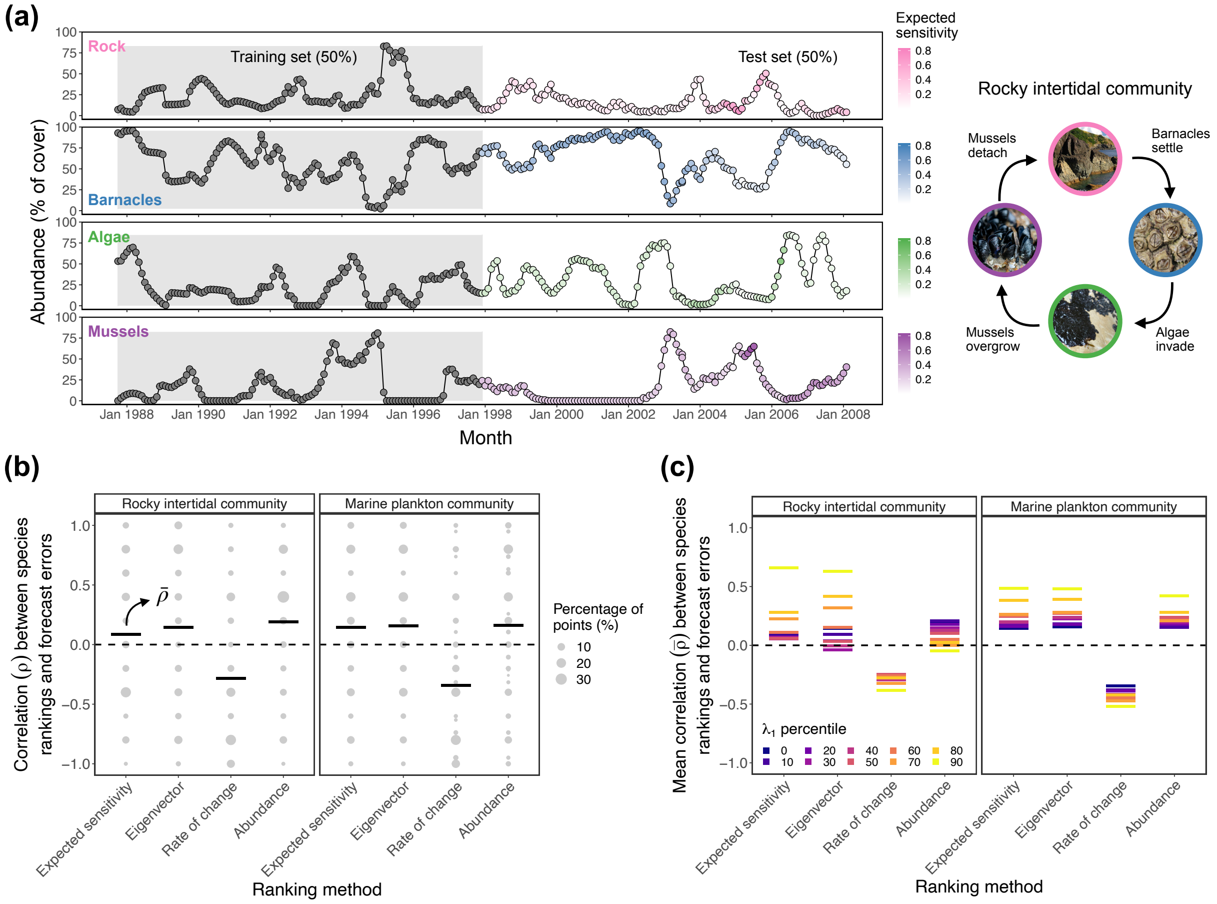


**Figure S24.** Same as Fig. 4 in the main text but using 50% instead of 70% of the each empirical time series as the moving training set (gray region in (a); see *Section 7*). (a) Time series of a rocky intertidal community containing four species with point color depicting their expected sensitivity value ($\mathbb{E}(s_i)$). (b) Rank correlation ($\rho$) between $\epsilon_i$ and four different approaches (expected sensitivity, $\mathbb{E}(s_i)$; eigenvector, $|\mathbf{v}_{1i}|$; rate of change, $\Delta N_i(t)$; and abundance, $-N_i(t)$). Each panel shows the percentage of points with a given $\rho$ value (size of gray points) and the average of these values across the test set ($\bar{\rho}$, horizontal lines) for a given empirical time series. (c) Average correlation ($\bar{\rho}$) between $\epsilon_i$ and the different ranking approaches computed for points in the test set that have a $\lambda_1$ value higher than a given percentile of the $\lambda_1$ distribution.

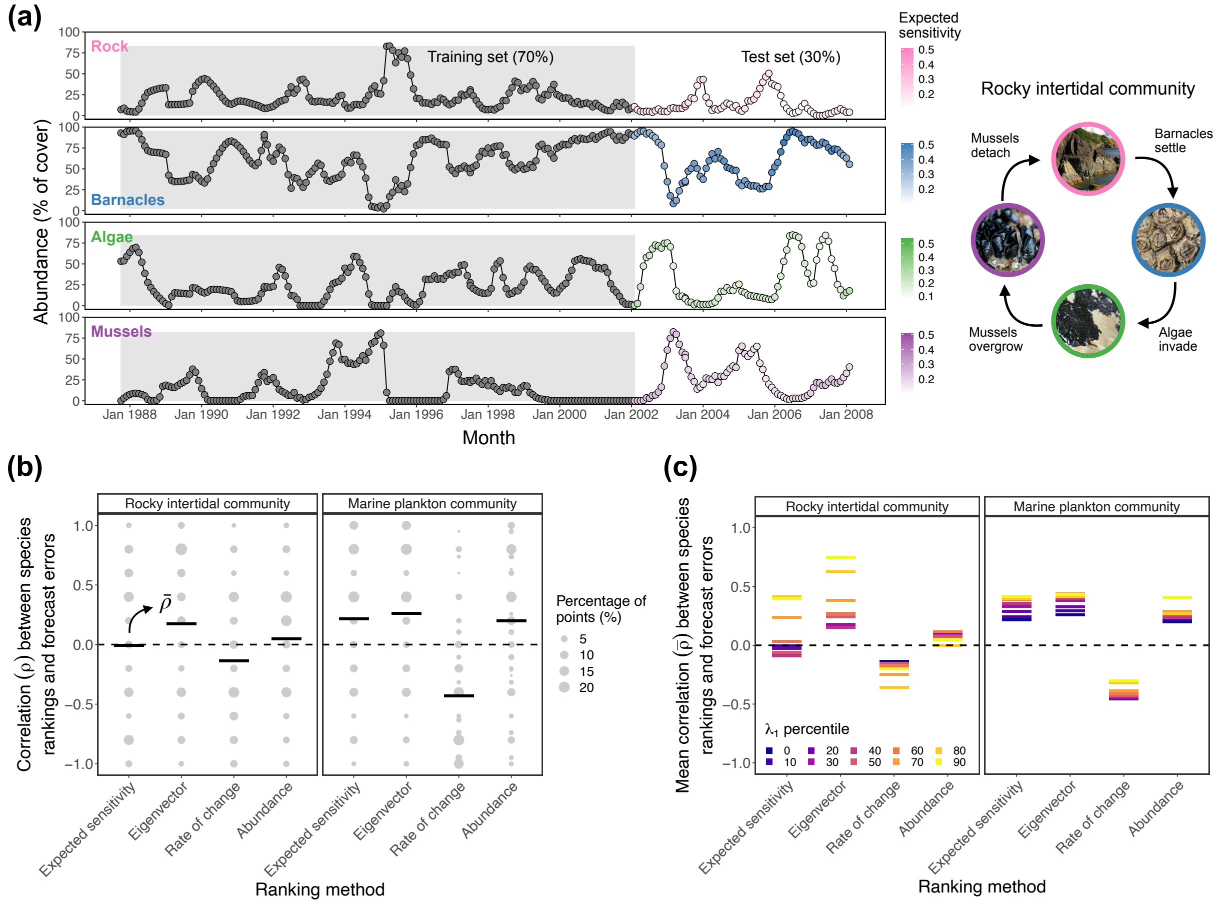


**Figure S25.** Same as Fig. 4 in the main text but normalizing the abundances of each species $i$ ($N_i(t)$) in the training set to mean zero and unit standard deviation before performing the S-map (see *Section 7*). Note that we always normalize abundances before the forecast analyses (i.e., LSTM neural network). (a) Time series of a rocky intertidal community containing four species with point color depicting their expected sensitivity value ($\mathbb{E}(s_i)$). (b) Rank correlation ($\rho$) between $\epsilon_i$ and four different approaches (expected sensitivity, $\mathbb{E}(s_i)$; eigenvector, $|\mathbf{v}_{1i}|$; rate of change, $\Delta N_i(t)$; and abundance, $-N_i(t)$). Each panel shows the percentage of points with a given $\rho$ value (size of gray points) and the average of these values across the test set ($\bar{\rho}$, horizontal lines) for a given empirical time series. (c) Average correlation ($\bar{\rho}$) between $\epsilon_i$ and the different ranking approaches computed for points in the test set that have a $\lambda_1$ value higher than a given percentile of the $\lambda_1$ distribution.
